# An Innate Neuroprotective Reflex Activates NRF2 via Phosphorylation of p62 at Ser351 to Mitigate Traumatic Brain Injury

**DOI:** 10.64898/2026.11.03.763957

**Authors:** Chunyan Li, Keren Powell, Steven Wadolowski, David LeDoux, Amir R. Dehdashti, Jamie S. Ullman, Max Brenner, Christopher LeDoux, Kaitlyn Rams, Daniel Sciubba, Ping Wang

## Abstract

**Rationale:** The oxidative crisis in traumatic brain injury (TBI) has remained therapeutically intractable to direct antioxidant therapies. Activating nuclear factor erythroid 2-related factor 2 (NRF2) offers a promising strategy, but its translation is hindered by a key question: whether TBI’s depleted antioxidant reserves can accommodate canonical electrophilic activation. The diving reflex (DR) is a potential non-electrophilic alternative, but its efficacy in TBI is unknown and its natural induction is clinically impractical. This study tests whether artificial DR, induced by trigeminal nerve stimulation (TNS), can safely engage this non-electrophilic NRF2 pathway and exert a protective role in the acutely injured brain.

**Methods:** In a rat model of severe TBI, we compared TNS-induced DR, natural DR, and exogenous calcitonin gene-related peptide (CGRP) against the electrophilic NRF2 activator dimethyl fumarate (DMF). To define the mechanism of action, we quantified p62 phosphorylation at Ser351 (NRF2-activating) versus Ser403 (autophagy-related), alongside redox homeostasis, lesion volume, and multi-domain functional outcomes.

**Results:** TNS-induced DR fully recapitulated the neuroprotection of natural DR, an effect mimicked by exogenous CGRP, and prevented by CGRP inhibition, confirming a shared, neuropeptide-driven mechanism. This non-electrophilic pathway operated by selectively phosphorylating p62 at Ser351, which preserved glutathione reserves, halted lipid peroxidation, and reduced chronic lesion volume. In contrast, while DMF activated NRF2, its electrophilic mechanism was associated with a further reduction in the brain’s already compromised glutathione pools and increased lipid peroxidation. This mechanistic difference was functionally significant, as only the non-electrophilic approaches delivered multi-domain recovery.

**Conclusion:** The TNS-evoked artificial DR is a mechanistically tailored strategy for activating NRF2 in acute TBI. By engaging the endogenous CGRP-p62-KEAP1 pathway, it circumvents the fundamental limitations of both passively consumed direct antioxidants and redox-costly electrophilic activators. Our findings establish a clear translational roadmap for harnessing an innate reflex as a clinically deployable therapy for hyper-oxidative states.

## Introduction

Traumatic brain injury (TBI) affects approximately 18% of US adults and 20-40% of military personnel, representing a major public health crisis with no approved neuroprotective treatments [1–3]. Central to TBI pathophysiology is oxidative stress, a catastrophic surge of reactive species that overwhelms endogenous antioxidant defenses within seconds of injury, initiating immediate neuronal damage [4,5]. Critically, this establishes a self-perpetuating cycle wherein damaged neurons become autonomous reactive species generators, driving progressive neurodegeneration for months post-injury [6]. Despite oxidative stress being the primary therapeutic target in >10% of registered TBI trials on ClinicalTrials.gov, antioxidant interventions have not demonstrated overall clinical efficacy in TBI, with no intervention has advancing beyond Phase III [4,5]. This universal failure reflects fundamental limitations: single agents cannot sufficiently neutralize TBI’s heterogeneous reactive species cocktail, while combination therapies paradoxically demonstrate no superiority over monotherapies [7]. These failures underscore a critical insight, rather than attempting to scavenge individual reactive species after their formation, effective therapy is better directed toward preemptively enhancing the brain’s entire antioxidant network. Nuclear factor erythroid 2–related factor 2 (NRF2) may represent this paradigmatic shift [4,8]. As a master regulator of cellular antioxidant responses, NRF2 orchestrates expression of >250 cytoprotective genes containing antioxidant response elements, simultaneously upregulating multiple defense systems. This comprehensive, multi-modal activation may transform the therapeutic approach from reactive scavenging to proactive cellular protection, matching TBI’s complex oxidative pathology.

Current NRF2-based therapies have largely relied on electrophilic activators, compounds that induce mild oxidative stress to alkylate KEAP1 cysteines and release NRF2 [9,10]. In the hyperacute phase of TBI, however, this approach has the capacity to exacerbate the surge of reactive species and collapse of glutathione stores which occurs within minutes of impact [11]. Preclinical data show that, despite significant activation of NRF2, electrophilic activators have been demonstrated to further deplete glutathione, amplify lipid peroxidation, and worsen both tissue damage and functional outcomes [9,12]. While these paradoxical observations may be due to specific dosing or compound constraints, they also suggest that NRF2 activation in TBI may be more effective with a redox-neutral mechanism rather than exacerbating an already exhausted antioxidant pool. Non-electrophilic activation via the p62–KEAP1 axis illustrates just this, achieving comparable NRF2 activation potency while maintaining a redox-neutral state [10]. Phosphorylation of p62 at Ser351 directs KEAP1 into selective autophagic degradation, thereby liberating bound NRF2 without consuming thiol reserves, while also preventing the binding of future NRF2 through reduction of KEAP1 reserves. Critically, TBI itself triggers a hyperacute suppression of p62 [13–15], creating a molecular bottleneck that prevents the very pathway needed to defend against oxidative damage. Restoring p62-mediated KEAP1 turnover thus has the potential to provide a mechanistically significant way to revive NRF2’s full antioxidant program exactly when and where it matters most, without exacerbating oxidative stress.

The diving reflex (DR), or oxygen conserving reflex, an evolutionarily conserved autonomic response triggered by trigeminal afferents, may offer a unique solution to a central therapeutic paradox in TBI: how to bolster antioxidant defenses without imposing additional oxidative stress. Our prior work in a voluntary-diving rat model has elucidated the unique, non-electrophilic mechanism driving antioxidant-mediated neuroprotection in natural DR [16]. We discovered that within minutes of breath-hold diving, trigeminal-derived calcitonin gene-related peptide (CGRP) drives the phosphorylation of p62 at Ser351. This crucial modification specifically directs the autophagic degradation of KEAP1 and subsequent liberation of NRF2 for rapid nuclear translocation, all achieved while increasing, rather than depleting, glutathione reserves and retaining cellular redox balance. The power of this endogenous pathway is significant; it achieves rapid pharmacological-level NRF2 activation, and when applied chronically in a vascular dementia model, it prevents hippocampal neuronal NRF2 depletion and preserves cognitive performance [16,17]. While compelling, two critical hurdles have thus far prevented its translation to TBI: the clinical impracticality of breath-hold diving for DR activation in incapacitated patients, and the unknown efficacy of this mechanism in the profoundly pro-oxidant TBI milieu. To overcome the first barrier, we have previously developed an electrical trigeminal nerve stimulation (TNS) method that faithfully replicates DR’s distinct hemodynamic signatures [18], providing a clinically viable surrogate.

This study was therefore designed to address the pivotal remaining question: can a clinically viable TNS surrogate replicate the natural DR to confer strong neuroprotection in the acutely injured brain? We hypothesized that TNS-induced DR, by engaging the non-electrophilic CGRP-p62-KEAP1 pathway, would provide superior, redox-neutral neuroprotection compared to canonical electrophilic activators. To rigorously test this, we pursued a multi-pronged investigation. First, we established the mechanistic fidelity of TNS-induced artificial DR to natural DR, specifically by demonstrating its capacity to drive selective phosphorylation of p62 at Ser351 (for NRF2 activation) while bypassing the general autophagy marker Ser403. Second, we directly contrasted the neuroprotective efficacy of our non-electrophilic strategies against the canonical electrophilic activator, DMF, to determine their divergent performance in TBI’s oxidatively compromised environment. Third, we confirmed the CGRP-dependency of the entire cascade through pharmacological blockade and mimicry. Finally, we evaluated the long-term structural and multi-domain functional outcomes of our TNS strategy to assess its clinical utility in reducing chronic morbidity. By systematically validating this endogenous pathway from its physiological trigger to its precise molecular execution and long-term functional impact, this work aims to establish a mechanistically sound and redox-neutral strategy to address TBI’s devastating oxidative crisis after decades of translational failure.

## Materials and methods

### Experimental animals and design

All procedures were approved by the Institutional Animal Care and Use Committee of the Feinstein Institutes for Medical Research and conducted in accordance with the NIH Guide for the Care and Use of Laboratory Animals. Male Sprague-Dawley rats were obtained at 5 weeks old and weighing 100–125g at the beginning of the experiment, were used, and all rats were between 350–450g at the time of collection (Charles River Laboratories, Kingston, NY). Animals were housed in a temperature-controlled room (12-h light/dark cycle), in cages lined with Enrich-o’Cobbs bedding (The Anderson, Inc, Maumee, OH) and were given access to food and water *ad libitum*. A total of 240 rats were randomized across three experimental protocols (**Figure 1A**). Pharmacological CGRP inhibition was achieved using two distinct agents tailored to the experimental context. In Experiment 1, which involved conscious animal behavior, the long-acting monoclonal antibody Fremanezumab (30 mg/kg, intranasal; Ajovy, Teva Pharmaceuticals, Parsippany, NJ, USA) was administered. For Experiment 3, which required targeted, short-term CGRP antagonism exclusively during a 30-minute TNS session, the CGRP receptor antagonist, CGRP_8-37_ (200 µg/kg, IV infusion; Tocris Biosciences, Bristol, UK), was utilized.

**Figure 1.**
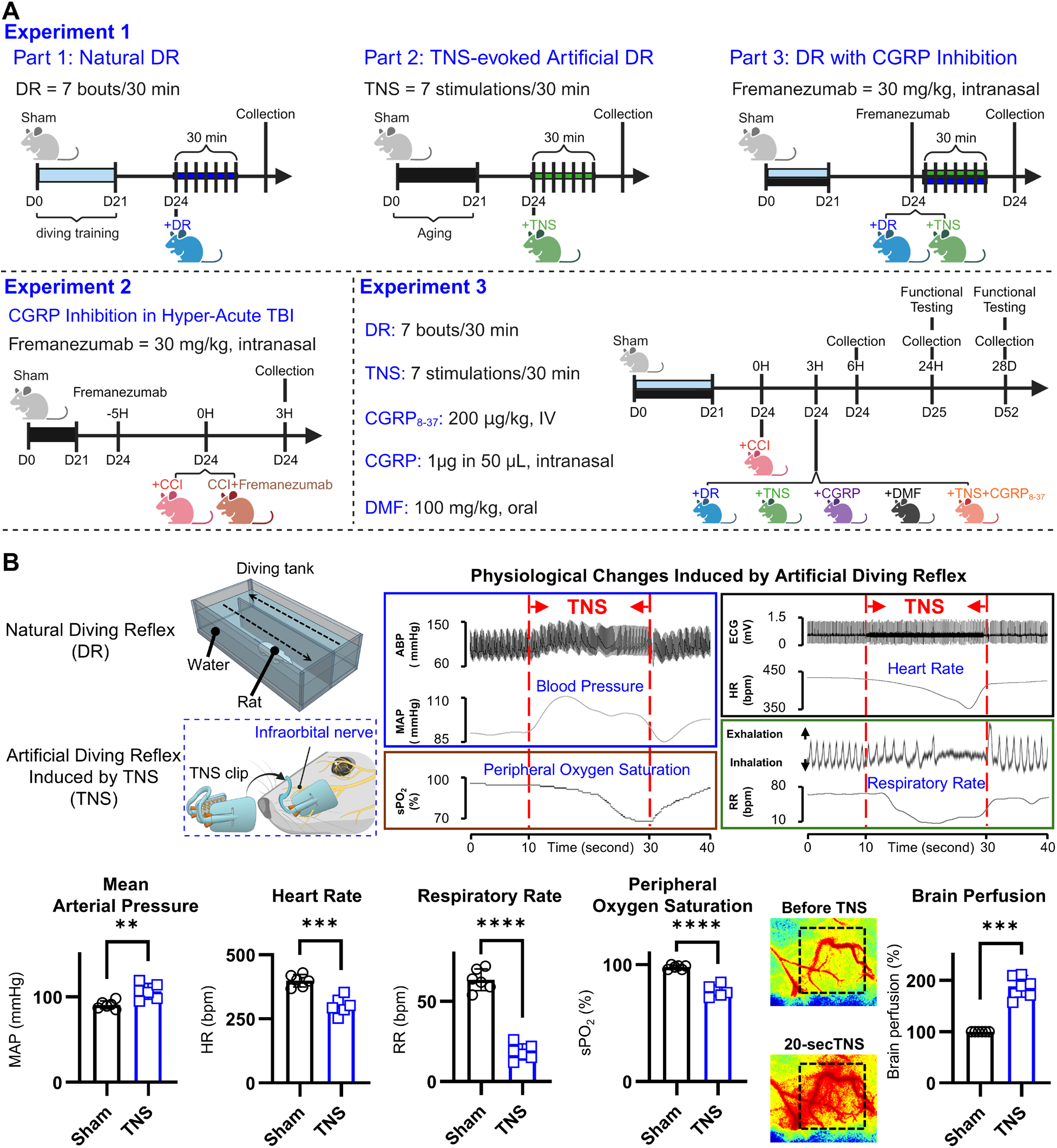
Experimental design and physiological validation of TNS-induced artificial diving reflex. **(A)** Schematic overview of the three experimental protocols. Experiment 1: Comparison of natural diving reflex (DR) versus TNS-induced DR (TNS), with and without CGRP inhibition. Experiment 2: Assessment of hyper-acute (3-hour) TBI with and without CGRP inhibition. Experiment 3: Acute and chronic evaluation of TBI with different therapeutic interventions (DR, TNS, CGRP, DMF) delivered at 3 hours post-injury. **(B)** Physiological validation of TNS-induced artificial DR. Representative traces demonstrate that brief 20-sec TNS trains induce the characteristic hemodynamic signature of the natural DR, including increased mean arterial pressure (MAP), decreased peripheral oxygen saturation (SpO₂), bradycardia (HR), and apnea (RR). This systemic response is accompanied by a significant enhancement of cerebral perfusion. Data are presented as mean ± standard deviation (SD). Statistical significance was determined by a paired t-test comparing values during the 20-second TNS period to the pre-stimulation baseline, where \**P* < 0.05, \*\**P* < 0.01, \*\*\**P* < 0.001, and \*\*\*\**P* < 0.0001.

#### Experiment 1 design

To systematically characterize the diving reflex and its artificial induction, 60 rats were allocated to this study. Following the exclusion of three animals that failed to meet predetermined performance criteria during conditioning protocols, the remaining cohort was randomized into five experimental groups (n=12 each): (1) natural diving reflex (DR), (2) trigeminal nerve stimulation induced artificial diving reflex (TNS), (3) DR with CGRP antagonism (DR+aCGRP), (4) TNS with CGRP antagonism (TNS+aCGRP), and (5) a sham control group. Animals in the CGRP antagonist groups received Fremanezumab (30 mg/kg, intranasal) 5 hours prior to their final DR or TNS session. Each experimental group was further subdivided (n=6 each) for downstream processing: one half was designated for fresh tissue harvest and subsequent biochemical analyses, while the other half underwent transcardial perfusion for histological and immunofluorescence evaluation. The small number of animals that did not meet conditioning criteria were reallocated for use in Experiments 2 or 3 to minimize animal usage.

#### Experiment 2 design

To investigate the role of endogenous CGRP signaling in the hyper-acute phase of TBI, 36 animals were systematically randomized into three experimental groups: (1) sham surgery, (2) TBI with vehicle, and (3) TBI with CGRP antagonism (TBI+aCGRP). Animals in the antagonist group were pre-treated with Fremanezumab. All animals were sacrificed at 3 hours post-injury. Each cohort (n=12) was equally divided (n=6 each) for downstream analysis, with one half designated for fresh tissue harvest and biochemical assays, and the other half for transcardial perfusion and histological processing.

#### Experiment 3 design

To evaluate the acute and chronic efficacy of DR-based interventions, animals were allocated to a controlled cortical impact (CCI) model of severe TBI with a complex factorial design. Animals were systematically randomized across three distinct survival timepoints, 6 hours (hyper-acute), 24 hours (acute), and 28 days (chronic), and up to seven different treatment conditions: (1) sham surgery, (2) CCI with vehicle, (3) CCI + natural DR, (4) CCI + artificial TNS-induced DR, (5) CCI + dimethyl fumarate (DMF), (6) CCI + TNS with CGRP inhibition, and (7) CCI + exogenous CGRP. The cohort distribution was tailored to the specific endpoints at each time point. The 6-hour survival cohort included all seven treatment conditions, though the CCI+TNS-CGRP and CCI+CGRP groups only underwent perfusion (6 animals/group for fresh tissue collection and biochemical analysis, and 6/group for perfusion and histological analysis), totaling 72 animals. The 24-hour survival cohort consisted of six treatment groups (excluding CCI+TNS+aCGRP), with 8 animals per group (totaling 48 animals) designated for perfusion following functional assessments. The 28-day survival cohort focused on the long-term efficacy of TNS and consisted of three groups (sham, CCI, CCI+TNS), with 8 animals per group (totaling 24 animals), all of whom underwent perfusion after the final behavioral testing battery. Overall, animals in the 24-hour and 28-day cohorts were primarily used for histological and functional analyses, while the 6-hour cohort provided tissues for both biochemical and histological evaluations.

### Rat voluntary diving protocol

Rats were trained to voluntarily traverse a multi-channel underwater tunnel using a modified version of a validated protocol [17]. All 228 animals completed the same training regimen, which required unassisted swimming by day two, consistent voluntary dives by the end of week one, and minimal agitation when handled or submerged; six rats were excluded prior to experimental group assignment for failing to meet these criteria. Rats were trained within a rectangular tank (100 x 38 x 15 cm) comprised of ¾-in thick acrylic pieces, with a ½-in thick top (MazeEngineers, Skokie, IL). The tank was modified to include 2 channels instead of the 5 channels specified in the original protocol, or the 3 channels we have used previously, due to the functional limitations of the animals following TBI [17]. During a one-week period, the rats were acclimated to the aquatic environment and trained to voluntarily swim through the two-channel route, covering a total distance of approximately 1.5 meters. The training protocol involved initially acclimating the rats to the water environment to ensure their comfort and familiarity, followed by a gradual increase in the distance they were required to swim, ultimately covering the total 1.5 meters at a depth of approximately 10 cm in water maintained at 32±2°C. This temperature was chosen to prevent the development of hypothermia in the animals and reduce stress when undergoing full-body submersion, as established by the original source of the model [19], despite indications in humans that DR exhibits a temperature-dependent effect [20]. Rats were trained daily, with 3-5 trials per day per rat. Each rat was placed in the water at increasing distances from the finish platform. When the rats showed consistent successful negotiation of the path, they started on dive training to last another 2-weeks. It consisted of first developing the rats’ ability to dive underwater comfortably and voluntarily, and then gradually increasing the length of the underwater portion of the swimming path, until they showed consistent negotiation of the underwater multi-channel tunnel. Rats were required to show swimming ease within the first two days of training, voluntarily initiate dives within the first week of diving training and display minimal signs of agitation when handled or diving. Upon successful completion of training, rats were subjected to a 30-minute diving session 3 days later, involving repeated dives every 5 minutes for a total of 7 dives, followed by sample collection. Rats in the anti-CGRP group were administered Fremanezumab (Ajovy, 30 mg/kg, i.n., Teva Pharmaceuticals Industries Ltd., Parsippany, NJ) to inhibit CGRP 5 hours before the final session and histological collection. All rats underwent swim and diving training, however only natural DR animals underwent sessions of repetitive DR. Three days after training completion, rats assigned to the DR group underwent a 30-minute session comprising seven dives at 5-minute intervals, after which samples were collected. Remaining rats were randomly assigned to other experimental groups.

### TNS-induced artificial diving reflex

To induce a controllable artificial DR, rats received non-invasive trigeminal nerve stimulation (TNS) via a custom-engineered nasal clip stimulator designed to bilaterally target the infraorbital nerve at the whisker barrel (**Figure 1B**). This method mimics the specific trigeminal afferent input experienced during natural facial immersion. The clip electrode, consisting of two conductive pads, delivered rectangular biphasic pulses (50 Hz, 0.5 ms pulse width, 2 V amplitude) generated by an Isolated Pulse Stimulator (Model 2100, A-M Systems, Sequim, WA, USA). To replicate the temporal pattern of natural, intermittent diving, stimulation was applied in 20-second trains every 5 minutes over a 30-minute session [18]. In healthy animals (Experiment 1), this TNS session was administered three days after the completion of the DR training regimen. In the TBI model (Experiment 3), the TNS session was initiated 3 hours post-injury.

### Physiological recording of TNS-induced artificial diving reflex

To quantify the physiological effects of TNS-induced artificial DR, age-matched, naïve rats were utilized. Animals were anesthetized with 5% isoflurane and surgically instrumented with a catheter in the left femoral artery for continuous recording of arterial blood pressure (ABP) and mean arterial pressure (MAP). Each rat was then secured in a stereotaxic apparatus where a surgical plane of anesthesia was maintained at 1.5% isoflurane, allowing for spontaneous respiration while core body temperature was kept at 37.0±0.2°C via a homeothermic monitoring system. Key cardiorespiratory parameters were continuously recorded. Subcutaneous needle electrodes (MLA1213, ADInstruments, USA) were placed in the limbs to monitor electrocardiogram (ECG) and heart rate (HR). A thermocouple (MLT1402, ADInstruments, USA) positioned at the nose measured temperature changes between inspiration and expiration to calculate respiratory rate (RR). A pulse oximeter on the hindlimb (MLT323, ADInstruments, USA) measured peripheral oxygen saturation (SpO₂). Cerebral blood flow (CBF) was measured using a Laser Speckle Perfusion Imaging System (RFLSI III, RWD Life Science, San Diego, CA, USA). A midline scalp incision was made to expose the skull, and the LSF camera was positioned 30 cm above the surface using an angulating arm. Imaging was performed at a display rate of 25 Hz with a 1-second time constant and a 3-millisecond camera exposure. A single rectangular region of interest was identified on the left hemisphere. After a 1-minute baseline recording, CBF flux was measured continuously during the 20-second TNS period. All physiological signals were digitized at a sampling rate of 1 KHz using a PowerLab 16/SP data acquisition system and were analyzed with LabChart 7.0 software (ADInstruments, USA). For cardiorespiratory and hemodynamic parameters (MAP, HR, RR, SpO₂), the effect of stimulation was calculated as the mean value during the 20-second TNS period. For cerebral blood flow, results are presented as the average flux during the 20-second TNS period, normalized to the pre-stimulation baseline.

### Controlled cortical impact model and therapeutic interventions

Three days after completing DR training, severe TBI was induced using a well-characterized controlled cortical impact (CCI) paradigm [21]. Animals were anesthetized with 5% isoflurane and positioned in a stereotaxic apparatus where a surgical plane was maintained at 1.5% isoflurane and core body temperature was kept at 37.0±0.2°C. A 6-mm diameter craniotomy was performed over the left parietal cortex (4 mm lateral to the sagittal suture, equidistant between bregma and lambda). A cortical impact was delivered using an electromagnetic device (Leica Biosystems) with the following parameters: 6.0 m/s velocity, 3.0 mm deformation depth, and 100 ms dwell time. Sham animals underwent identical procedures excluding the impact. Post-surgical wound closure was achieved with 4-0 silk sutures and rapid-seal wound gel. Animals were pre-randomized to survival endpoints of 3 h, 6 h, 24 h, or 28 days to accommodate Fremanezumab pre-administration where required. A critical 3-hour post-injury timepoint was established for intervention based on our data showing it as the peak of hyperacute NRF2 dysregulation. At this 3-hour mark, animals in Experiment 2 were collected for biochemical analysis. For Experiment 3, animals were further randomized to receive one of the following single-dose treatments: (1) a 30-minute natural DR session, (2) a 30-minute TNS session, (3) intranasal CGRP (1 µg in 50 µL saline; Tocris Biosciences), or (4) oral gavage of DMF (100 mg/kg in 0.6% methylcellulose; MilliporeSigma). Sham and vehicle groups concurrently received oral saline, while the sham, vehicle, CGRP, and DMF groups experienced nasal clip placement and handling, without current delivery, for parity. Importantly, pilot testing confirmed that animals retained the capacity to perform the diving task at 3 hours post-injury, provided analgesics were withheld until after the session. Therefore, analgesic management consisting of subcutaneous buprenorphine (0.05–0.1 mg/kg) with topical antibiotic prophylaxis was administered immediately *after* the 30-minute DR or TNS intervention. Following treatment and analgesia, animals were monitored in temperature-controlled recovery chambers until achieving sternal recumbency before returning to their home cages.

### Multi-domain functional assessments

To comprehensively evaluate the therapeutic efficacy of each intervention, functional outcomes were assessed at 24 hours (acute) and 28 days (chronic) post-TBI using a battery of behavioral tests designed to probe sensorimotor, cognitive, affective, and nociceptive functions. All behavioral testing was conducted by an experienced technician blinded to the treatment groups. Animal activity and location were tracked using EthoVision XT video software (Version 17, Noldus Information Technology, Netherlands) and manually verified for accuracy. Between each trial, all surfaces and apparatuses were thoroughly cleaned with a hydrogen peroxide solution followed by 75% ethanol to eliminate olfactory cues.

#### Sensorimotor function

Sensorimotor function was assessed using the adjusted neurological severity score (A-NSS), a validated composite neurobehavioral assessment tool adapted from the neurological severity score protocol. This comprehensive scoring system quantifies both global and domain-specific neurological deficits across multiple functional modalities. The A-NSS battery evaluates locomotor patterns, resistance to passive manipulation, hindlimb and forelimb motor function, proprioceptive responses, righting reflexes, exploratory behavior, cranial nerve reflexes, visuospatial orientation, and vestibulomotor coordination. Individual test components are scored on a graded scale, with higher numerical values indicating greater functional impairment. The cumulative score ranges from 0 (normal neurological function) to 45 (maximal neurological deficit). Baseline neurological testing was performed 24-hours prior to injury induction to establish pre-injury functional status. All animals demonstrated intact neurological function during baseline assessment; consequently, no subjects required exclusion based on pre-injury evaluation criteria.

#### Short-term memory

Short-term memory was assessed using the novel-object recognition (NOR) test. Animals are exposed to one object prior to the test itself with which to become familiarized within a 50cm x 50cm enclosure with opaque black walls. After a 1-hour interval, the animals are placed back into the enclosure with two objects. One object is the familiarized object and the other object is a novel object comprised of a similar size and material. The animal is placed equidistant from each object and allowed to freely interact for 5-minutes, within which the amount of time spent interacting with each object is recorded and expressed as a discrimination ratio with a value from 0.0 (only interaction with familiar object) to 1.0 (only interaction with novel object). An animal with intact short-term memory will interact more with the novel object, rather than the familiar one.

#### Anxiety-like behavior

To assess anxiety-like behavior, rats underwent the elevated plus maze assessment. The maze consists of four arms set perpendicularly to one another to form a cross. Two arms are open to the outside and two arms are surrounded by opaque walls on three sides, to form semi-enclosed spaces. Each rat is placed in the center of the maze facing towards an open arm and away from the assessor. The rat is allowed 5 minutes to explore the maze freely; the number of entries into each arm is recorded. This is represented as percentage of total entries/time, with a higher percentage indicating a less anxious state. If a rat freezes or falls from the maze, it is disqualified from the test. No rats were disqualified from testing during the course of this experiment.

#### Pain threshold

The nociceptive withdrawal threshold was assessed using the Randall-Selitto electronic algesimeter (IITC 2500 Digital Paw Pressure Meter, IITC Life Science, Woodland Hills, CA). Each animal was firmly held by a technician to whom the animals had been previously familiarized. A separate technician applied an increasing mechanical force on the plantar surface of the contralateral forepaw until a withdrawal response was observed.

### Fresh tissue and serum collection

At the designated survival endpoints, animals allocated for biochemical analysis were deeply anesthetized with isoflurane. Blood samples were collected via cardiac puncture from the vena cava, transferred to serum separator tubes, and allowed to clot at 4°C for 30 minutes. The samples were then centrifuged at 2000 × *g* for 10 minutes to isolate the serum, which was carefully aliquoted and stored at −80°C. Immediately following blood collection, the animals were decapitated, and the brains were rapidly extracted. The brains were placed on an ice-cold dissection matrix where the infarct, peri-infarct, and contralateral cortices were microdissected. These tissue samples were immediately flash-frozen in liquid nitrogen and stored at −80°C for subsequent molecular analyses.

### Transcardial perfusion and tissue processing

At their designated survival endpoints, a separate cohort of animals was deeply anesthetized and transcardially perfused to prepare brain tissue for histological analysis. The circulatory system was first flushed with 0.1 M ice-cold phosphate-buffered saline (PBS; Sigma, St. Louis, MO, USA), followed by fixation with 4% paraformaldehyde (PFA) in PBS. Following perfusion, the brains were carefully extracted, post-fixed overnight in 4% PFA at 4°C, and then cryoprotected via sequential immersion in graded sucrose solutions (10%, 20%, and 30% in PBS). The brains were then embedded in a mixture of 30% sucrose and optimal cutting temperature (OCT) compound (Electron Microscopy Sciences, Hatfield, PA, USA), flash-frozen, and stored at −80°C. Serial coronal sections were cut at a thickness of 14 µm using a cryostat (Leica Biosystems, Deer Park, IL, USA) at 400-µm intervals throughout the entire rostrocaudal extent of the brain, yielding 28-32 sections per specimen. Sections were mounted onto Polysine-coated glass slides (Thermo Fisher Scientific, Waltham, MA, USA) and stored at −80°C until histological and immunofluorescence staining.

### Proteomics analysis

Aliquots of pulverized brain tissue from sham (N=4), DR (N=4) and TNS (N=4) animals underwent comprehensive proteomic analysis via liquid chromatography-tandem mass spectrometry at Creative Proteomics (Shirley, NY, USA). Raw spectral data underwent standardized preprocessing including peak detection, peptide identification, and protein quantification by Creative Proteomics, who were blinded to any specific groups. The resulting data was analyzed using a multi-platform bioinformatics workflow integrating R (MaxQuant v2.6.7.0, XLSTAT (v2024.4.2), and Python (Matplotlib v3.10.0) analytical environments. Data was subjected to median normalization followed by log-2 transformation. Missing intensity values were addressed through a hierarchical imputation strategy based on data sparsity patterns: k-Nearest Neighbor (kNN) for proteins with moderate missingness (n=1 missing values), Quantile Regression Imputation for proteins with substantial missingness (n=2 or 3 missing values), and Minimum Value Imputation for proteins with complete absence in one experimental group (n=4 missing values). Differentially expressed proteins were identified using dual selection criteria comprising Log2 fold-change thresholds >1.0 or <-1.0 combined with statistical significance (Student’s t-test with Benjamini-Hochberg correction, p<0.05). Visual representation of the proteomic landscape was accomplished through volcano plots and clustered heat maps generated using Python Matplotlib. Further protein-protein interaction, and potential effects in humans, was examined and visualized using the Gene Ontology Consortium, the Protein ANalysis THrough Evolutionary Relationships (PANTHER) classification system, and STRING network analysis, using both cluster and functional enrichment visualization keyed for human proteins.

### Subcellular fractionation and Western blotting

Frozen cortical tissue samples were pulverized under liquid nitrogen, and all subsequent processing and analysis were performed by personnel blinded to the treatment groups. To assess subcellular protein localization, nuclear and cytoplasmic fractions were prepared using a commercial extraction kit (**Table S1**). Briefly, tissue homogenates were first incubated in a cytoplasmic extraction buffer and centrifuged, with the resulting supernatant collected as the cytoplasmic fraction. The remaining nuclear pellet was then resuspended in a nuclear extraction buffer, subjected to high-speed vortexing, and centrifuged to isolate the nuclear fraction. Total protein concentrations for nuclear, cytoplasmic, and whole-tissue lysates (prepared using RIPA buffer supplemented with protease and phosphatase inhibitors) were determined using a BCA assay. Equal amounts of protein were resolved by SDS-PAGE, transferred to polyvinylidene difluoride (PVDF) membranes, and blocked for 1 hour. Membranes were incubated overnight at 4°C with validated primary antibodies (**Table S2**) [22–29], followed by a 1-hour incubation with HRP-conjugated secondary antibodies (Abcam, Waltham, MA). Protein bands were visualized using enhanced chemiluminescence (ECL) on a Bio-Rad ChemiDoc System. Densitometry was performed using ImageJ software, with loading controls β-actin and Lamin B1 used for cytoplasmic/total and nuclear fractions, respectively. Each Western blot was independently replicated twice. Data are presented as fold change relative to sham controls. For proteins where the ratio of phosphorylated to total expression was calculated (e.g., p62) or for the LC3-II/I ratio, values were derived from the β-actin normalized expressions of each band rather than from the final fold-change data.

### Glutathione quantification

To assess the redox state, levels of reduced (GSH) and oxidized (GSSG) glutathione were measured in both pulverized brain tissue samples and serum. All analyses were performed by personnel blinded to the treatment groups. The quantification of GSH, GSSG, and the resulting GSH/GSSG ratio was carried out using a commercial colorimetric assay kit according to the manufacturer’s protocol (**Table S1**). Each sample was run in technical duplicate for confirmation. Tissue concentrations are reported as µmol per gram of tissue (µmol/g), while serum concentrations are presented as µmol per milliliter of serum (µmol/mL).

### Measurement of malondialdehyde levels

To evaluate oxidative damage, lipid peroxidation was quantified by measuring malondialdehyde (MDA) levels in both pulverized brain tissue and serum samples. All analyses were conducted by personnel blinded to the experimental groups. A commercial assay kit (**Table S1**) was used, which measures thiobarbituric acid reactive substances (TBARS) as an index of MDA concentration. Tissue samples were processed following a protocol optimized for high-protein content to ensure accurate quantification. Each sample was analyzed in technical duplicate, and results are reported as a measure of lipid peroxidation levels.

### Multiplex immunofluorescence and quantitative analysis

To evaluate the spatial and cellular distribution of key signaling molecules and oxidative stress markers, all immunofluorescence staining and subsequent image analysis were performed by personnel blinded to the experimental groups. Cryosectioned brain slides were blocked with 5% goat serum (Abcam, Waltham, MA) in 1% bovine serum albumin (Sigma, Burlington, MA) for 1 hour at room temperature. The slides were then incubated overnight at 4°C with primary antibodies against CGRP, 4-hydroxynonenal (4-HNE), or nitrotyrosine (**Table S2**), followed by a 1-hour room temperature incubation with their corresponding fluorophore-conjugated secondary antibodies. Subsequently, slides were co-stained with a primary antibody against phospho-NRF2 (Ser40) and its corresponding secondary antibody. All primary antibodies used were selected based on their validation across multiple species and experimental conditions [27,28,30–34]. Finally, nuclei were counterstained with DAPI, and the slides were mounted using Vectashield Antifade Mounting Medium (Vector Laboratories, Burlingame, CA). High-resolution, whole-brain images were captured using an EVOS M7000 imaging system (Thermo Fisher Scientific, Waltham, MA) with a 20X objective and an automated XY-stitching function. ImageJ software was used for semi-automated, quantitative analysis of signal expression. Pixel intensities, ranging from 0 (weakest) to 255 (strongest), were converted to reciprocal intensity (arbitrary units, AU) for data presentation. To assess the relationship between NRF2 activation and intracellular oxidative stress, co-expression of phospho-NRF2 and 4-HNE was quantified within 20 randomly selected cells per region of interest (ROI), with three ROIs analyzed per anatomical region per sample. A density ratio of 4-HNE to phospho-NRF2 was calculated to determine the balance between oxidative burden and the antioxidant response. Similarly, to quantify p62 activation specifically within neurons, the expression of total p62 and phospho-p62 (Ser351) was measured in 20 NeuN-positive cells per ROI (three ROIs per region). The average phospho-p62(Ser351) signal was then divided by the average total p62 signal to generate a neuronal activation ratio, analogous to the ratio calculated from Western blot data.

### Histological analysis of lesion volume

To quantify cortical tissue loss, tissue sections mounted on Superfrost Plus slides were processed for standard hematoxylin and eosin (H&E) staining by personnel blinded to all experimental groups. Digital whole-slide images were acquired using a Pathscan Enabler 5 imaging system (Meyer Instruments, Houston, TX, USA). Quantitative lesion analysis was performed with ImageJ software (NIH, Bethesda, MD, USA) by manually delineating the boundaries of the lesion, identified by loss of normal tissue architecture and cellular staining, in each coronal section. Total lesion volume was calculated by integrating the cross-sectional lesion areas across all 28 serial sections obtained from each specimen. To account for inter-animal variability in brain size and post-traumatic edema, the final lesion volume was normalized to the volume of the contralateral hemisphere and expressed as a percentage.

### Statistical analysis

All quantitative data are presented as mean ± standard deviation (SD), with individual data points displayed on graphs to illustrate data distribution. Sample sizes were determined by *a priori* power analysis α=0.05, β=0.8), establishing a minimum of n = 6 per group for molecular and histological endpoints and n = 8 per group for neurobehavioral assessments to account for higher anticipated inter-subject variability. The normality of each dataset was confirmed using the Shapiro–Wilk and Kolmogorov–Smirnov tests, and homogeneity of variance was verified using the Brown–Forsythe test. For acute physiological recordings comparing pre-stimulation baseline to intra-stimulation values within the same animals, statistical significance was determined using a paired two-tailed Student’s t-test. For experiments involving multiple treatment arms (e.g., Experiment 1: sham, DR, TNS, DR+aCGRP, TNS+aCGRP; Experiment 2 and 3: sham, CCI, and respective treatment cohorts), between-group comparisons were performed independently within each anatomical region and survival timepoint using ordinary one-way analysis of variance (ANOVA) followed by Tukey’s post-hoc test for multiple comparisons. Associations between continuous biochemical variables were evaluated using simple linear regression and reported as the coefficient of determination (R^2^). Statistical significance was defined *a priori* at P < 0.05 (denoted as *P < 0.05, **P < 0.01, ***P < 0.001, and ****P < 0.0001). All univariate statistical analyses were executed using GraphPad Prism software (version 9.0; GraphPad Software, San Diego, CA, USA). For the visualization and quantification of multi-dimensional recovery profiles, three-dimensional scatterplots with transposed 95% confidence interval ellipsoids were constructed in Python (v3.10). Group centroids were calculated in three-dimensional feature space, and the Euclidean distance (d = √[(x₂–x₁)² + (y₂–y₁)² + (z₂–z₁)²]) between each treatment group centroid and the sham control centroid was computed as a quantitative metric of global phenotypic normalization.

## Results

### Non-invasive trigeminal nerve stimulation recapitulates the hemodynamic signature of the diving reflex

To establish a clinically feasible method for inducing artificial DR, we first validated a non-invasive TNS protocol. In lightly anesthetized rats, brief 20-second trains of TNS (50 Hz, 0.5 ms, 2 V) targeting the infraorbital (V2) nerve branch reproducibly evoked the complete tetrad of physiological changes that define natural DR [18]. Stimulation induced a hypertensive response, with mean arterial pressure (MAP) increasing from 91±5 to 106±9 mmHg, alongside peripheral vasoconstriction, evidenced by a drop in peripheral oxygen saturation (SpO₂) from 98±2% to 78±5%. Concurrently, animals exhibited bradycardia, with heart rate (HR) slowing from 399±24 to 302±34 bpm, and a near-complete intermittent apnea, with respiratory rate (RR) decreasing from 64±7 to 19±5 breaths/min (**Figure 1B**). Critically, this systemic autonomic response was coupled with a profound enhancement of brain perfusion, increasing cerebral perfusion by 88±21% over baseline. These convergent findings confirm that our TNS protocol reliably induces the full, integrated physiological state of the natural DR, from systemic autonomic shifts to targeted cerebral hyperemia [35].

### Tuned TNS induces a more potent and consistent antioxidant proteomic signature than the natural DR

To define the molecular signature of DR and determine if our artificial TNS model could replicate or even enhance it, we performed untargeted liquid chromatography mass spectrometry on cortical tissue from sham, natural DR, and TNS-treated rats. The analysis of 2,388 quantified proteins revealed a profound and distinct proteomic shift induced by both interventions. A single 30-minute DR session resulted in 485 differentially expressed proteins, while TNS elicited an even broader response, altering 732 proteins relative to sham controls (**Figure 2A, SF1**). Functional pathway mapping demonstrated a striking focus on oxidative stress. In natural DR-exposed animals, 69% of the 29 most significantly altered proteins belonged to this category. TNS-induced DR showed a similarly targeted, though more extensive, effect, with 61% of 46 significant proteins being oxidative stress-related. Functional enrichment analysis (**Figure SF2**) and ranked heat mapping further highlighted a pronounced upregulation of canonical NRF2 targets, including *GPX1*, *GSS*, *GLRX2*, *TXNRD2*, and *SOD1*, that drive peroxide and superoxide detoxification and boost glutathione synthesis [36–38]. In parallel, classical stress markers (*ADH7*, *NDRG1*, *TXNRD3*, *MAOB*, *MDH1*, *AK4*) were markedly downregulated, consistent with reduced ROS generation [10,39]. This signature, the induction of antioxidant defenses coupled with the quieting of stress responses in the absence of electrophilic challenge, provides compelling evidence that the DR activates the NRF2 pathway via a beneficial, redox-neutral mechanism. Critically, while natural DR and TNS produce qualitatively similar proteomic changes, quantitative analysis suggests two key advantages of the TNS model, when compared to this specific form of natural DR. First, the TNS-induced changes tended to be of a relatively greater magnitude. Second, the inter-animal variance was generally reduced compared to the natural DR group. Together, these data demonstrate that TNS not only faithfully recapitulates the DR’s antioxidant program but also has the capacity for superior potency, consistency, and control, establishing it as a promising therapeutic modality for translation to TBI.

**Figure 2.**
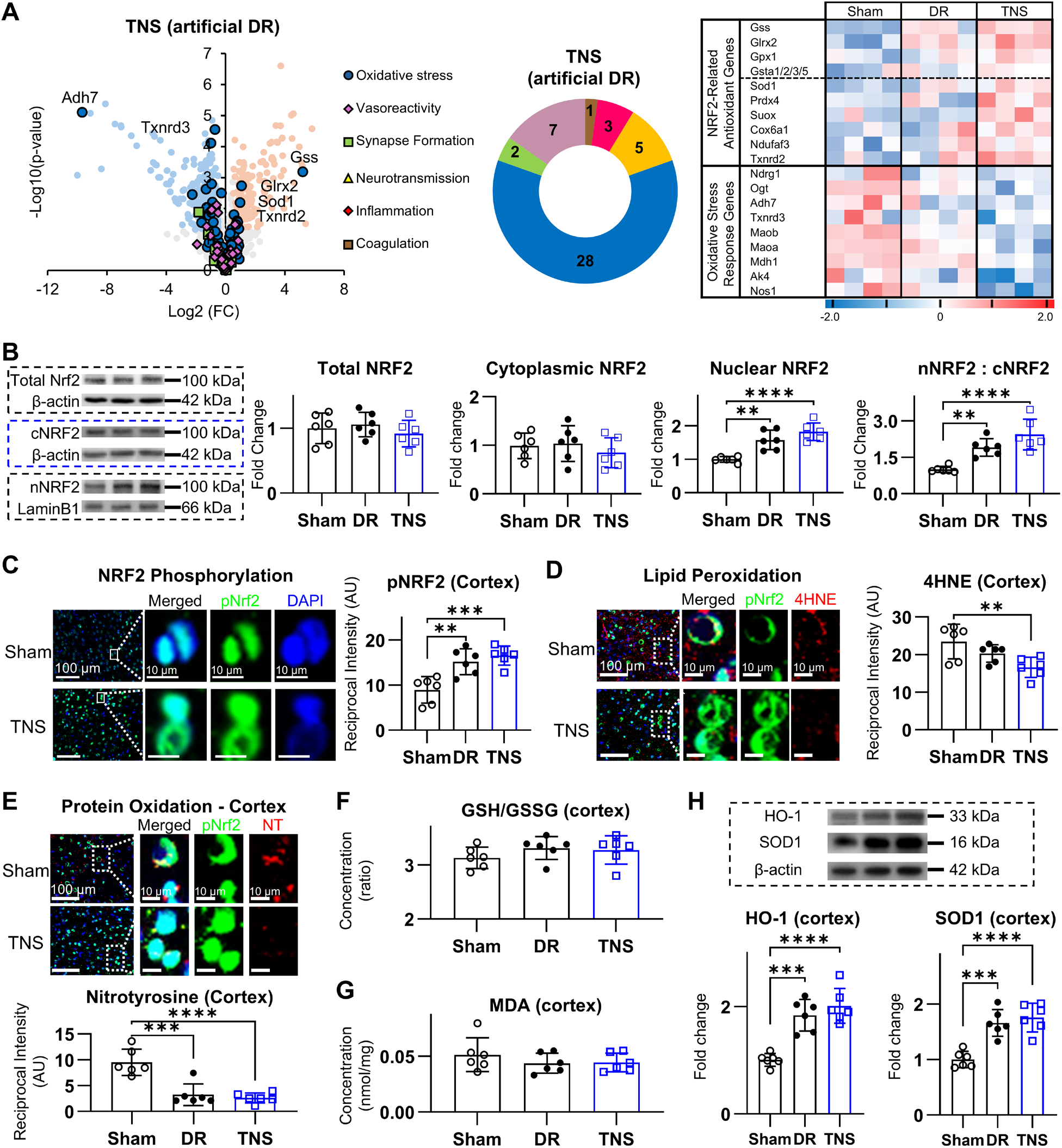
TNS-induced artificial diving reflex (DR) recapitulates and potentiates the natural DR’s non-electrophilic antioxidant response. **(A)** Cortical proteomic analysis shows TNS recapitulates natural DR’s upregulation of NRF2 antioxidant targets and downregulation of stress markers. The TNS-induced DR response is more potent and consistent. **(B)** Immunoblotting demonstrates increased NRF2 nuclear translocation 30 minutes after DR and TNS, with no change in total NRF2. **(C)** Immunofluorescence reveals increased phospho-NRF2 (pNRF2 at ser40) in the cortex after both interventions. **(D-E)** DR and TNS decrease cortical markers of lipid peroxidation (4-HNE) and protein oxidation (3-NT). **(F-G)** The glutathione redox ratio (GSH/GSSG) and malondialdehyde (MDA) levels are unchanged post-intervention, confirming a non-electrophilic, redox-neutral mechanism. **(H)** Both interventions increase protein expression of the NRF2 targets HO-1 and SOD1. Data are presented as mean ± SD. For comparisons between the sham, DR, and TNS groups, statistical significance was determined by one-way ANOVA followed by Tukey’s post-hoc test. Asterisks denote significance relative to the sham control group: \**P* < 0.05, \*\**P* < 0.01, \*\*\**P* < 0.001, and \*\*\*\**P* < 0.0001.

### Rapid, non-electrophilic NRF2 activation by natural and TNS-induced DR

We next investigated whether TNS-induced artificial DR recapitulates the rapid, non-electrophilic activation of the antioxidant master regulator NRF2 observed with diving-induced natural DR. Cortical tissues collected 30 minutes after a single DR or TNS session were subjected to subcellular fractionation and immunoblotting (**Figure 2B**). Both interventions produced a ∼1.5-fold increase in nuclear NRF2 and a ∼1.9-fold rise in the nuclear-to-cytoplasmic ratio, without altering total NRF2 levels, confirming enhanced nuclear translocation. Phospho-NRF2 (Ser40) immunofluorescence in the dorsal cortex, hippocampal CA1, amygdala, and thalamus, regions stratified by vulnerability in TBI, revealed significant fluorescence increases (1.6–2.0-fold; **Figure 2C**, **SF3**), indicating rapid pathway engagement. Interestingly, TNS-induced artificial DR induced a greater increase in phospho-NRF2 than natural DR within the amygdala and thalamus (**SF3**), intimating an increased efficacy or efficiency. To confirm the absence of electrophilic stress, we quantified lipid peroxidation (4-HNE) and protein oxidation (3-nitrotyrosine) by immunostaining: 4-HNE decreased by 15–40% across all regions, while 3-nitrotyrosine decreased by 30-80% (**Figure 2D-E**, **SF4**). High-performance assays of glutathione redox status (GSH/GSSG) and malondialdehyde (MDA) showed no significant changes versus sham (**Figure 2F-G**), while canonical NRF2 targets heme oxygenase-1 (HO-1) and superoxide dismutase-1 (SOD1) [40,41] were upregulated ∼1.8-2.0 fold (**Figure 2H**). Collectively, these findings demonstrate that both natural and TNS-induced DR rapidly activate NRF2-dependent antioxidant defenses through a mechanism that preserves, rather than consumes, cellular antioxidant reserves.

### CGRP is a necessary mediator of NRF2 activation in natural and TNS-induced DR

Given that both natural DR and its TNS surrogate rapidly activate NRF2, we next sought to confirm the upstream mediator responsible. We focused on CGRP, a neuropeptide released by trigeminal afferents that enhances NRF2 signaling [16] and drives its phosphorylation at Ser40 [21] (**Figure 3A**). Importantly, CGRP’s peptide structure lacks electrophilic moieties (**Figure SF5A**), making it an ideal candidate for mediating a redox-neutral response.

**Figure 3.**
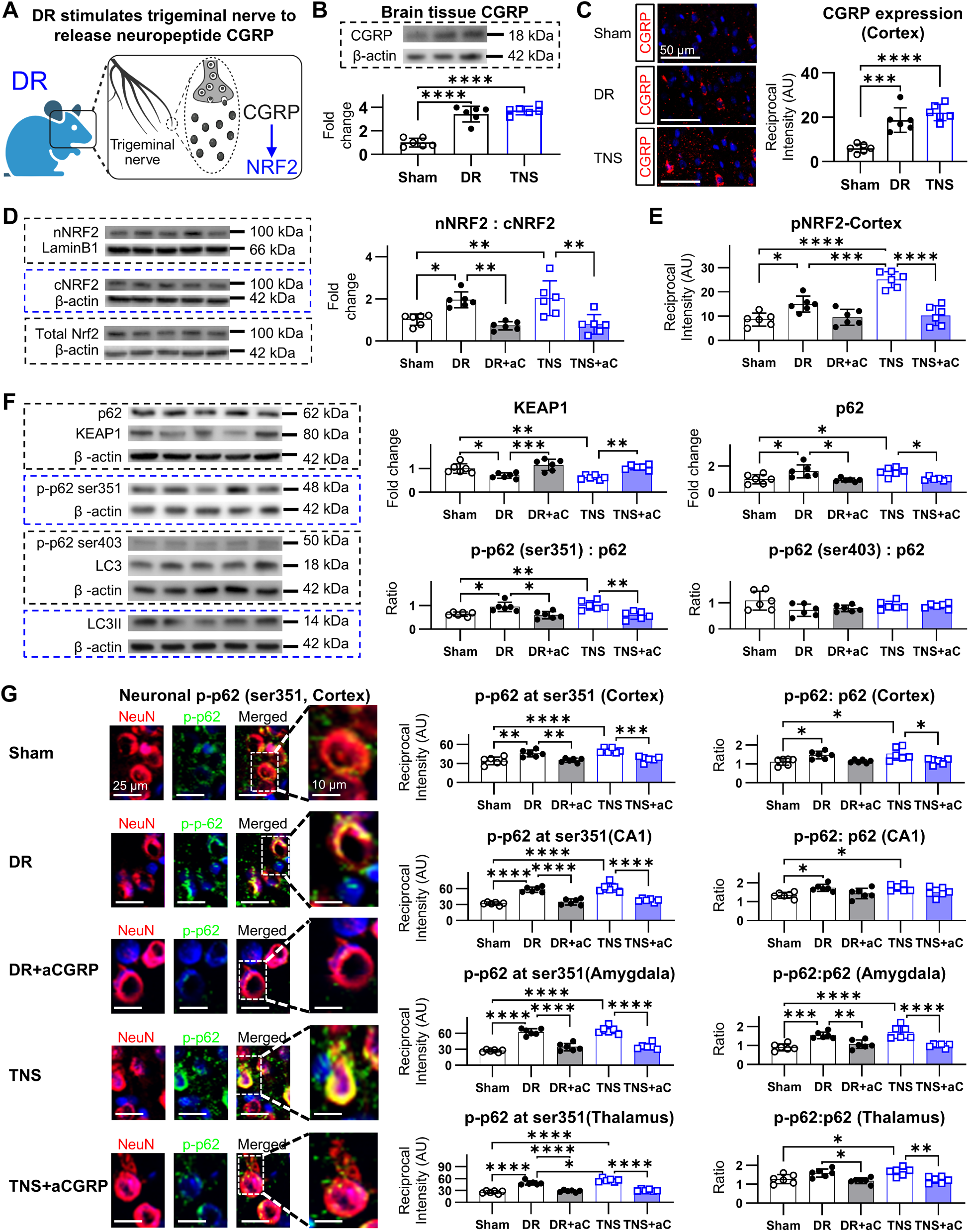
CGRP mediates NRF2 activation by selectively engaging the p62-KEAP1 pathway. **(A)** Schematic illustrating the proposed signaling cascade: Trigeminal stimulation (DR/TNS) releases CGRP, which drives NRF2 activation via the p62-KEAP1 axis. **(B)** Both DR and TNS induce a significant increase in cortical CGRP levels. **(C)** Immunofluorescence reveals a spatial co-localization of increased CGRP expression and phospho-NRF2 (ser40) in the cortex and hippocampus following TNS. **(D)** Pharmacological blockade of CGRP with Fremanezumab abolishes TNS-induced NRF2 nuclear translocation. **(E)** CGRP blockade also prevents the TNS-induced increase in phospho-NRF2 (ser40), confirming CGRP is required for this activation step. **(F)** DR and TNS increase total and, critically, phospho-p62 (ser351) levels, leading to a concurrent decrease in KEAP1 protein, indicating selective engagement of the NRF2 activation pathway. **(G)** CGRP blockade abrogates the TNS-induced phosphorylation of p62 and subsequent reduction of KEAP1, demonstrating that the entire p62-KEAP1-NRF2 cascade is CGRP-dependent. Data are presented as mean ± SD. Statistical significance for comparisons across all groups was determined by one-way ANOVA followed by Tukey’s multiple comparisons test. Asterisks denote significance relative to the sham control group, where \**P* < 0.05, \*\**P* < 0.01, \*\*\**P* < 0.001, and \*\*\*\**P* < 0.0001.

Consistent with this, a single 30-minute session of either DR or TNS increased cortical CGRP levels by ∼3.6-fold compared with sham (**Figure 3B**). This CGRP elevation strongly correlated with the magnitude of nuclear NRF2 accumulation (R² = 0.6687; **Figure SF5B**). Region-specific immunofluorescence further confirmed a parallel increase in both CGRP expression and phospho-Ser40-NRF2 in the cortex and hippocampus (**Figure 3C**, **SF5C**), reinforcing the tight association between this neuropeptide and NRF2 activation.

To establish causality, we performed a pharmacological blockade using the anti-CGRP monoclonal antibody Fremanezumab. Pre-treatment with Fremanezumab significantly abolished the DR- and TNS-driven increases in both nuclear NRF2 accumulation and the nuclear-to-cytoplasmic NRF2 ratio (**Figure 3D**, **SF6A**). Furthermore, CGRP blockade significantly attenuated the corresponding elevation in Ser40-phospho-NRF2 (**Figure 3E**, **SF6B**). These data confirm that CGRP is a necessary upstream mediator for the activation of the NRF2 antioxidant response following trigeminal stimulation, solidifying its role as a critical, non-electrophilic effector.

### DR activates the TBI-Critical p62–KEAP1–NRF2 pathway via CGRP signaling

We next sought to assess whether TNS-induced NRF2 activation could proceed via the p62-KEAP1 axis, a non-electrophilic pathway known to be impaired in TBI [9], as was seen in natural DR [16]. This pathway relies on the phosphorylation of the ubiquitin-binding adaptor p62 (SQSTM1), specifically at Ser351, which targets NRF2’s bound inhibitor, KEAP1, for selective autophagic degradation, thereby liberating NRF2 for nuclear translocation [42,43]. To test this, we assayed cortical lysates 30 minutes after a single DR or TNS session. We found that both DR and TNS induced a significant increase in total p62 (∼1.5-fold) and, more critically, its active form, phospho-p62 (Ser351) (∼2.1-fold), leading to a ∼1.4-fold elevation in the phospho-to-total p62 ratio (**Figure 3F-G**). Concurrently, total KEAP1 protein levels were reduced by approximately 30%, consistent with enhanced p62-mediated degradation. Importantly, the thalamus, which illustrated significantly increased NRF2 phosphorylation with TNS, as compared to DR, also exhibited a significant increase in immunofluorescent expression of phospho-p62 (Ser351). To confirm the specificity of this mechanism, we assessed markers of global autophagy. Importantly, phosphorylation at the canonical autophagy-related site p62-Ser403 and the conversion of LC3-I to LC3-II were both unchanged (**Figure SF7**). This indicates that the phosphorylation of p62 following natural DR and TNS is directed toward engagement of the p62–KEAP1 axis for NRF2 activation, rather than toward inducing generalized autophagy.

To link this effect to our upstream mediator, we confirmed the cascade was CGRP-dependent. Pre-treatment with the anti-CGRP antibody Fremanezumab completely abrogated the DR- and TNS-induced increases in total and phospho-p62 (Ser351) and restored KEAP1 to baseline levels (**Figure 3G**, **SF8**). Together, these findings demonstrate that DR and its TNS surrogate utilize a CGRP-dependent signaling cascade to drive targeted p62 phosphorylation and subsequent selective KEAP1 degradation, providing a potent, redox-neutral mechanism to liberate NRF2.

### TBI induces a hyper-acute collapse of the p62-NRF2 axis in peri-infarct tissue, a vulnerability opposed by endogenous CGRP signaling

To characterize the integrity of the NRF2 pathway in the hyper-acute phase of TBI, where oxidative stress rapidly undermines therapeutic efforts [5,11], we analyzed brain tissue at 3 hours post-injury using a CCI model. Our analysis distinguished between the infarct, the vulnerable peri-infarct zone, and the contralateral hemisphere (**Figure 4A**). First, subcellular fractionation at 3 hours post-injury revealed a paradoxical NRF2 response: while the severely damaged infarct core exhibited a stress-induced increase in nuclear NRF2 (1.7-fold vs. sham), the viable peri-infarct tissue suffered a significant loss of nuclear NRF2 (40% reduction), indicating a rapid collapse of the antioxidant response in this critical, salvageable region (**Figure 4B**, **SF9**). Crucially, when we probed the role of endogenous neuroprotection, pre-treatment with the CGRP-antagonist Fremanezumab not only exacerbated the extant loss of activated NRF2 within the peri-infarct, it also induced a *de novo* reduction in nuclear NRF2 within the contralateral region, suggesting a fundamental regulation of the NRF2 pathway by CGRP.

**Figure 4.**
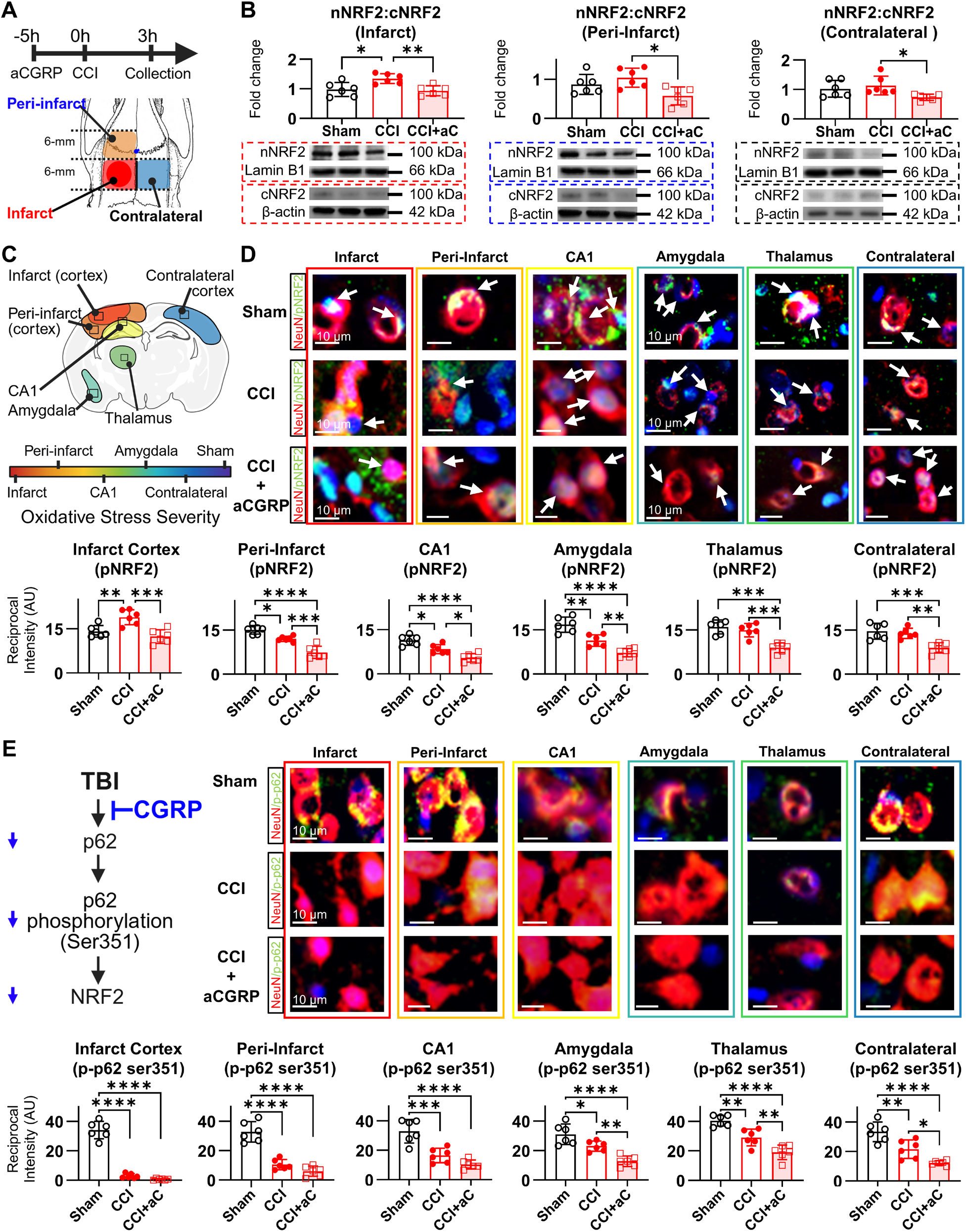
TBI induces a hyper-Acute, CGRP-dependent collapse of the p62-NRF2 antioxidant pathway. **(A)** Schematic of the controlled cortical impact (CCI) model defining the infarct, peri-infarct, and contralateral zones for analysis. **(B)** Immunoblot analysis at 3h post-CCI shows a paradoxical NRF2 response: nuclear NRF2 is increased in the infarct core but significantly depleted in the salvageable peri-infarct zone. **(C)** Region-specific immunofluorescence quantification reveals a widespread loss of activated phospho-NRF2 (ser40) across multiple brain regions following CCI. White arrows point to the approximate center of the neuronal nuclei, which are stained with DAPI (blue). **(D)** Pharmacological blockade of CGRP with Fremanezumab (aC) significantly exacerbates the post-CCI loss of phospho-NRF2, demonstrating a protective role for endogenous CGRP. **(E)** The upstream p62-KEAP1 pathway is also compromised. Quantification at 3h post-CCI shows a significant loss of total p62 and a more profound loss of its activating form, phospho-p62 (ser351). CGRP blockade worsens the depletion of both total and phospho-p62. Data are presented as mean ± SD. Statistical significance for comparisons across all groups was determined by one-way ANOVA followed by Tukey’s multiple comparisons test. Asterisks denote significance relative to the sham group, where \**P* < 0.05, \*\**P* < 0.01, \*\*\**P* < 0.001, and \*\*\*\**P* < 0.0001.

To further map this pathway failure across functionally distinct areas, we quantified the activated form, phospho-NRF2 (Ser40), in six regions stratified by injury severity: the infarct core, peri-infarct zone, hippocampal CA1, thalamus, amygdala, and contralateral cortex. Immunofluorescence confirmed a widespread loss of neuronal phospho-NRF2 post-TBI in all regions barring the infarct core (**Figure 4C-D**, **SF9**). Visual assessment of sub-cellular phospho-NRF2 localization further reveals that, aside from the infarct core and peri-infarct zone, the degree of extranuclear phospho-NRF2 present is decreased. This implies a reactionary increase of the NRF2 import mechanisms in response to the TBI-induced increase of cellular stressors. Crucially, however, pre-treatment with Fremanezumab not only caused an additional 20–30% reduction in phospho-NRF2 accumulation across all assessed regions but also appears to have reduced phospho-NRF2 nuclear import within the infarct core. Contrary to the vehicle, which exhibited an increase in nuclear neuronal phospho-NRF2, animals within the anti-CGRP group exhibited a similar distribution of nuclear and extranuclear phospho-NRF2 as that exhibited by sham. This not only confirms a lack of increase in bio-available NRF2, but also a retardation of the reactionary increase in nuclear import observed within the vehicle group, implying a requirement for the presence of functional CGRP for multiple factors of NRF2 pathway regulation.

Moving upstream to identify the cause, we found this NRF2 activation failure likely stemmed from a collapse of the p62-KEAP1 axis. At 3 hours post-injury, we observed a profound, region-specific loss of total p62 protein. More critically, phosphorylation of p62 at Ser351, the specific modification required to neutralize KEAP1 and liberate NRF2, declined even more precipitously by 65–80% across regions (**Figure 4E**, **SF10**). Consistent with its effect on NRF2, CGRP blockade also aggravated the depletion of both total and phospho-p62(Ser351) by an additional 15–25% (**Figure 4E**). Collectively, these data provide indications that endogenous CGRP provides a critical, albeit insufficient, protective signal that actively opposes the rapid, cascading failure of the entire p62-KEAP1-NRF2 signaling axis in the immediate aftermath of TBI.

### Non-electrophilic DR and TNS show superior efficacy in restoring NRF2 signaling post-TBI

Building on our demonstration of DR’s redox-neutral NRF2 activation in healthy brain, we tested its therapeutic utility in severe CCI–induced TBI. At the 3-hour peak of oxidative stress, rats received one of three interventions: 30-minute natural DR, TNS–induced DR (TNS; 20-s trains every 5 min), or the electrophilic NRF2 activator DMF. Nuclear and cytoplasmic NRF2 levels were then quantified in the infarct core, peri-infarct zone, and contralateral cortex (**Figure 5, SF11**). In the infarct core, CCI alone yielded only a 1.1 ± 0.2-fold increase in nuclear NRF2 without altering the nuclear-to-cytoplasmic ratio. Both DR and DMF induced significant NRF2 nuclear translocation, but DR was superior: DR raised nuclear NRF2 to 2.0 ± 0.3-fold and the nuclear-to-cytoplasmic ratio to 2.5 ± 0.2, whereas DMF achieved only a 1.6 ± 0.2-fold nuclear increase and a ratio of 1.8 ± 0.2. TNS produced a similar but non-significantly greater effect than DR (nuclear 2.2 ± 0.3-fold; ratio 2.7 ± 0.3). The peri-infarct zone, which displayed a 50% loss of cytoplasmic NRF2 and no compensatory nuclear translocation after CCI, was rescued most effectively by DR. DR preserved cytoplasmic NRF2 and drove nuclear levels to 2.8 ± 0.4-fold, outperforming DMF’s 1.8 ± 0.3-fold nuclear increase. Again, TNS showed a consistent trend toward greater NRF2 activation (nuclear 3.0 ± 0.4-fold; ratio 4.5 ± 0.4). Even in the contralateral cortex, DR and DMF induced modest NRF2 nuclear translocation (DR: 1.5 ± 0.2-fold, ratio 1.6 ± 0.2; DMF: 1.4 ± 0.2-fold, ratio 1.5 ± 0.2I), with TNS again trending higher but not significantly so (1.7 ± 0.3-fold; ratio 1.8 ± 0.2). These data demonstrate that, at the tested dosages, non-electrophilic induction of the diving reflex, whether natural or via controlled TNS, delivers a more robust and region-selective NRF2 activation than the conventional electrophilic agent DMF, especially in the peri-infarct zone where therapeutic rescue is most critical.

**Figure 5.**
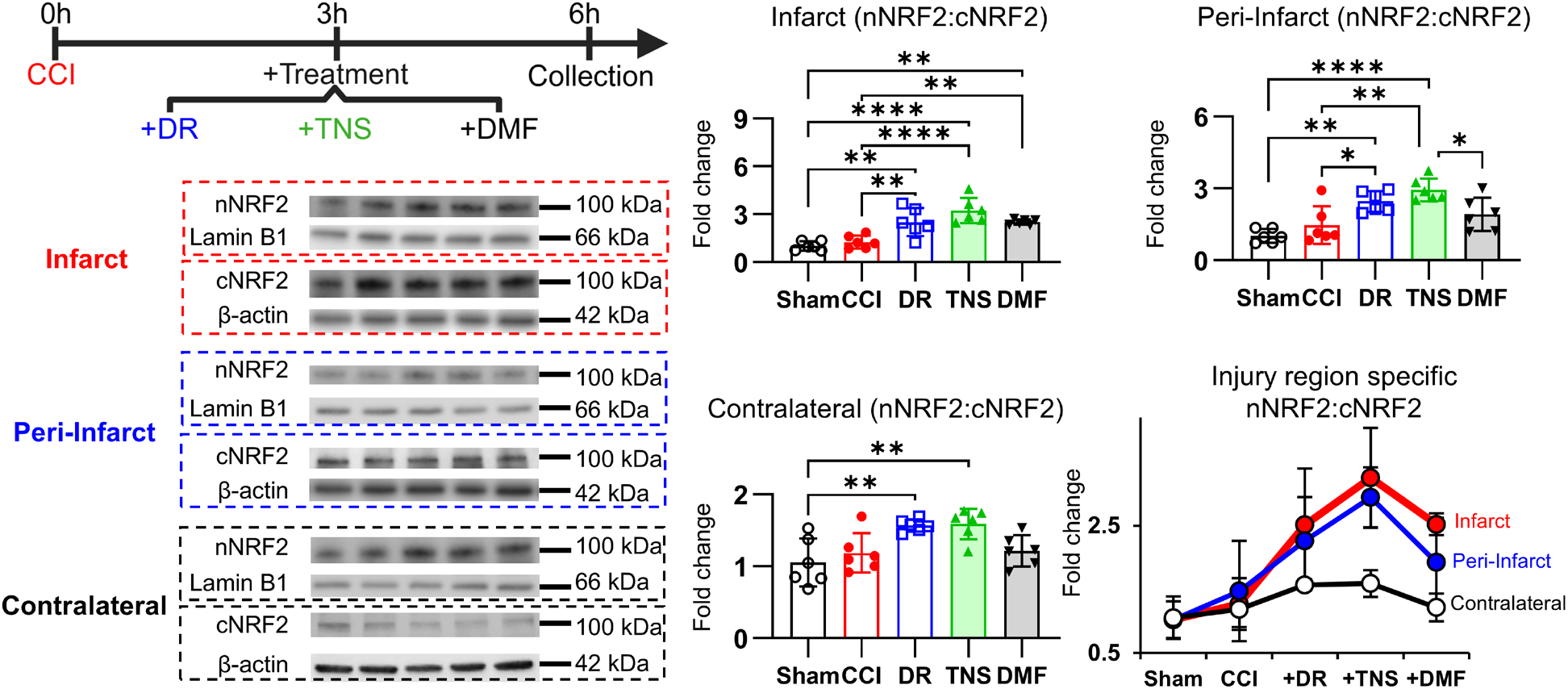
Non-electrophilic natural DR and TNS-induced DR outperform electrophilic DMF in restoring NRF2 signaling post-TBI. Immunoblot quantification of nuclear NRF2 levels in different brain regions 3 hours after TBI and subsequent treatment. Both natural DR and TNS-induced DR robustly increased NRF2 nuclear translocation, significantly outperforming the conventional electrophilic activator DMF. This therapeutic superiority was most pronounced in the salvageable peri-infarct zone, where DR and TNS effectively rescued NRF2 pathway collapse. TNS consistently demonstrated the most potent effect across all regions. Data are presented as mean ± SD. Statistical significance for comparisons across all groups (sham, CCI, CCI+DR, CCI+TNS, CCI+DMF) was determined by one-way ANOVA followed by Tukey’s multiple comparisons test. Asterisks denote significance relative to the sham group, where \**P* < 0.05, \*\**P* < 0.01, \*\*\**P* < 0.001, and \*\*\*\**P* < 0.0001.

### DR drives superior p62-Ser351 phosphorylation and prevents autophagic dysregulation in peri-infarct cortex

To define the upstream mechanisms responsible for the differential NRF2 activation, we quantified key proteins in the p62-KEAP1 axis and general autophagy pathways in peri-infarct cortex harvested 6 hours after CCI (**Figure 6A**, **SF12**). At this 6-hour timepoint, severe TBI alone did not affect KEAP1 or total p62 levels versus sham; however, it decreased phospho-p62(Ser351) to 68 ± 13% of sham, yielding a phospho-to-total p62 ratio of 0.71 ± 0.05 (versus 1.00 ± 0.09 in sham). Treatment with the electrophilic activator DMF decreased total p62 marginally but failed to normalize phospho-p62. In contrast, natural DR significantly lowered KEAP1 to 65 ± 10% of sham, decreased total p62 to 57 ± 7%, yet prevented any decrease in phospho-p62(Ser351). This produced a phospho-to-total p62 ratio of 1.66 ± 0.19, which is 2.3-fold higher than CCI alone and 1.9-fold greater than DMF. TNS recapitulated these effects, trending toward slightly higher phospho-p62 (105 ± 5%) and a ratio of 1.91 ± 0.37. To assess the specificity of this p62 activation, we measured phospho-p62(Ser403), a marker of general autophagy, and LC3-II/I conversion. Severe TBI alone significantly elevated phospho-p62(Ser403) to 273 ± 72% of sham, and DMF treatment exacerbated this to 357% of sham. This was coupled with an increase in the LC3-II/I ratio in both TBI and DMF groups (1.34 ± 0.2 and 1.53 ± 0.2, respectively), indicating a state of dysregulated, excessive general autophagy. In stark contrast, both natural DR and TNS-induced DR prevented this pathological increase in phospho-p62(Ser403) and completely normalized the LC3-II/I ratio. Heatmap analysis visually confirmed that DR and TNS interventions drive signaling toward selective KEAP1 degradation (via p-p62 Ser351), whereas TBI and DMF drive signaling toward generalized autophagy (via p-p62 Ser403 and LC3).

**Figure 6.**
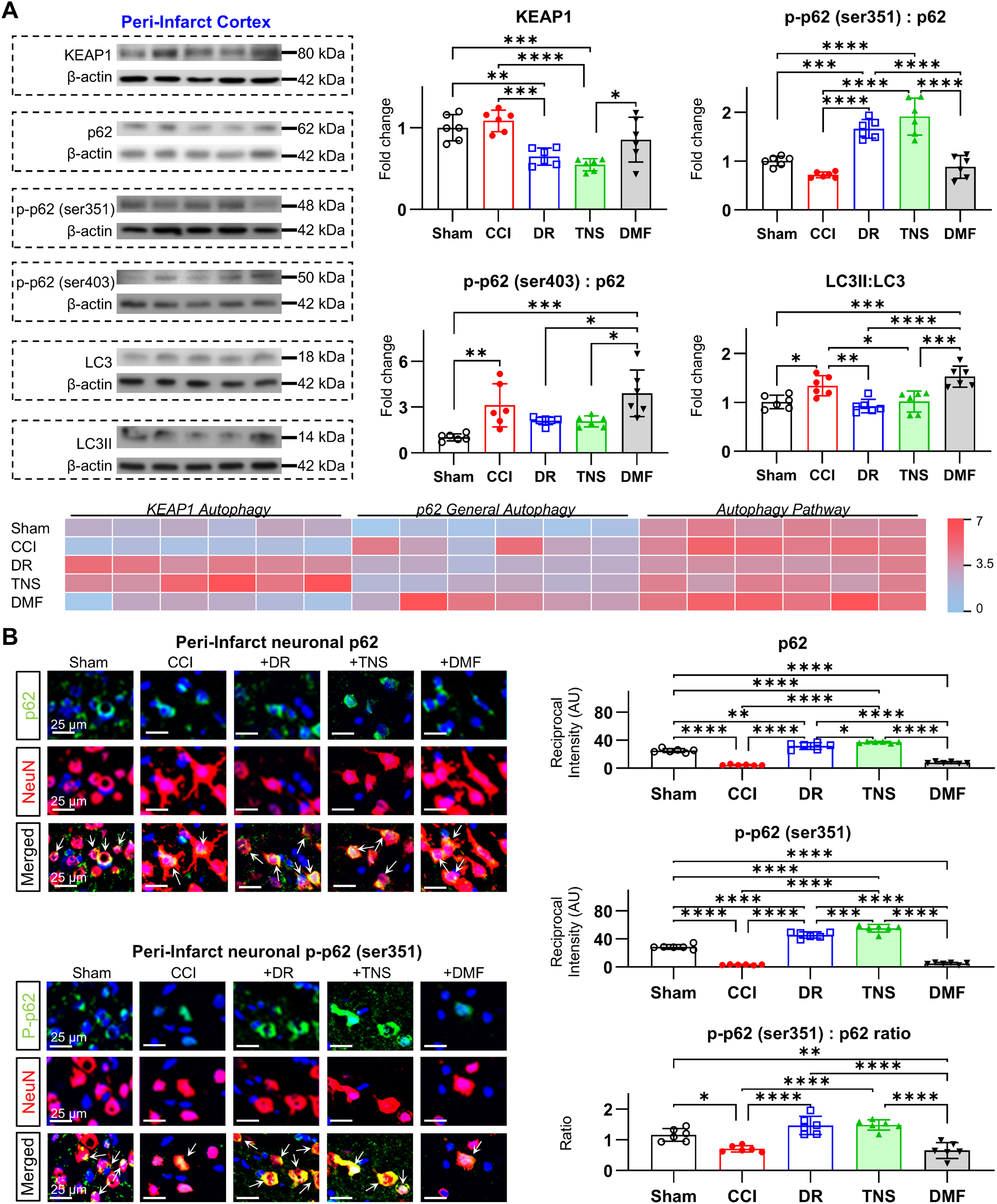
DR and TNS selectively target p62-Ser351 to activate the NRF2 pathway and prevent Post-TBI autophagic dysregulation. **(A)** Western blot analysis of peri-infarct cortex at 6 hours post-CCI. Natural DR and TNS selectively increase the activating phosphorylation of p62 (Ser351), leading to subsequent KEAP1 degradation. Concurrently, DR and TNS prevent the pathological increase in the general autophagy markers phospho-p62(Ser403) and the LC3-II/I ratio, which are elevated following TBI and further exacerbated by the electrophilic activator DMF. A visual heatmap summarizes the divergent signaling, with DR and TNS favoring the NRF2-activating pathway while TBI and DMF trend toward generalized, dysregulated autophagy. **(B)** Immunofluorescence in peri-infarct neurons (NeuN+) confirms these biochemical findings at the cellular level. Representative images and quantification show that DR and TNS restore total p62 levels and, more importantly, significantly augment the activating phospho-p62(Ser351) signal, resulting in a markedly increased phospho-to-total p62 ratio. This effect is not observed with DMF treatment, which exacerbates the loss of phospho-p62(Ser351). Data are presented as mean ± SD. Statistical significance for comparisons across all groups (sham, CCI, CCI+DR, CCI+TNS, CCI+DMF) was determined by one-way ANOVA followed by Tukey’s multiple comparisons test. Asterisks denote significance relative to the sham group, where \**P* < 0.05, \*\**P* < 0.01, \*\*\**P* < 0.001, and \*\*\*\**P* < 0.0001.

Immunofluorescent analysis in NeuN-positive peri-infarct neurons confirmed the importance of phospho-p62(Ser351) for NRF2 activation and tissue protection. CCI reduced neuronal total p62 and phospho-p62(Ser351) immunoreactivity by 82% and 89%, respectively, and DMF exacerbated this phospho-p62(Ser351) loss (**Figure 6B**). In contrast, both DR and TNS not only reversed these deficits but increased total p62 by 25% and phospho-p62(Ser351) by 70% above sham levels, leading to a significantly increased phospho-to-total p62 ratio. These data demonstrate that DR and its TNS surrogate robustly and selectively activate the p62–KEAP1 axis via Ser351 phosphorylation to liberate NRF2, while simultaneously preventing the post-traumatic autophagic dysregulation that is exacerbated by electrophilic activation. This multifaceted mechanism positions DR/TNS as a highly specific and superior therapeutic strategy for the hyper-acute phase of TBI.

### Non-electrophilic DR, TNS, and CGRP restore redox homeostasis and neuronal viability across injury zones

To determine if the non-electrophilic NRF2 activation by DR, TNS, and exogenous CGRP translates into superior redox rescue compared to the electrophilic activator DMF, we first quantified glutathione pools and malondialdehyde (MDA) in the peri-infarct cortex 6 hours after severe CCI. TBI alone halved total GSH, depressed the GSH/GSSG ratio by 60%, and raised MDA by 2.5-fold versus sham (**Figure 7A–B**). While DMF treatment did not normalize these markers, a single 30-minute DR session or TNS regimen fully restored GSH levels and the GSH/GSSG ratio to sham values and reduced MDA accumulation by 60%. Intranasal CGRP recapitulated these restorative effects, providing further support for its role as the upstream mediator of non-electrophilic NRF2 activation. Next, we mapped neuronal phospho-NRF2 (Ser40), lipid peroxidation (4-HNE), and neuronal viability (NeuN immunoreactivity) across four regions stratified by injury severity: the infarct core cortex, the peri-infarct cortex/CA1 region, the amygdala/thalamus region, and the contralateral cortex, using high-resolution immunofluorescence (**Figure 7C–E**, **SF13–16**). DR, TNS, and CGRP each increased phospho-NRF2 and simultaneously reduced 4-HNE in all zones, preserving neuronal integrity to levels statistically indistinguishable from sham controls. By contrast, while DMF also elevated phospho-NRF2, it did not diminish 4-HNE, resulting in continued neuronal loss comparable to that seen in untreated TBI animals. Visual assessment of subcellular localization indicated that DR, TNS, and CGRP all appeared to increase the presence of nuclear phospho-NRF2, an effect not matched by DMF, implicating a fundamental difference in the functional capacity related to bioavailable NRF2 versus its functional nuclear import. Heatmap analysis of the phospho-NRF2-to-4-HNE ratio further illustrates this divergence (**Figure 7F**, **SF17A**); it is clear that, unlike the CCI+DR, CCI+TNS, and, to a lesser extent, the CCI+CGRP groups, the CCI and CCI+DMF groups are distinctly weighted towards the production of oxidative stress without a concomitant balance of phospho-NRF2 activation.

**Figure 7.**
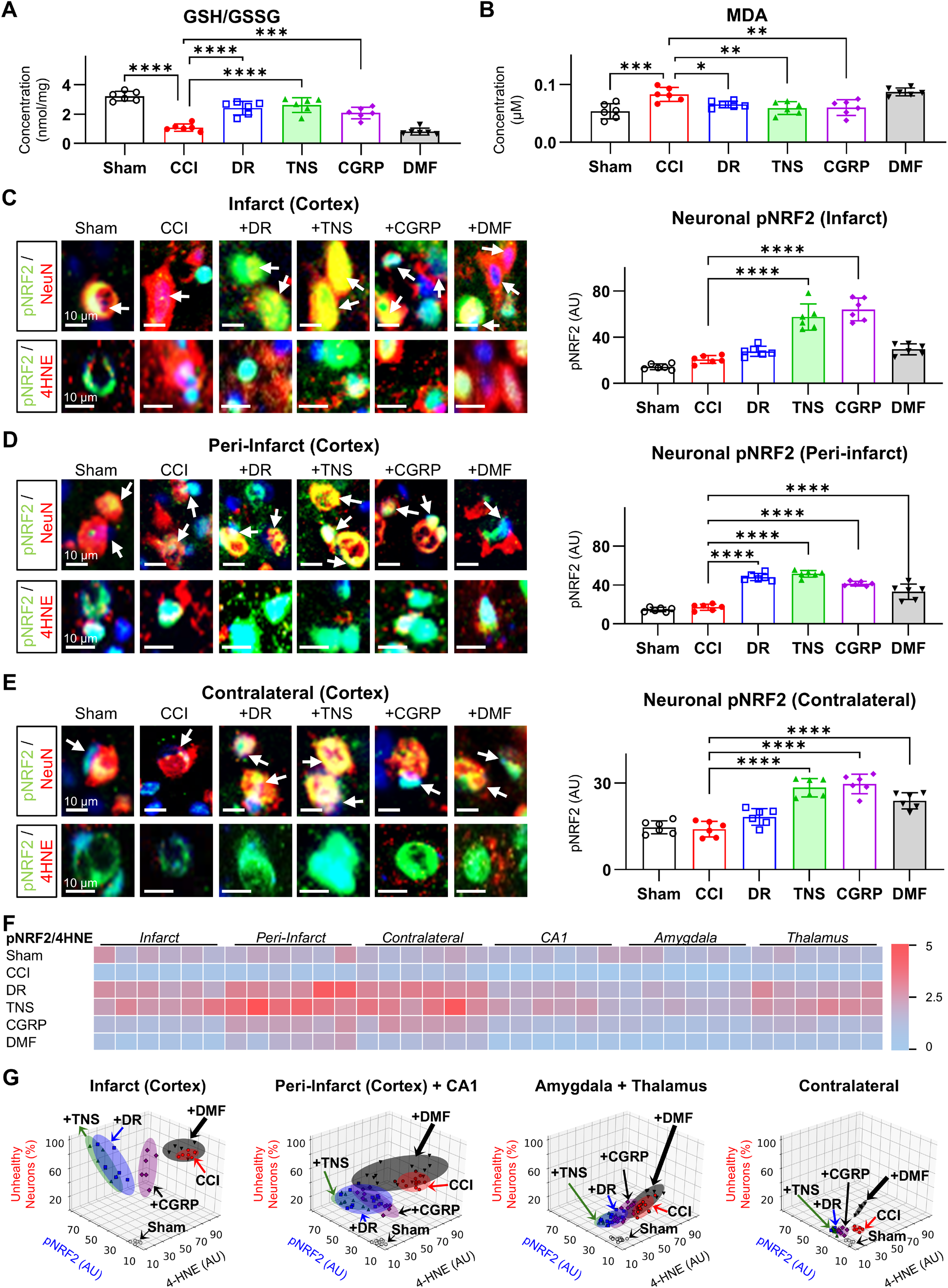
Non-electrophilic DR, TNS, and CGRP restore redox homeostasis and neuronal viability across injury zones. (A-B) Quantification of redox markers in peri-infarct cortex at 6 hours post-CCI. Natural DR, TNS, and exogenous CGRP fully restore the glutathione redox ratio (GSH/GSSG) and significantly reduce malondialdehyde (MDA) levels compared to untreated CCI. DMF treatment does not normalize these markers. **(C-E)** Region-specific immunofluorescence analysis of phospho-NRF2 (pNRF2) and lipid peroxidation (4HNE) in the **(C)** infarct core, **(D)** peri-infarct, and **(E)** contralateral cortex. Non-electrophilic treatments (DR, TNS, CGRP) strongly increase neuronal pNRF2 and concurrently reduce 4-HNE, particularly in the critical peri-infarct region. In contrast, while DMF also increases pNRF2, it fails to suppress 4-HNE accumulation. White arrows in representative images point to neuronal nuclei (DAPI, blue). **(F)** Heatmap analysis of the pNRF2-to-4HNE ratio, illustrating that DR, TNS, and CGRP treatments shift the cellular state toward a protective antioxidant response, whereas the CCI and DMF groups are weighted toward unabated oxidative stress. **(G)** Euclidean distance analysis in the pNRF2 vs. 4HNE feature space, quantifying the overall therapeutic effect relative to the uninjured sham state. The non-electrophilic treatment groups cluster closely with sham, demonstrating a profound restorative effect, while the DMF and untreated CCI groups remain distant, indicating sustained pathology. Data are presented as mean ± SD. Statistical significance was determined by one-way ANOVA followed by Tukey’s post-hoc test for multiple comparisons against the CCI vehicle group. Asterisks denote significance relative to the CCI group, where \**P* < 0.05, \*\**P* < 0.01, \*\*\**P* < 0.001, and \*\*\*\**P* < 0.0001.

To fully confirm that these observed relationships were reliant on CGRP signaling, we administered the CGRP receptor antagonist CGRP_8-37_ concurrently with TNS application. Due to the requirement for constant infusion, this experiment was performed only in the CCI+TNS group and not the CCI+DR group. Pharmacological CGRP blockade inhibited the neuronal p62 activation previously observed with TNS application (**SF18A**). This occurred alongside a diminution of neuronal phospho-NRF2 throughout the brain and an increase in 4-HNE expression relative to TNS alone (**SF18B-C**). The resulting profile trended towards oxidative stress production without NRF2 activation, much like that seen in the CCI vehicle group, confirming the critical role of CGRP in mediating these protective effects (**SF18D**).

Finally, we quantified the overall deviation of each treatment group from the uninjured sham state by computing Euclidean distances in the phospho-NRF2 versus 4-HNE feature space (**Figure 7G**, **SF17B**). In the critical peri-infarct cortex+CA1 region, the DR (ED = 14.3), TNS (17.6), and CGRP (15.3) groups clustered closely with sham animals, whereas the DMF (36.0) and CCI alone (39.7) groups were positioned far apart. A similar pattern emerged in the amygdala+thalamus zone (DR = 10.0; TNS = 13.2; CGRP = 13.4; DMF = 31.4; CCI = 21.3) and even in the ostensibly spared contralateral cortex (DR/TNS/CGRP ED = 10.1–15.6 vs. DMF = 26.4; CCI = 5.9). Importantly, the divergence of the non-electrophilic groups from the sham state was driven almost entirely by a beneficial increase in phospho-NRF2, not by an increase in 4-HNE or a decline in neuronal health. This underscores a fundamental difference between the outcomes of electrophilic versus non-electrophilic NRF2 activation and highlights the superior protective potential of DR, TNS, and CGRP in the hyper-acute TBI environment.

### DR and TNS confer superior acute neuroprotection over electrophilic DMF in TBI

Building on our demonstration that non-electrophilic induction of the DR robustly activates NRF2 in hyper-acute TBI, we compared the efficacy of single-dose DR, TNS-induced DR, exogenous CGRP, and the electrophilic NRF2 activator DMF for their effects on lesion size, NRF2 signaling, and functional outcomes at 24 hours (acute) post-injury. At 24 hours post-CCI, vehicle-treated animals exhibited extensive cortical lesions. DMF had no effect on lesion size, whereas DR reduced acute lesion volume by 23 ± 4%, CGRP by 21 ± 5%, and TNS by 32 ± 3% (**Figure 8A**). To assess NRF2 pathway integrity, we quantified neuronal phospho-NRF2 (Ser40) and correlated it with functional outcomes in region-specific tasks (**Figure 8B**, **SF19-S20**). In the sensorimotor cortex (infarct and peri-infarct), vehicle and DMF groups showed a persistent depletion of phospho-NRF2 at 24 hours (–45 ± 6% and –30 ± 5%, respectively), which correlated with marked sensorimotor deficits (**Figure 8C**, **SF19**). In contrast, DR, TNS, and CGRP restored phospho-NRF2 to levels above sham at 24 hours (+25–40%), preventing sensorimotor impairment. In the hippocampal CA1 memory domain, DR and TNS each drove robust Ser40– NRF2 phosphorylation at 24 hours (+35 ± 7% and +38 ± 6%, respectively), whereas DMF yielded only a marginal +10 ± 5% increase (**Figure 8D**, **SF19**). In the amygdala/thalamus, regions governing anxiety and pain, TNS was the only group to produce a statistically significant improvement in anxiety-like behavior and a corresponding increase in phospho-NRF2 levels relative to vehicle at the 24-hour timepoint, suggesting a unique therapeutic efficacy for the TNS modality (**Figure 8E-F**, **SF19**).

**Figure 8.**
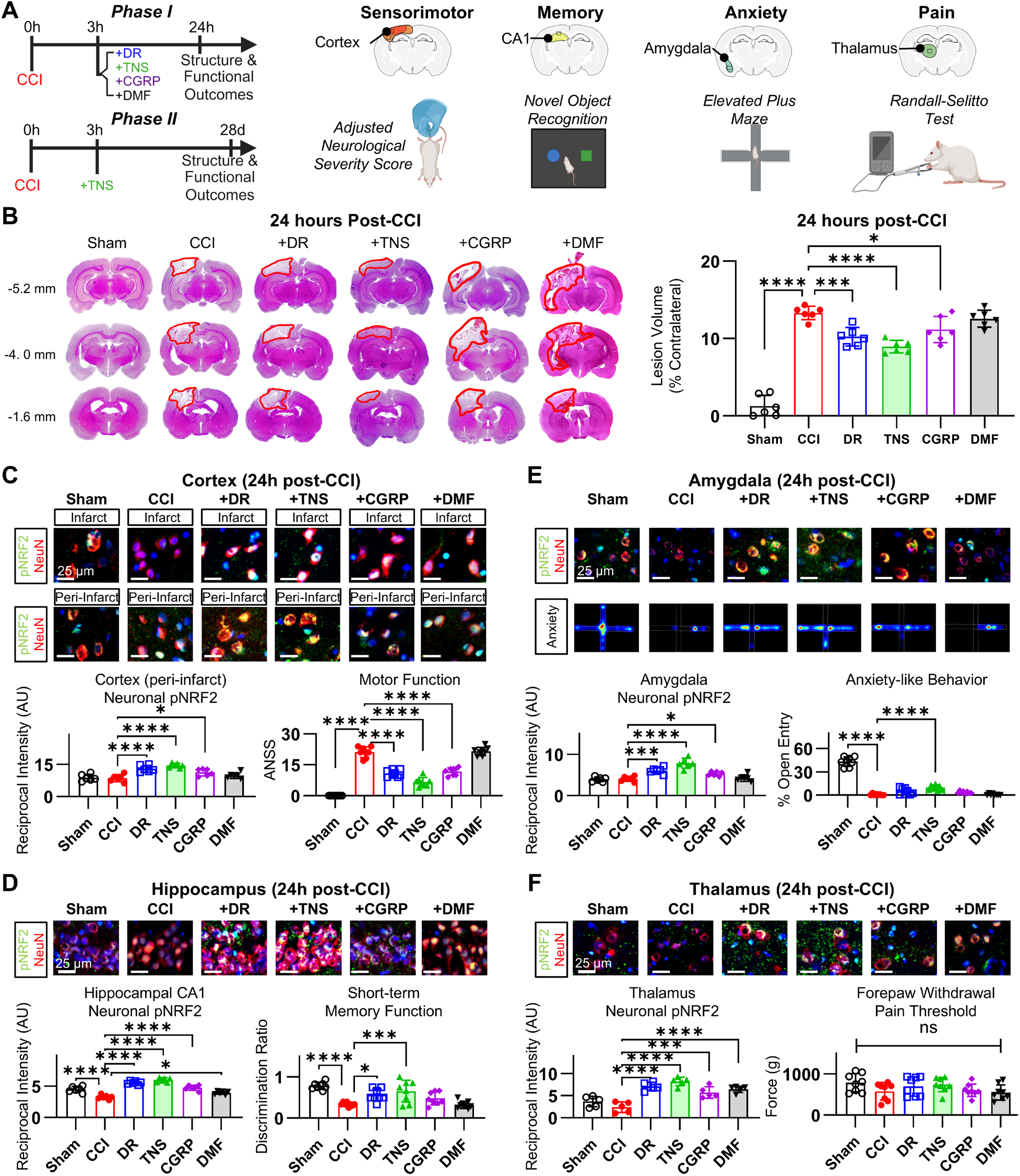
Non-electrophilic DR and TNS confer superior acute neuroprotection over electrophilic DMF in TBI. **(A)** Schematic illustrating the experimental design, correlating regional phospho-NRF2 (pNRF2) levels with their corresponding functional outcomes at 24 hours post-injury. **(B)** Quantification of cortical lesion volume at 24 hours post-CCI. A single post-injury dose of natural DR, TNS, or CGRP significantly reduces lesion size compared to the untreated CCI group, whereas the electrophilic activator DMF shows no significant effect. **(C)** In the sensorimotor cortex (infarct and peri-infarct), DR, TNS, and CGRP restore pNRF2 levels and prevent sensorimotor deficits (ANSS) at 24 hours post-injury. In contrast, CCI and DMF groups show persistent pNRF2 depletion and functional impairment. **(D)** In the hippocampal CA1, DR and TNS drive robust pNRF2 activation at 24 hours, correlating with preserved short-term memory in the novel object recognition test. **(E)** In the amygdala, TNS is the only intervention to produce a statistically significant increase in pNRF2 levels at 24 hours, correlating with reduced anxiety-like behavior in the elevated plus maze. **(F)** In the thalamus, the TNS group exhibits significant pNRF2 activation at 24 hours, though there is no corresponding change in pain threshold at this acute timepoint. Data are presented as mean ± SD. Statistical significance was determined by one-way ANOVA followed by Tukey’s post-hoc test for multiple comparisons against the CCI vehicle group. Asterisks denote significance relative to the CCI group, where \**P* < 0.05, \*\**P* < 0.01, \*\*\**P* < 0.001, and \*\*\*\**P* < 0.0001.

Despite receiving the highest CGRP levels, the CGRP supplementation group exhibited lower NRF2 activation (nuclear-to-cytoplasmic ratio) than DR or TNS, suggesting a non-monotonic dose-response effect for this neuropeptide. This indicates that exogenous CGRP may require fine dose titration to match the dynamic, reflex-driven release pattern for optimal NRF2 engagement. These data collectively demonstrate that non-electrophilic activation of NRF2 via DR or controlled TNS provides superior acute structural and functional preservation in TBI compared with the classical electrophilic activator DMF. Furthermore, the trend towards increased efficacy of TNS, highlighted by the difference in anxiety-related circuit modification, highlights a potential difference in its therapeutic profile compared to both natural DR and pharmacological mimicry at the tested dosages.

### TNS confers sustained chronic neuroprotection in TBI via persistent NRF2 Activation

To determine if the superior acute protection conferred by TNS translates into long-term therapeutic benefits, we assessed lesion volume, NRF2 pathway integrity, and functional outcomes at 28 days post-injury in sham, CCI-vehicle, and CCI-TNS treated animals. At 28 days, vehicle-treated TBI animals exhibited significant chronic pathology, characterized by severe tissue loss in the infarct region and near-complete atrophy of the ipsilateral hippocampus. In stark contrast, a single hyper-acute TNS session produced profound and durable neuroprotection, reducing the chronic lesion volume to 61 ± 6% of vehicle controls (**Figure 9A**). This long-term structural preservation was directly associated with sustained NRF2 pathway activation and corresponding functional recovery. In the peri-infarct sensorimotor cortex, vehicle-treated animals displayed a persistent depletion of phospho-NRF2, which correlated with significant sensorimotor deficits, including a 60 ± 8% increase in beam-walking errors (**Figure 9B**, **SF21**). The TNS-treated group, however, maintained elevated levels of phospho-NRF2 at 28 days, completely preventing these long-term sensorimotor impairments. In the hippocampal CA1 memory domain, analysis of the posterior hippocampus (Bregma −5.2 mm), the only region where CA1 was structurally preserved in vehicle animals, revealed near-complete neuronal loss and a sustained depletion of pNRF2 expression (**Figure 9C**, **SF21**). In contrast, the neurons within the TNS group retained a healthy morphology and pNRF2 levels comparable to sham controls. Similarly, in the amygdala and thalamus, which govern anxiety and pain, TNS treatment restored phospho-NRF2 levels and behavioral performance on the elevated plus maze to near-sham values (**Figure 9D-E**, **SF21**).

**Figure 9.**
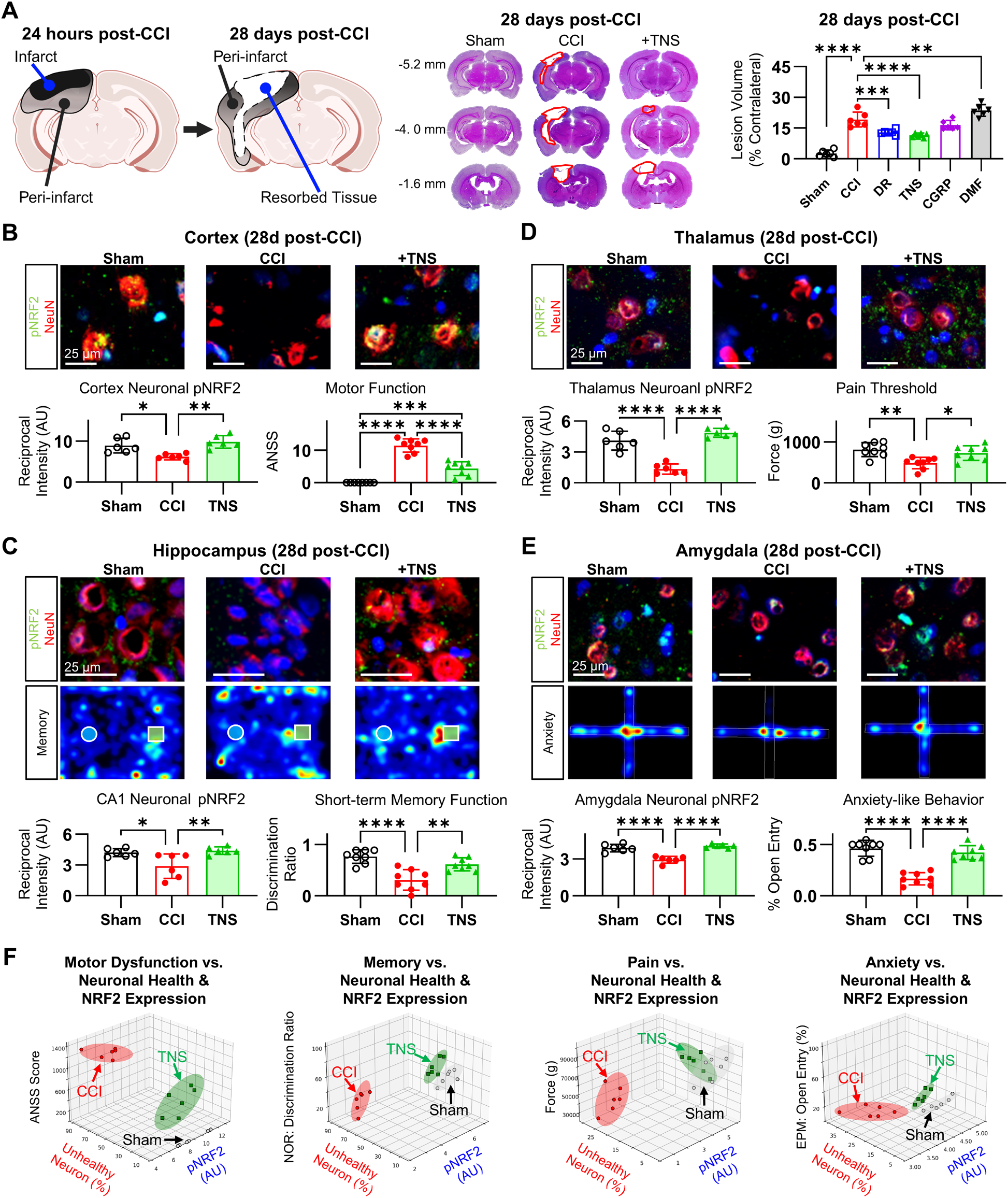
A single hyper-acute TNS intervention confers chronic neuroprotection and sustained NRF2 activation. **(A)** Quantification of chronic lesion volume at 28 days post-CCI reveals that a single TNS treatment administered at 3 hours post-injury significantly reduces long-term tissue loss compared to the untreated CCI group. **(B)** In the peri-infarct sensorimotor cortex, TNS treatment leads to a sustained elevation of phospho-NRF2 (pNRF2) at 28 days, which correlates with a significant prevention of motor dysfunction as measured by beam-walking performance. **(C)** Analysis of the hippocampal CA1 memory domain. By day 28, significant neuronal loss in the untreated CCI group precluded chronic molecular analysis in this region, highlighting the profound neuroprotective efficacy of TNS. In TNS-treated animals, the structural preservation of the CA1 corresponded with retained short-term memory performance. **(D)** In the amygdala, sustained pNRF2 activation at 28 days in TNS-treated animals correlates with reduced anxiety-like behavior in the elevated plus-maze test. **(E)** In the thalamus, preserved pNRF2 signaling at 28 days in the TNS group is associated with improved mechanical pain thresholds. **(F)** Euclidean distance analysis visualizing the overall therapeutic effect. The TNS-treated group clusters closely with the healthy sham group across multiple functional and molecular metrics, while the untreated CCI group remains distant, indicating a near-complete normalization of post-traumatic pathology with TNS. Data are presented as mean ± SD. Statistical significance was determined by one-way ANOVA followed by Tukey’s multiple comparisons test for comparisons across all three groups. Asterisks denote significance relative to the sham group, where \**P* < 0.05, \*\**P* < 0.01, \*\*\**P* < 0.001, and \*\*\*\**P* < 0.0001.

To visualize the overall therapeutic effect, we quantified the deviation of each group from the uninjured sham state by computing Euclidean distances in the phospho-NRF2 versus functional score feature space (**Figure 9F**). Across three of the four assessed regions, the cortex, hippocampus, and thalamus, the TNS group clustered closely with the sham group, indicating a near-complete normalization of both molecular and functional parameters. In fact, in the thalamus, the TNS and sham confidence ellipsoids overlapped. Only the amygdala showed any degree of similarity between the TNS and vehicle groups, an artifact driven more by the relative lack of initial damage in that region than by any retained deficit in the TNS group. These data collectively demonstrate that a single, hyper-acute TNS intervention provides profound and sustained neuroprotection, preserving NRF2 pathway integrity, preventing chronic tissue loss, and restoring multi-domain functional capacity at 28 days post-injury. This highlights the high degree of therapeutic potential that TNS holds as a clinically relevant intervention for mitigating the long-term morbidity of TBI.

## Discussion

This study establishes that DR provides a mechanistically distinct solution for activating NRF2 in the hyper-oxidative environment of TBI. We demonstrate that DR engages the CGRP-p62–KEAP1–NRF2 axis, a non-electrophilic pathway that selectively phosphorylates p62 at Ser351 to drive KEAP1 degradation and liberate NRF2 (**Figure 10**). The therapeutic importance of this redox-neutral mechanism is underscored by its functional consequences: it restores NRF2 signaling across all injury severities, preserves antioxidant reserves, and delivers significant improvements across sensorimotor, cognitive, affective, and pain domains where other multi-target therapies have not succeeded. Furthermore, our findings validate three parallel and clinically viable translational pathways, natural DR via breath-hold immersion, electrically-triggered artificial DR via TNS, and pharmacological mimicry via CGRP. By demonstrating that an evolutionarily conserved reflex can be harnessed to deliver potent, safe, and adaptable neuroprotection, this work provides a new and highly practical strategy for addressing the oxidative crisis at the heart of TBI.

**Figure 10.**
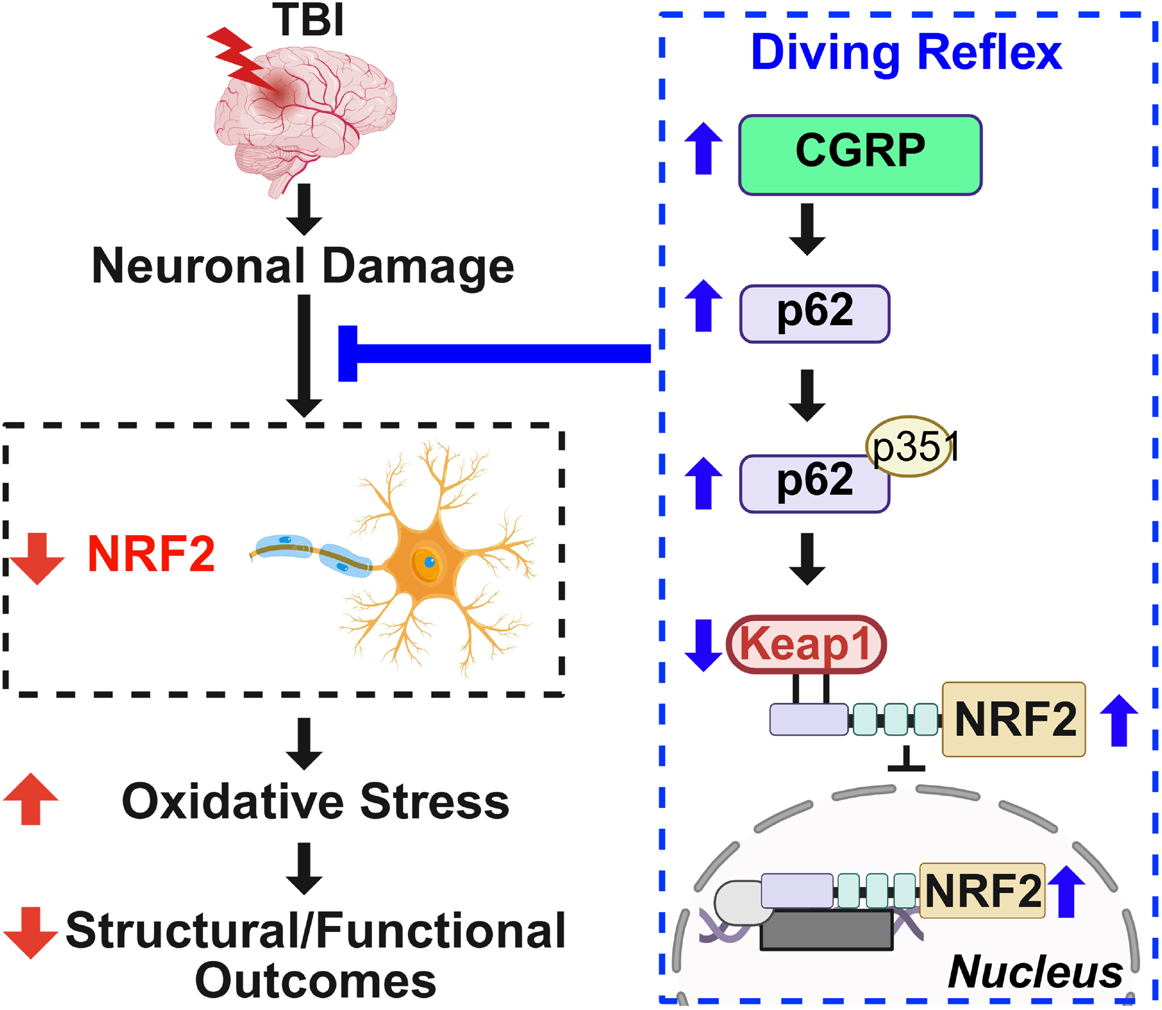
The DR-CGRP-p62-KEAP1-NRF2 axis: a complete pathway for non-electrophilic neuroprotection in TBI. Schematic illustrating the unified therapeutic cascade identified in this study. The diving reflex (DR), whether induced naturally or via its TNS surrogate, stimulates trigeminal afferents, triggering the release of the neuropeptide CGRP. CGRP signaling then engages the p62-KEAP1 axis by promoting both the upregulation of p62 protein and its specific, activating phosphorylation at Ser351. This active phospho-p62 sequesters the inhibitor KEAP1, targeting it for selective autophagic degradation. The removal of KEAP1 liberates NRF2, allowing its phosphorylation at Ser40 and subsequent translocation to the nucleus, where it activates the antioxidant gene program. This entire pathway is redox-neutral, achieving potent neuroprotection by preserving, rather than consuming, cellular antioxidant reserves.

NRF2 orchestrates the cellular response to TBI by simultaneously governing primary pathomechanisms (oxidative stress and neuroinflammation) and the downstream molecular pathways that determine functional outcomes (cognition, affect, and pain), making it a uniquely comprehensive therapeutic target [8,44,45]. Clinical evidence underscores NRF2’s prognostic significance in TBI: decreased 24-hour serum levels predict early mortality [46], while persistent NRF2 suppression correlates with chronic functional deficits, neuropsychiatric sequelae, and Parkinsonism [4,8,47,48]. Our findings confirm this clinical vulnerability at the cellular level, demonstrating that a significant decrease in both nuclear and cytoplasmic NRF2 occurs within 3 hours post-TBI in salvageable peri-infarct tissue, coinciding with the hyperacute oxidative burst that has limited the efficacy of conventional antioxidant therapies [5,11]. A key finding of this study is that a single application of natural or artificially-induced DR at 3 hours post-TBI, well after oxidative damage has been established, was sufficient to achieve robust NRF2 restoration that persisted through the hyperacute, acute, and chronic phases of injury. This delayed therapeutic window is clinically significant, as most TBI patients receive treatment hours after the initial insult. TNS-induced DR specifically restored nuclear NRF2 within neurons, the cells most vulnerable to oxidative stress in TBI [9,49,50], producing a 32% reduction in infarct volume, an effect comparable to or exceeding that reported for multi-drug neuroprotective regimens such as progesterone-nicotinamide [51]. The comprehensive functional improvements observed following DR-mediated NRF2 activation further distinguish it from many existing therapeutic strategies. While combination therapies have often failed to improve behavioral outcomes despite addressing multiple molecular targets [7,52], DR-mediated NRF2 activation simultaneously enhanced sensorimotor function, memory capacity, and ameliorated both anxiety-like behaviors and pain responses. This multi-domain efficacy may reflect a fundamental mechanistic advantage: by activating a single endogenous transcriptional pathway that coordinately regulates a broad spectrum of antioxidant and cytoprotective defense systems, TNS-induced DR achieves a degree of integrated biological protection that is inherently difficult to replicate through the administration of multiple exogenous compounds with potentially uncertain interactions. These findings indicate that TNS-induced DR can effectively engage the NRF2 pathway even after oxidative damage is established, producing both structural neuroprotection and functional recovery across multiple clinical domains.

Despite NRF2’s therapeutic promise, it remains critically underexplored in TBI, with most studies examining canonical electrophilic activators [9]. Our review of all TBI studies employing NRF2 activators whether electrophilic or non-electrophilic reveals fundamental mechanistic differences that may explain variable therapeutic outcomes. Non-electrophilic activators (sinomenine, melatonin, and fucoxanthin) operating through upstream kinases or p62 pathways, demonstrate superior NRF2 induction while consistently preserving mitochondrial function, reducing KEAP1 expression, and suppressing oxidative/inflammatory markers without documented adverse effects [53–55]. In contrast, electrophilic activators can activate NRF2 but produce paradoxical results, including mitochondrial dysfunction, paradoxical leukocyte infiltration, and a retention of inflammatory markers, potentially reflecting their inherent requirement for oxidative stress induction, which has the potential to further exacerbate the TBI-induced oxidative milieu [9,56,57]. The contextual limitations of electrophilic activation are illustrated by DMF, an FDA-approved NRF2 activator successful in low-oxidative multiple sclerosis [58] yet problematic in high-oxidative conditions. DMF exacerbates damage in ischemic stroke [59] and demonstrates inconsistent efficacy in TBI [56,60]. Although these inconsistencies may be due to dosing differences and addressable factors, our brain-wide comparative analysis reveals striking regional patterns that may contribute to translational challenges: DR demonstrated consistent neuroprotection across all brain regions regardless of injury severity, while DMF’s beneficial effects were limited to minimally damaged areas (amygdala, thalamus), with diminished efficacy or potential worsening in severely injured regions. These divergent patterns correlate with distinct biochemical impacts; under the current contexts and given dosage, DMF depleted glutathione and increased lipid peroxidation, whereas DR maintained antioxidant reserves while suppressing oxidative markers. The temporal dynamics reveal why the activation mechanism may determine therapeutic success in TBI. Electrophilic activators present an inherent temporal paradox: DMF depletes glutathione within 60 minutes yet has been demonstrated to require 8-24 hours for transcriptional activation [9,10], perhaps dependent on a relative lack of increase in nuclear import rate, creating a critical vulnerability window during TBI’s hyperacute phase when cellular fate is determined. DR demonstrates fundamentally different kinetics, inducing nuclear NRF2 accumulation (1.6-fold) and phosphorylation (2-30-fold) within 30 minutes, faster than DMF, and substantially outpacing omaveloxolone (∼6 hours), sulforaphane (∼8 hours), or even non-electrophilic compounds like CPUY192018 (∼8 hours) [58,61,62]. Crucially, this rapid activation occurs through mechanisms that build rather than deplete antioxidant capacity when preservation is most critical. These convergent findings suggest that matching activation mechanism and kinetics to TBI’s pathophysiological demands may be essential for therapeutic success, with electrophilic activators potentially requiring careful dosimetric assessment prior to usage in the high oxidative stress milieu of TBI, particularly in the context of severe TBI. DR’s distinctive profile, combining unprecedented activation speed with antioxidant preservation through non-electrophilic pathways, represents a promising therapeutic strategy warranting comprehensive investigation. While these results demonstrate potential mechanistic advantages and encouraging preliminary efficacy, rigorous validation through expanded studies remains necessary. Nevertheless, our data provide strong initial indications that the longstanding challenges in NRF2-based TBI therapeutics may stem from mechanistic misalignment rather than target invalidity. The observed benefits of DR indicate that leveraging endogenous protective pathways with appropriate kinetics and mechanism could enhance therapeutic outcomes, though this promising approach requires rigorous evaluation in larger, controlled studies to establish clinical utility and methodological restrictions.

The p62-KEAP1-NRF2 axis represents a potential molecular bottleneck preventing effective antioxidant responses in TBI, yet this pathway has remained unexplored as a therapeutic target. Our findings demonstrate that DR not only modulates total p62 but, critically, induces selective phosphorylation at Ser351, a post-translational modification that has the capability to fundamentally determine NRF2 activation capacity. Previous investigations in TBI have documented profound p62 dysregulation following a biphasic pattern. The hyperacute phase (1-24 hours) is characterized by rapid p62 decrease in the ipsilateral cortex and hippocampus [13–15], followed by paradoxical accumulation extending to 14 days post-injury [63,64]. As p62 is one of the more primary factors regulating autophagy, this temporal profile suggests an initial autophagic hyperactivation followed by autophagic flux impairment [65]. However, no previous study has examined the deeper phosphorylation status of p62 in TBI, a critical oversight given that phosphorylation site specificity determines whether p62 promotes autophagy or NRF2 activation. The functional dichotomy of p62 phosphorylation has profound therapeutic implications. Ser403 phosphorylation promotes LC3-dependent generalized autophagy, while Ser351 phosphorylation selectively enhances KEAP1 binding through the KIR domain, promoting KEAP1-specific degradation independent of LC3 [65]. Our data reveal that DR specifically drives the latter, whereas DMF treatment does not increase Ser351 phosphorylation and is instead associated with increased Ser403 phosphorylation, suggesting it may engage different cellular stress pathways. The therapeutic importance of this distinction is highlighted by studies in ischemic stroke, where neuroprotection correlates with decreased Ser403 phosphorylation [66], indicating that specifically targeting Ser351 phosphorylation represents a more refined therapeutic strategy for acute brain injury. The potential therapeutic superiority of Ser351 phosphorylation is quantitatively demonstrated by comparative efficacy studies. Interventions achieving p62 phosphorylation at Ser351 produced approximately 2-fold greater NRF2 activation compared to those merely increasing total p62 levels [54,67]. This enhanced efficacy reflects the fundamental transformation in p62 function upon Ser351 phosphorylation: the modified KIR domain achieves approximately 30-fold enhanced KEAP1 binding affinity through additional hydrogen bond formation, matching the affinity of NRF2’s high-affinity ETGE motif [68]. Consequently, phosphorylated p62 both competitively displaces NRF2 from KEAP1 and targets KEAP1 for degradation, dual mechanisms that synergistically restore NRF2 function. While the p62-KEAP1-NRF2 axis has demonstrated therapeutic efficacy in other brain injuries such as intracerebral hemorrhage through ferroptosis inhibition, its targeting in TBI represents an unexplored opportunity. Our demonstration that DR induces Ser351 phosphorylation, absent of Ser403 phosphorylation, begins to address this critical gap, providing the initial evidence that this pathway can be therapeutically accessed in TBI through endogenous mechanisms.

Our findings identify CGRP as a pharmacological mimic of DR that activates NRF2 through a non-electrophilic mechanism fundamentally distinct from canonical activators. CGRP’s peptide molecular structure inherently lacks electrophilic functional groups such as the α,β-unsaturated carbonyl systems required for Michael acceptors that enable cysteine alkylation [69]. While canonical NRF2 activators rely on electrophilic modification of KEAP1 cysteine residues, potentially causing off-target alkylation, antioxidant depletion, and paradoxical pro-oxidant effects, our study demonstrates that CGRP, the key molecular mediator of DR-induced NRF2 activation, has the capacity to achieve NRF2 liberation through p62 phosphorylation at Ser351, thereby preserving cellular redox homeostasis while enhancing the p62-KEAP1-NRF2 axis. Our work demonstrated that DR induces an approximately 3-fold CGRP surge via stimulation of the trigeminal nerve [70,71]. Previous studies establish that CGRP increases total p62 expression [72] while activating PKC and PKA [65,70], which preferentially phosphorylate p62 at Ser351. The observed difference in select phosphorylation form may reflect CGRP-PKA modulation of autophagic signaling. PKA inhibits Foxo activity, prevents LC3 lipidation [73], and phosphorylates ULK1 at inhibitory sites [74]. Since ULK1 drives Ser403 phosphorylation [65], its inhibition would favor Ser351 modification, ensuring NRF2 activation without promoting autophagy, relevant in TBI where autophagic flux is compromised. The translational potential spans multiple modalities: endogenous DR provides physiological CGRP release, electrical trigeminal nerve stimulation enables parametric control, and exogenous CGRP offers an alternative when physical and electrical interventions are impractical. This establishes non-electrophilic NRF2 activation through post-translational modification as a prospective therapeutic strategy. By demonstrating that select p62 phosphorylation effectively activates NRF2 without KEAP1 alkylation, we identify a tenable pathway that maintains cytoprotective benefits while avoiding oxidative stress exacerbation.

Several limitations of this study warrant consideration and guide future research. First, while our DR protocols demonstrate clear efficacy, the parameters used for natural DR, TNS, and CGRP administration represent initial estimations and have not been fully optimized. Systematic investigation into the frequency, duration, and intensity of stimulation is required to define the optimal therapeutic window and dosing regimen. This will also clarify whether the trend towards enhanced efficacy of TNS over natural DR stems from greater consistency or from fundamental differences in the induced biological dose. Second, although a single hyperacute intervention produced remarkable and sustained 28-day benefits, we have not yet explored the potential for repetitive DR activation. It is plausible that repeated TNS sessions during the subacute and chronic phases of recovery could further amplify neuroprotection and functional recovery, a hypothesis that requires dedicated investigation. Third, our mechanistic conclusions, while strongly supported by correlative evidence, require definitive validation using loss-of-function studies. Future work must employ selective inhibitors or genetic knockout models to confirm that the therapeutic effects of DR are causally dependent on the CGRP-p62-NRF2 axis and, specifically, on the phosphorylation of p62 at Ser351. Finally, and most critically, this study’s exclusive use of male animals limits the generalizability of our findings. Given the well-established sexual dimorphism in both CGRP signaling [21] and TBI pathophysiology [45,70], it is essential that future studies assess sex-specific outcomes to inform the safe and effective clinical translation of this therapy. These points, however, represent clear opportunities for refinement rather than fundamental constraints on our conclusions. The core discovery that an endogenous, neuropeptide-driven reflex can safely activate the NRF2 pathway in acute TBI provides a robust and compelling foundation for these future optimization and validation studies.

## Conclusions

This investigation provides evidence that the DR may address the ’safe-vs-potent’ challenge of NRF2 activation in TBI. By engaging a CGRP-mediated p62-Ser351 signaling axis, it achieves significant NRF2 activation without the oxidative cost associated with some conventional electrophilic activators. This redox-sparing mechanism translates into broad therapeutic efficacy, including structural preservation and functional improvements across sensorimotor, cognitive, affective, and pain systems, a systemic response that single-target pharmacology has difficulty replicating. The clinical relevance of this approach is supported by a viable 3-hour therapeutic window that aligns with emergency medicine timelines. Critically, this study also shows that this innate neuroprotective reflex can be reliably induced. We demonstrate that non-invasive TNS provides the key to this control, creating a clinically deployable ‘artificial DR.’ By validating that TNS can convert an involuntary reflex into an on-demand, bedside therapy, this work establishes a tangible strategy for addressing the oxidative crisis not only in TBI, but potentially across a range of acute neurological injuries.

## Acknowledgements

The authors gratefully acknowledge the use of Grammarly, which was employed solely to improve the clarity and grammatical accuracy of the language.

## Author Contributions

CL conceptualized the study, administered the project, acquired the funding, provided the resources, and wrote the original draft of the manuscript. CL, KP, SW, CLD and KR performed experiments, data collection, and data analysis. DLD, AD, JSU and PW contributed to the discussion and interpretation. CLD, DLD, AD, JSU, MB, DS and PW reviewed and edited the manuscript. All authors read and approved the final report and made the decision to submit for publication.

## Data and materials availability

Individual data points are plotted for all graphs. The data that support the findings of this study are available from the corresponding author upon reasonable request.

---

**SF1.**
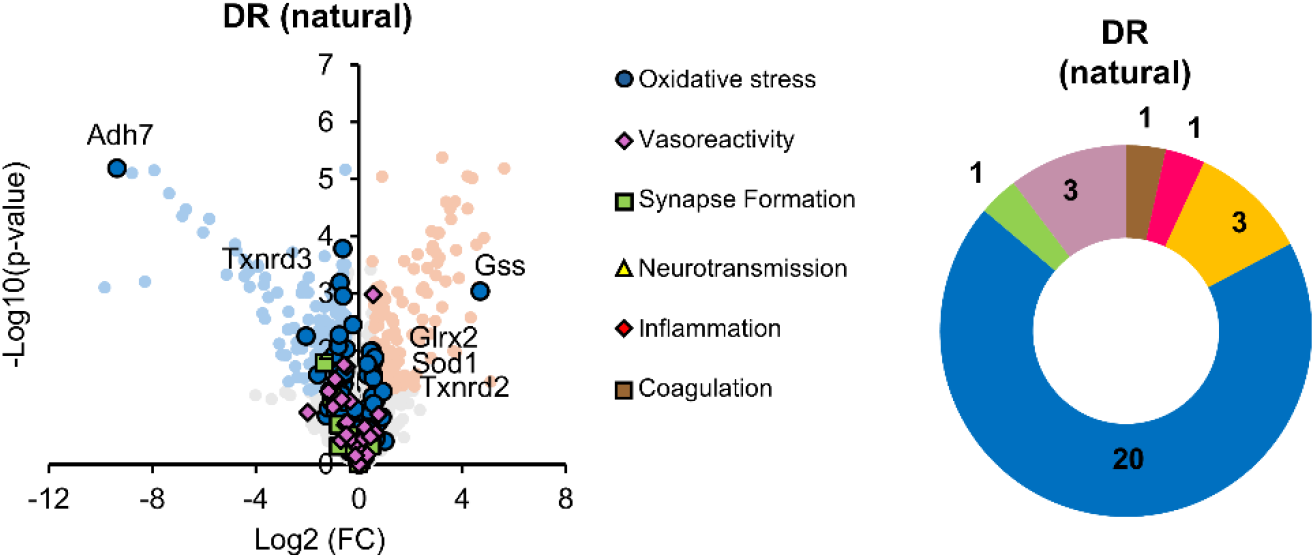
Proteomic signature of the natural diving reflex. Volcano plot illustrating the 485 differentially expressed proteins (DEPs) in the cortex following a single 30-minute natural DR session compared to sham controls. The x-axis represents the log₂(fold-change), and the y-axis represents the −log₁₀(*P*-value). Proteins meeting significance criteria (|log₂ fold-change| ≥ 1 and *P* < 0.05) are highlighted. Note the pronounced upregulation of NRF2-related antioxidant proteins and the concurrent downregulation of classical stress markers, indicating a redox-neutral activation mechanism.

**SF2.**
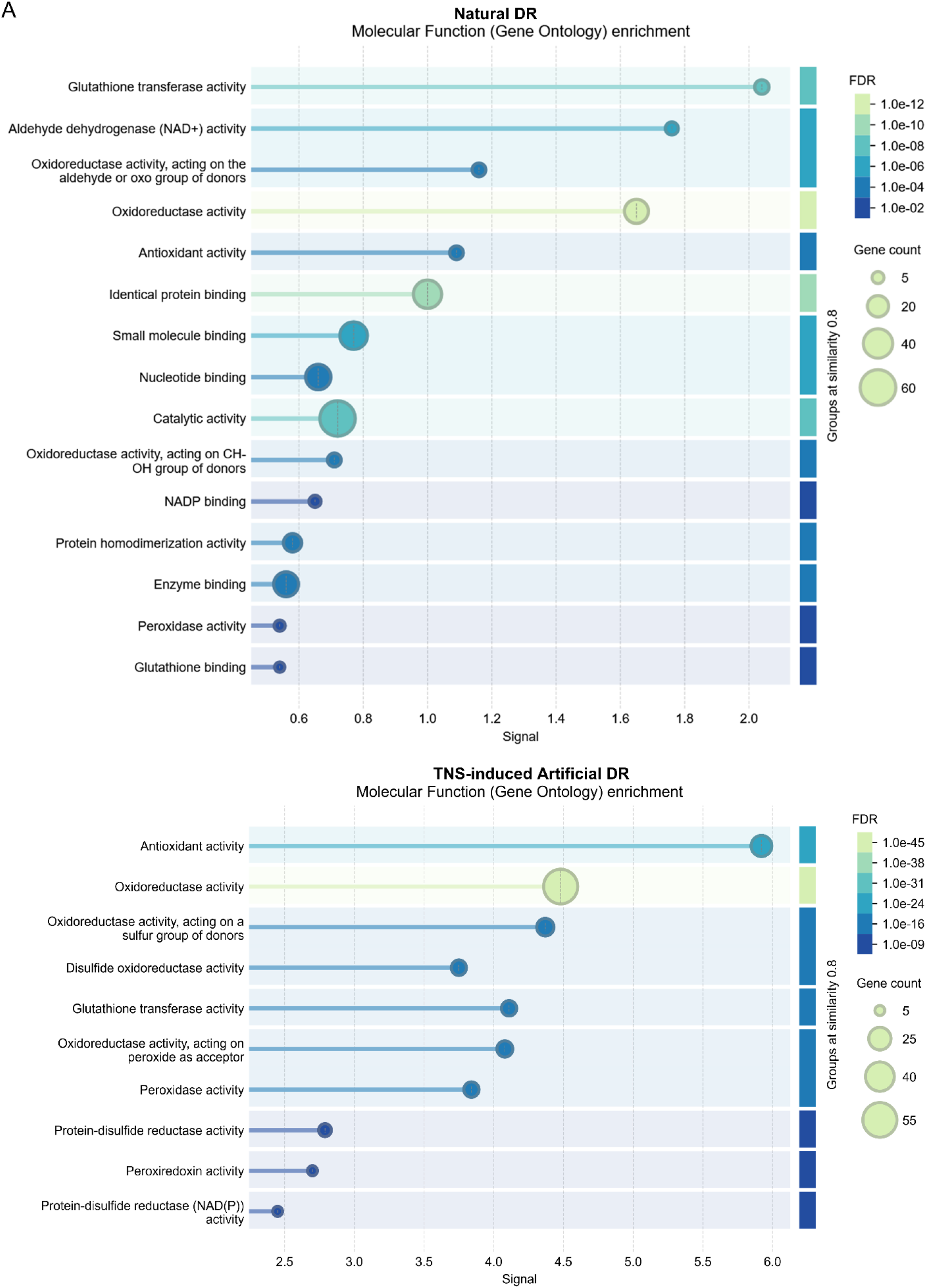
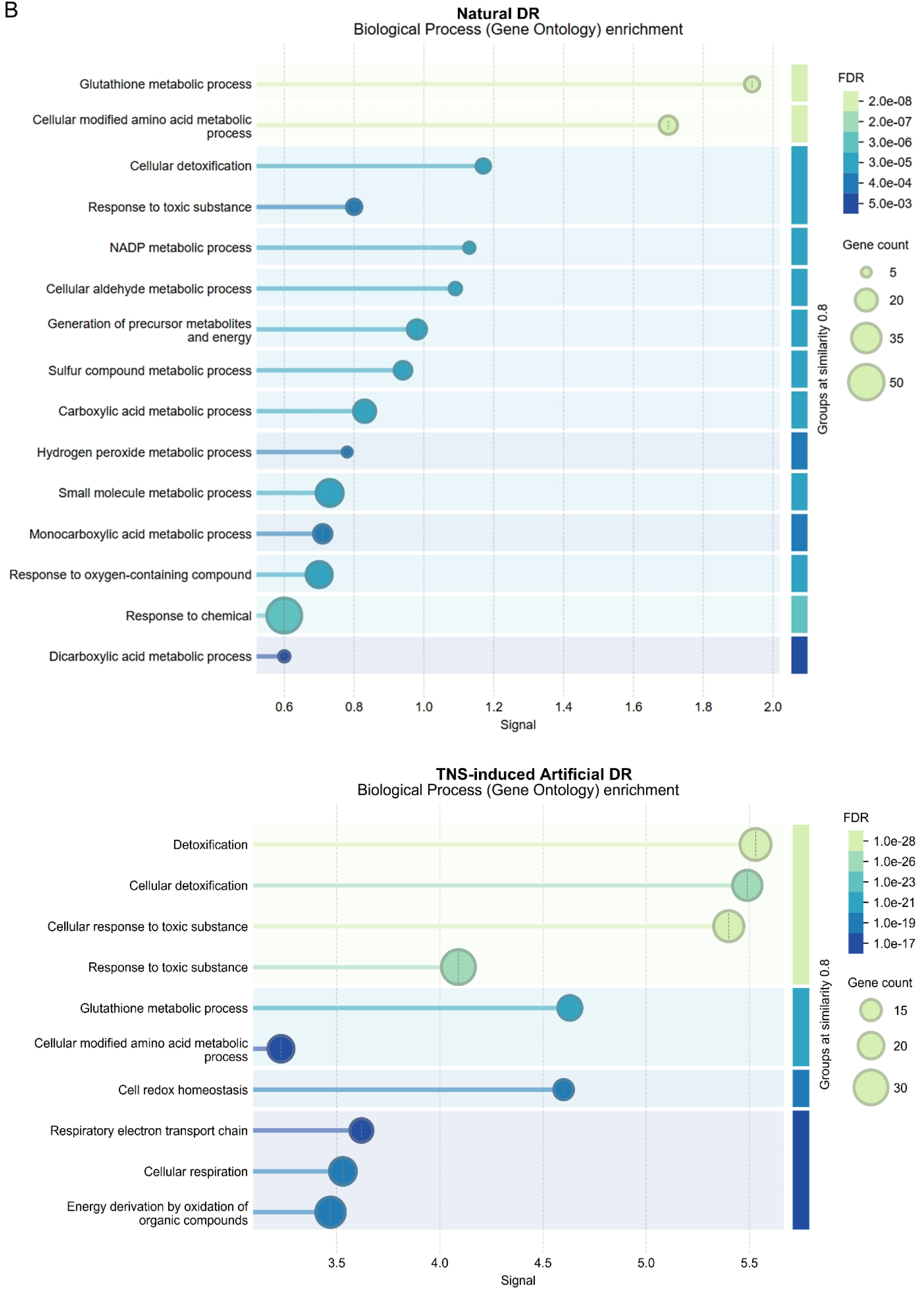
Functional enrichment analysis of proteins modulated by natural DR and TNS-induced artificial DR. Gene Ontology (GO) enrichment analysis of differentially expressed proteins from the cortical proteomics screen. **(A) Molecular function.** The analysis reveals a significant over-representation of proteins involved in antioxidant activity, oxidoreductase activity, and glutathione transferase activity. **(B) Biological process.** The analysis confirms a strong and significant enrichment for pathways related to cellular detoxification, glutathione metabolic processes, and the response to oxidative stress. *Enrichment score on the x-axis indicates statistical significance. Analysis was performed on proteins meeting significance criteria of |log₂ fold-change| ≥ 1 and P < 0.05*.

**SF3.**
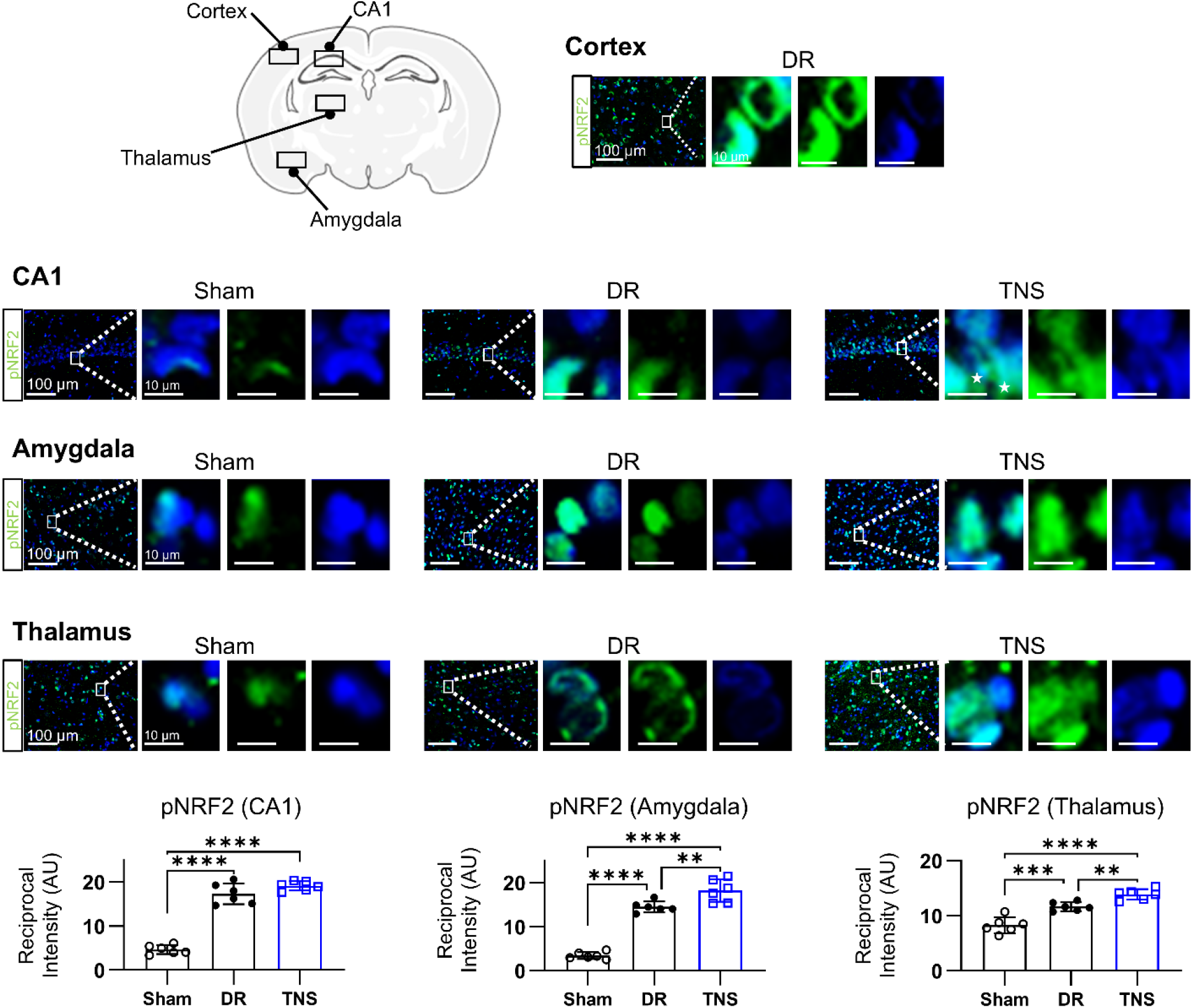
Region-specific NRF2 activation by DR and TNS. Quantification of pNRF2 (Ser40) immunofluorescence across multiple brain regions at 30 minutes post-intervention. Both natural DR and TNS induce significant increases in pNRF2 in the **(A)** hippocampal CA1, **(B)** amygdala, and **(C)** thalamus compared to sham controls. This confirms that NRF2 activation occurs rapidly and across brain regions with varying vulnerability to injury. Data are presented as mean ± SD. Statistical significance was determined by one-way ANOVA followed by Tukey’s post-hoc test. Asterisks indicate significance relative to the sham group, where \**P* < 0.05, \*\**P* < 0.01, \*\*\**P* < 0.001, and \*\*\*\**P < 0.0001*.

**SF4.**
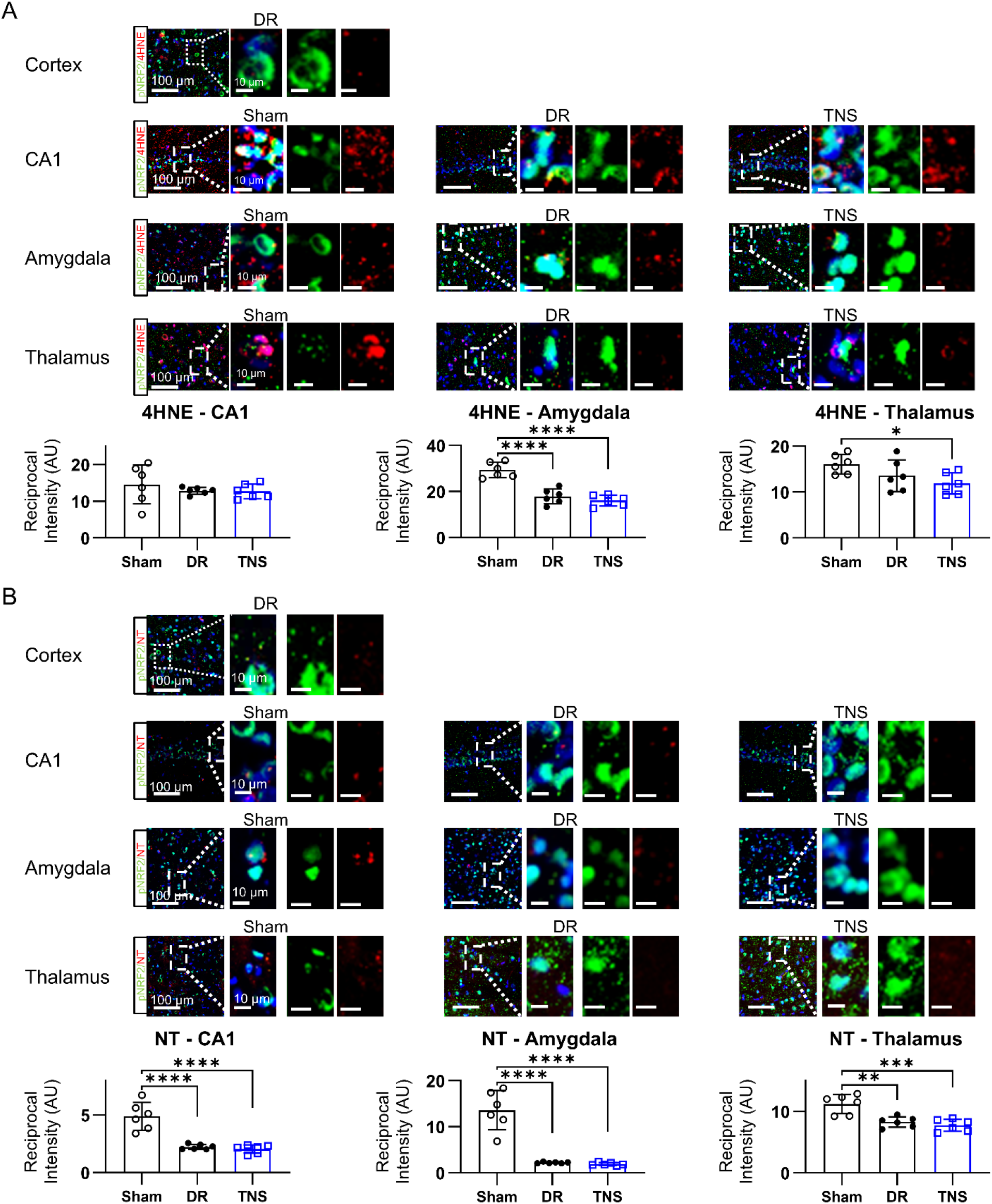
DR and TNS reduce baseline oxidative stress markers across brain regions. Quantification of immunofluorescence for markers of lipid peroxidation (4-hydroxynonenal, 4HNE) and protein oxidation (3-nitrotyrosine, NT) at 30 minutes post-intervention. **(A)** Both natural DR and TNS significantly decrease 4-HNE levels in the dorsal cortex, hippocampal CA1, amygdala, and thalamus. **(B)** Both interventions also significantly reduce NT levels in the dorsal cortex, hippocampal CA1, amygdala, and thalamus. These data confirm a reduction in baseline oxidative stress following both natural and artificial DR. *Data are presented as mean ± SD. Statistical significance was determined by one-way ANOVA with Tukey’s post-hoc test. Asterisks indicate significance relative to the sham group, where \**P* < 0.05, \*\**P* < 0.01, \*\*\**P* < 0.001, and \*\*\*\**P < 0.0001*.

**SF5.**
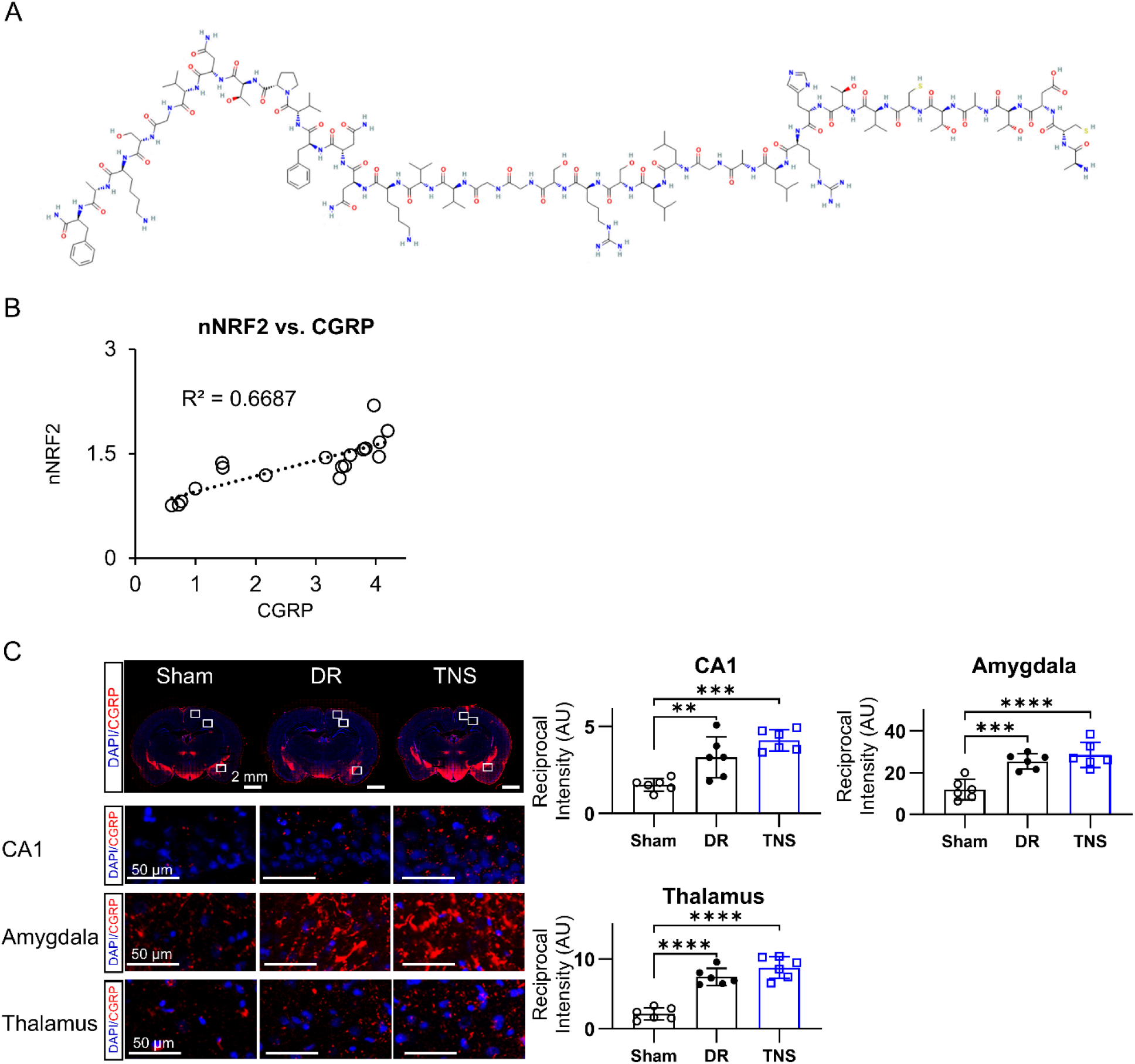
CGRP release and NRF2 activation are spatially and quantitatively correlated. **(A)** Chemical structure of the neuropeptide CGRP, highlighting the absence of electrophile-prone functional groups. **(B)** Correlation analysis showing a strong positive correlation between cortical CGRP concentrations and the magnitude of nuclear NRF2 accumulation. **(C)** Immunofluorescence analysis of CGRP expression across multiple brain regions. Representative images from the cortex show increased CGRP signal following DR and TNS. Quantification confirms a significant increase in CGRP expression in the hippocampal CA1, amygdala, and thalamus in both DR and TNS groups compared to sham. Data are presented as mean ± SD. Statistical significance was determined by one-way ANOVA with Tukey’s post-hoc test. Asterisks indicate significance relative to the sham group, where \**P* < 0.05, \*\**P* < 0.01, \*\*\**P* < 0.001, and \*\*\*\**P < 0.0001*.

**SF6.**
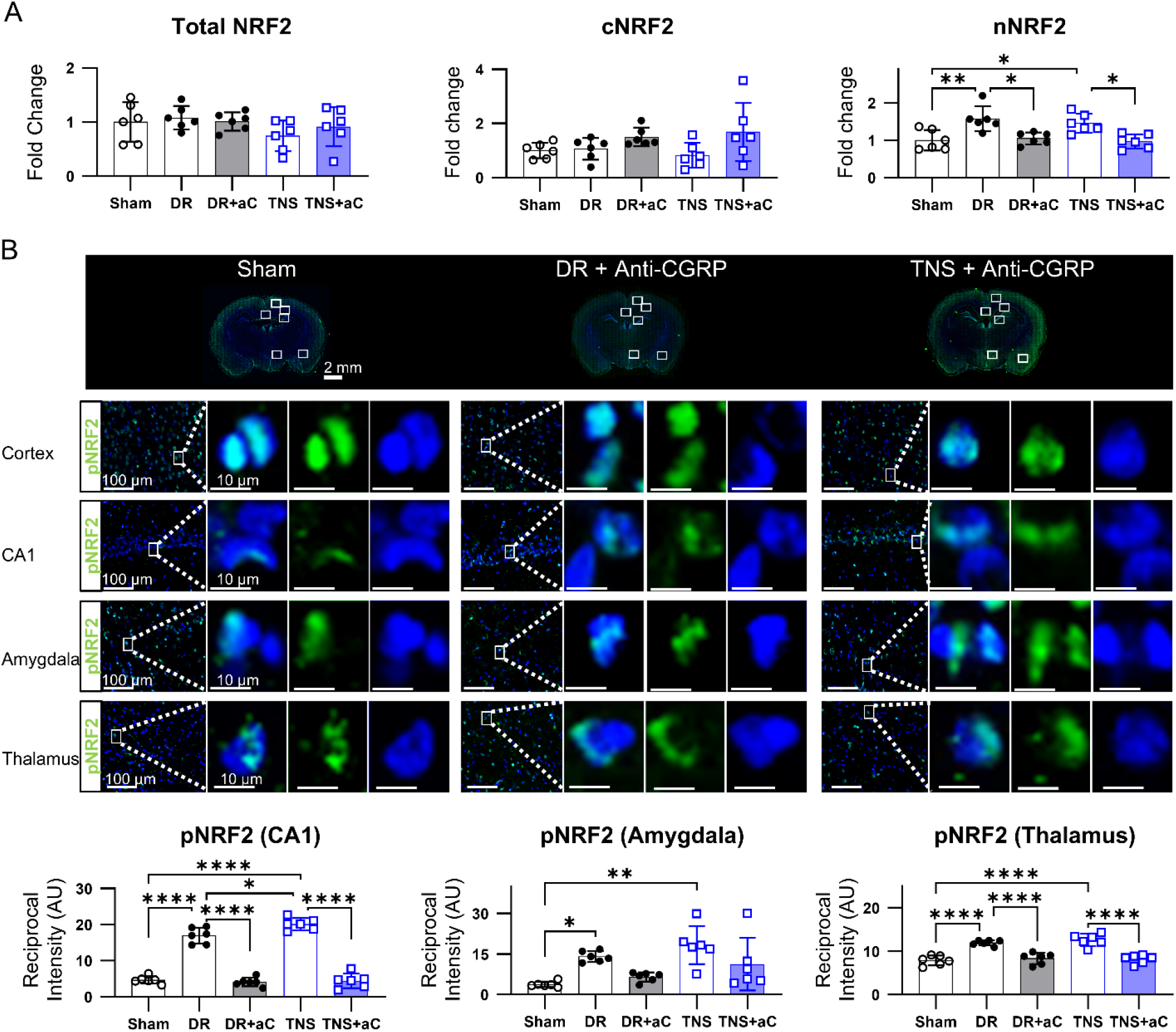
CGRP blockade prevents NRF2 activation in both natural DR and TNS cohorts. **(A)** Pre-treatment with the anti-CGRP antibody Fremanezumab prevents the increase in total, cytoplasmic, and nuclear NRF2 in both natural DR and TNS groups, keeping levels comparable to sham controls. **(B)** Region-specific immunofluorescence analysis of phospho-NRF2 (Ser40) following CGRP blockade. Representative images show that Fremanezumab pre-treatment prevents the increase in pNRF2 signal in both DR and TNS groups. Corresponding quantification confirms that CGRP blockade significantly attenuates the DR- and TNS-induced increase in phospho-NRF2 in the cortex, hippocampal CA1, amygdala, and thalamus. Data are presented as mean ± SD. Statistical significance was determined by one-way ANOVA followed by Tukey’s post-hoc test for multiple comparisons among all groups. Significance is denoted as \**P* < 0.05, \*\**P* < 0.01, \*\*\**P* < 0.001, and \*\*\*\**P < 0.0001*.

**SF7.**
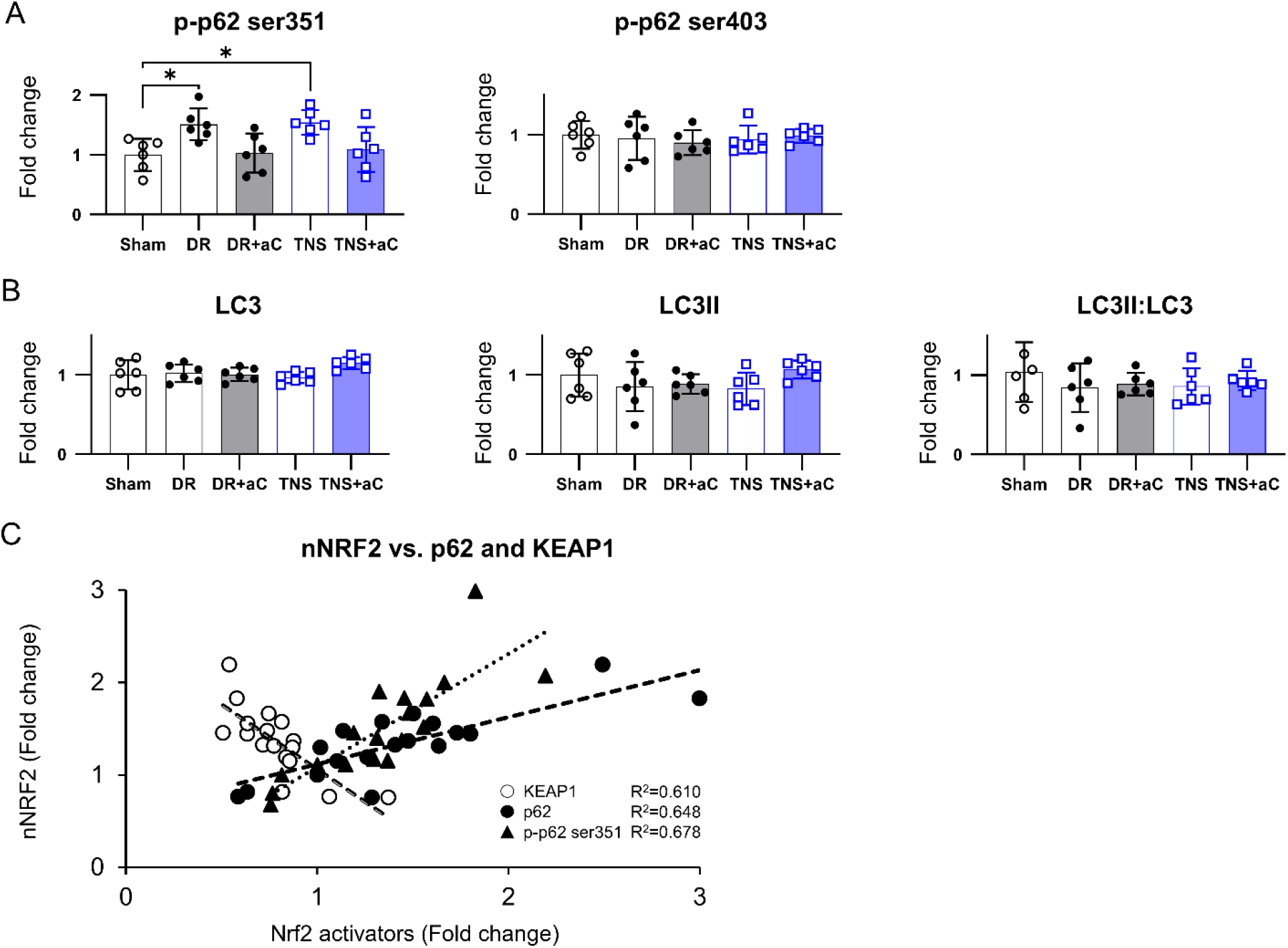
DR and TNS selectively activate the NRF2 pathway via p62-Ser351 phosphorylation, an effect abrogated by CGRP blockade. Analysis of p62 phosphorylation specificity and autophagy markers in cortical lysates 30 minutes post-intervention across five groups: sham, DR, DR + anti-CGRP, TNS, and TNS + anti-CGRP. **(A)** Quantification of phospho-p62 at two distinct sites. Both DR and TNS significantly increase phosphorylation at Ser351 (NRF2-activating) but not at Ser403 (autophagy-related). This effect is completely abolished by pre-treatment with the CGRP antagonist Fremanezumab. **(B)** Quantification of the autophagy markers LC3-I and LC3-II. There are no significant changes in the LC3-II:LC3-I ratio across any of the five experimental groups, confirming the absence of a generalized autophagic response. **(C)** Correlation analysis demonstrating the relationship between specific p62 phosphorylation events and NRF2 activation across the different treatment conditions. Data are presented as mean ± SD. Statistical significance was determined by one-way ANOVA followed by Tukey’s post-hoc test for multiple comparisons among all groups. Significance is denoted as \**P* < 0.05, \*\**P* < 0.01, \*\*\**P* < 0.001, and \*\*\*\**P < 0.0001*.

**SF8.**
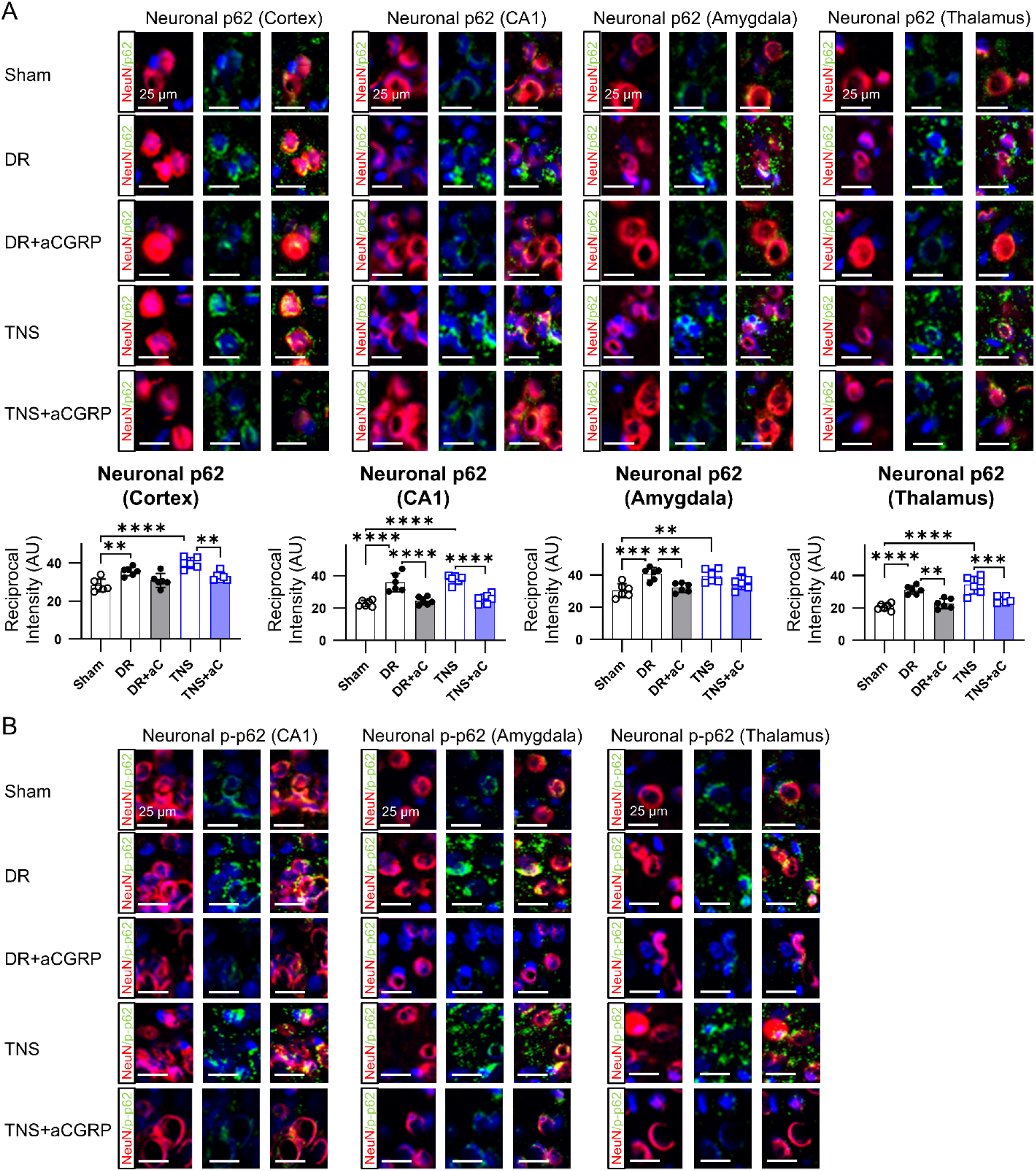
CGRP blockade prevents neuronal p62 activation in both natural DR and TNS cohorts. **(A)** Region-specific immunofluorescence analysis of total neuronal p62. Representative images from the cortex show that both DR and TNS increase p62 signal in NeuN-positive neurons, an effect that is prevented by pre-treatment with the CGRP antagonist Fremanezumab. Corresponding quantification confirms that CGRP blockade abrogates the DR- and TNS-induced increase in total p62 in the cortex, hippocampal CA1, amygdala, and thalamus. **(B)** Region-specific immunofluorescence analysis of the activating form, phospho-p62 (Ser351). Representative images demonstrate that Fremanezumab pre-treatment prevents the increase in neuronal p-p62(Ser351) signal in the hippocampal CA1, amygdala, and thalamus. in both DR and TNS groups. Data are presented as mean ± SD. Statistical significance was determined by one-way ANOVA followed by Tukey’s post-hoc test for multiple comparisons among all groups. Significance is denoted as \**P* < 0.05, \*\**P* < 0.01, \*\*\**P* < 0.001, and \*\*\*\**P < 0.0001*.

**SF9.**
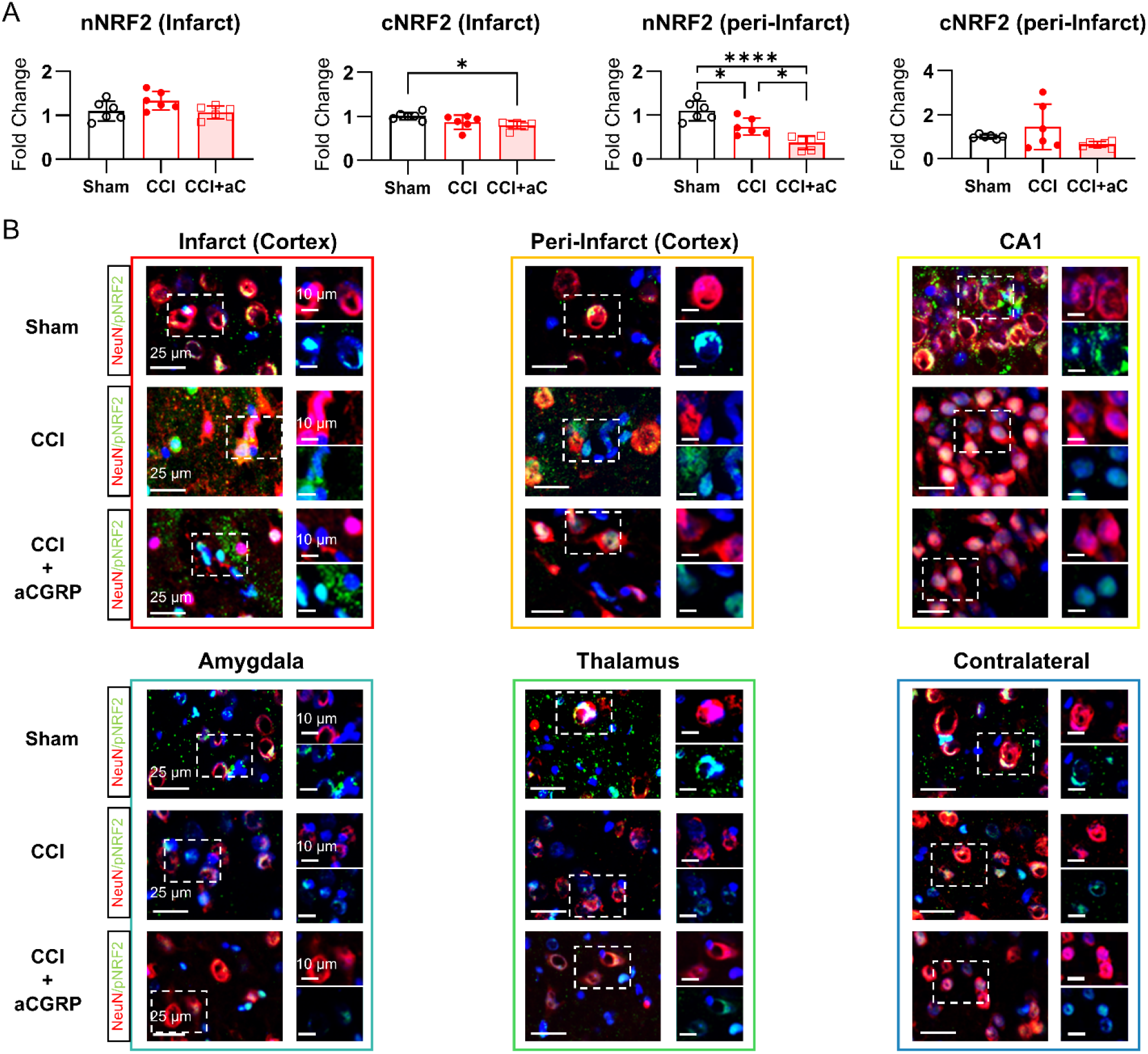
Region-specific NRF2 dysregulation at 3 hours post-TBI is exacerbated by CGRP blockade. **(A)** Quantification of nuclear (nNRF2) and cytoplasmic (cNRF2) NRF2 protein levels in the infarct and peri-infarct cortex at 3 hours post-TBI. While the infarct shows a stress-induced increase in nNRF2, the peri-infarct exhibits a significant loss. Pre-treatment with a CGRP antagonist (aCGRP) exacerbates the loss of nNRF2 in the peri-infarct zone. **(B)** Representative immunofluorescence images of neuronal phospho-NRF2 (pNRF2; Ser40) across six distinct brain regions (infarct cortex, peri-infarct cortex, CA1, amygdala, thalamus, and contralateral cortex) in sham, CCI, and CCI+aCGRP groups. The images visually confirm the widespread loss of pNRF2 and the exacerbating effect of CGRP blockade.

**SF10.**
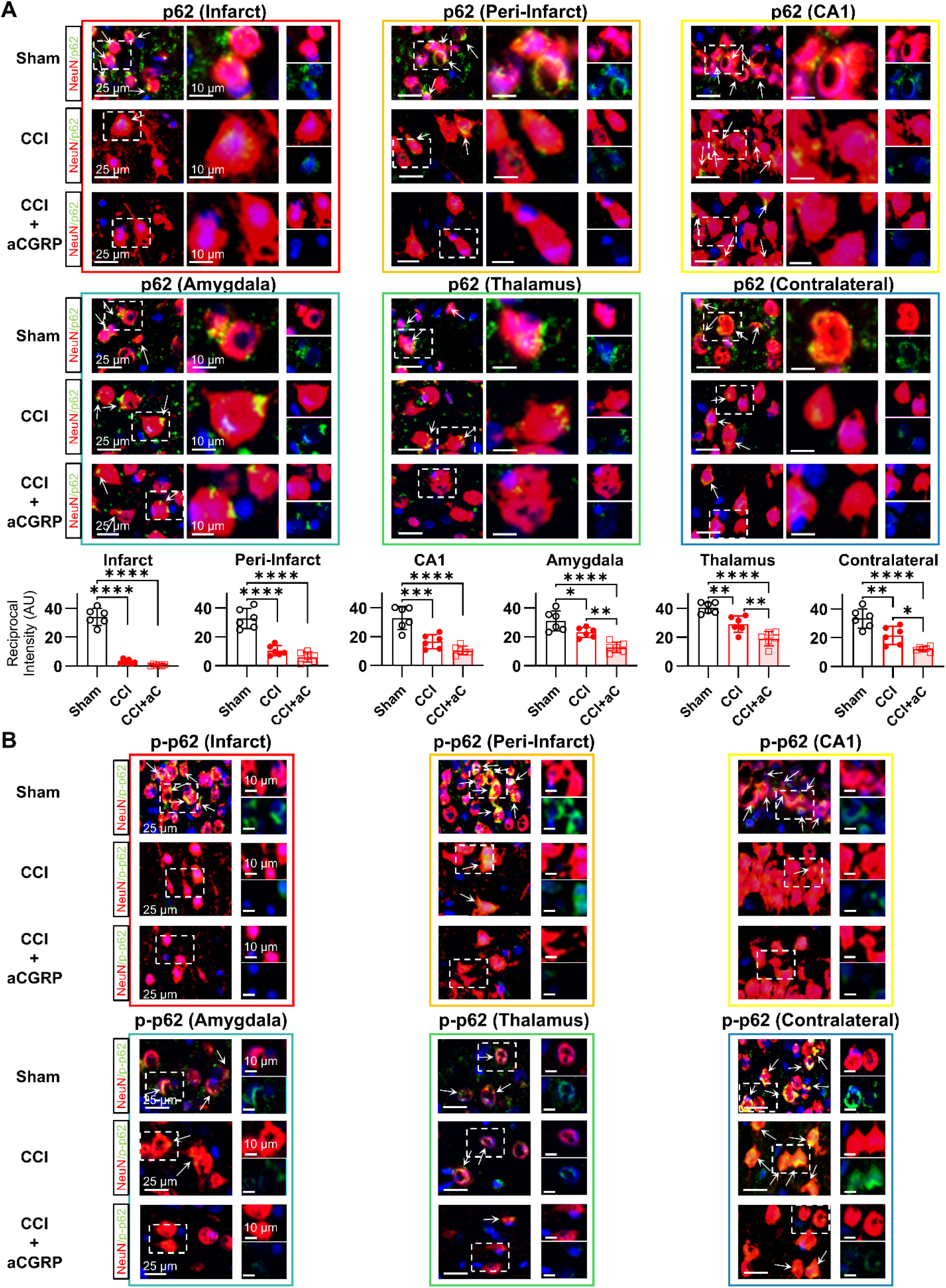
Region-specific p62 dysregulation at 3 hours post-TBI. **(A)** Representative immunofluorescence images and corresponding quantification of total p62 expression in six brain regions across sham, CCI, and CCI+aCGRP groups. The data show a region-specific loss of total p62 protein at 3 hours post-injury, a deficit that is worsened by CGRP blockade. **(B)** Representative immunofluorescence images of the activating phospho-p62 (Ser351) in the same six brain regions. The decline in p-p62(Ser351) is even more precipitous than the loss of total p62, indicating a severe failure of the upstream NRF2 activation pathway. CGRP blockade further aggravates this loss. Data are presented as mean ± SD. Statistical significance was determined by one-way ANOVA followed by Tukey’s post-hoc test for multiple comparisons among all groups. Significance is denoted as \**P* < 0.05, \*\**P* < 0.01, \*\*\**P* < 0.001, and \*\*\*\**P < 0.0001*.

**SF11.**
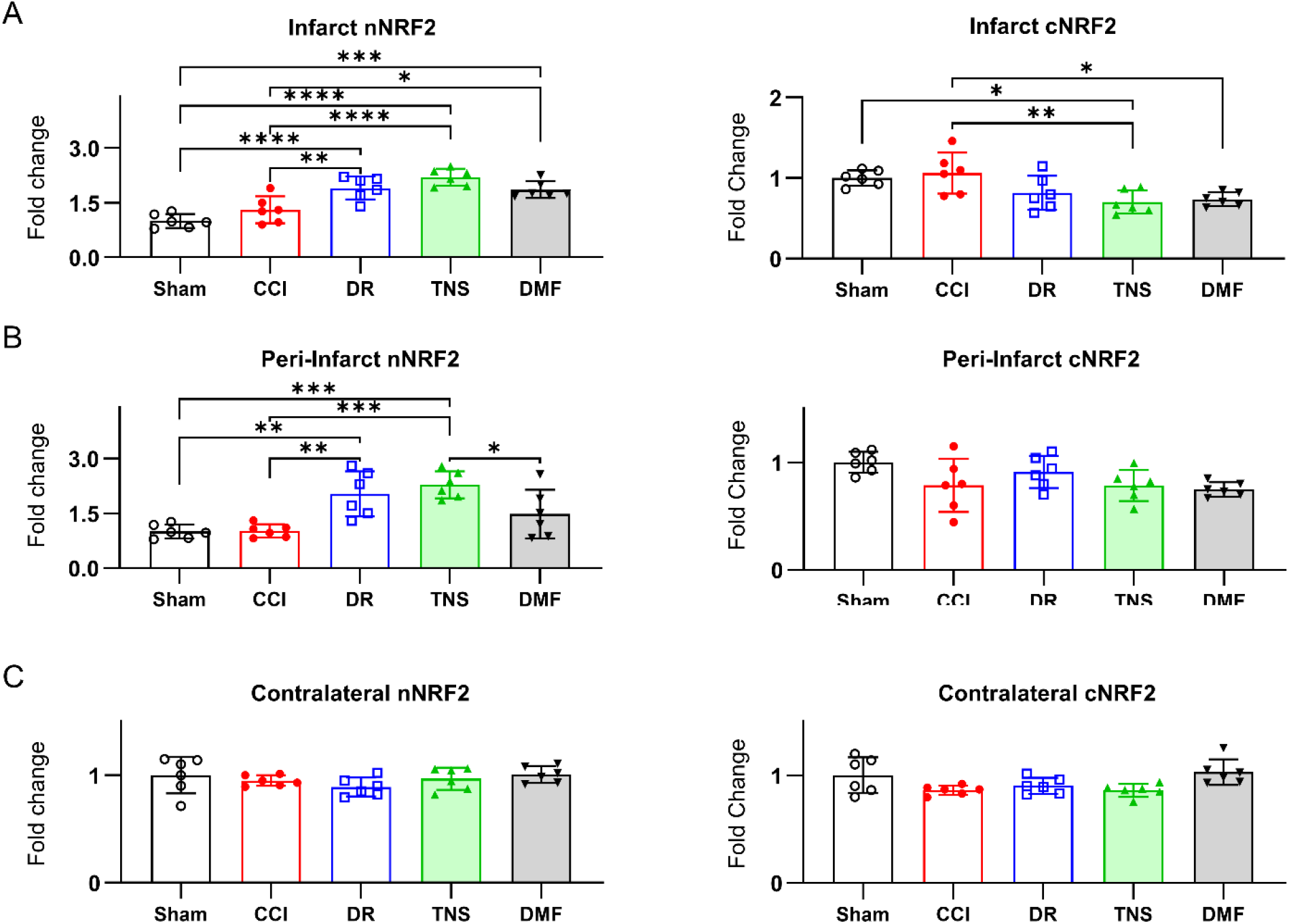
Region-specific NRF2 translocation after DR, TNS, and DMF treatment post-TBI. Quantification of nuclear (nNRF2) and cytoplasmic (cNRF2) NRF2 protein levels at 3 hours post-CCI and subsequent treatment. Five experimental groups are shown: sham, CCI, CCI+DR, CCI+TNS, and CCI+DMF. **(A)** Infarct cortex. **(B)** Peri-infarct cortex. **(C)** Contralateral cortex. Data are presented as mean ± SD. Statistical significance was determined by one-way ANOVA followed by Tukey’s post-hoc test for multiple comparisons among all groups. Significance is denoted as \**P* < 0.05, \*\**P* < 0.01, \*\*\**P* < 0.001, and \*\*\*\**P < 0.0001*.

**SF12.**
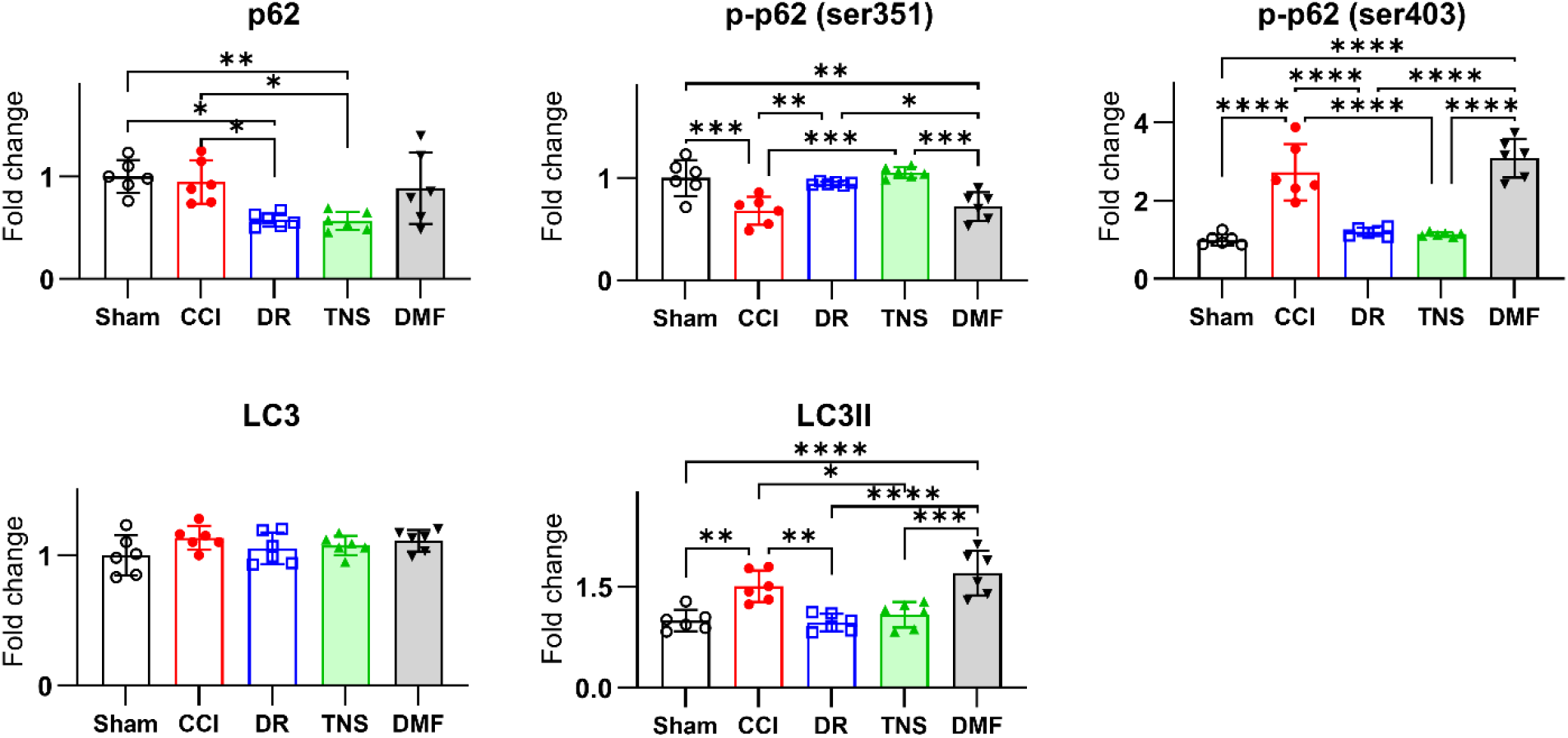
DR and TNS selectively target the p62-KEAP1-NRF2 pathway without inducing generalized autophagy. Detailed quantification of key signaling proteins in the peri-infarct cortex at 6 hours post-CCI across five experimental groups: sham, CCI, CCI+DR, CCI+TNS, and CCI+DMF. The graphs display protein levels of total p62, the NRF2-activating form phospho-p62 (Ser351), the autophagy-related form phospho-p62 (Ser403), and the autophagy markers LC3-I and LC3-II. Data are presented as mean ± SD. Statistical significance was determined by one-way ANOVA followed by Tukey’s post-hoc test for multiple comparisons among all groups. Significance is denoted as \**P* < 0.05, \*\**P* < 0.01, \*\*\**P* < 0.001, and \*\*\*\**P < 0.0001*.

**SF13.**
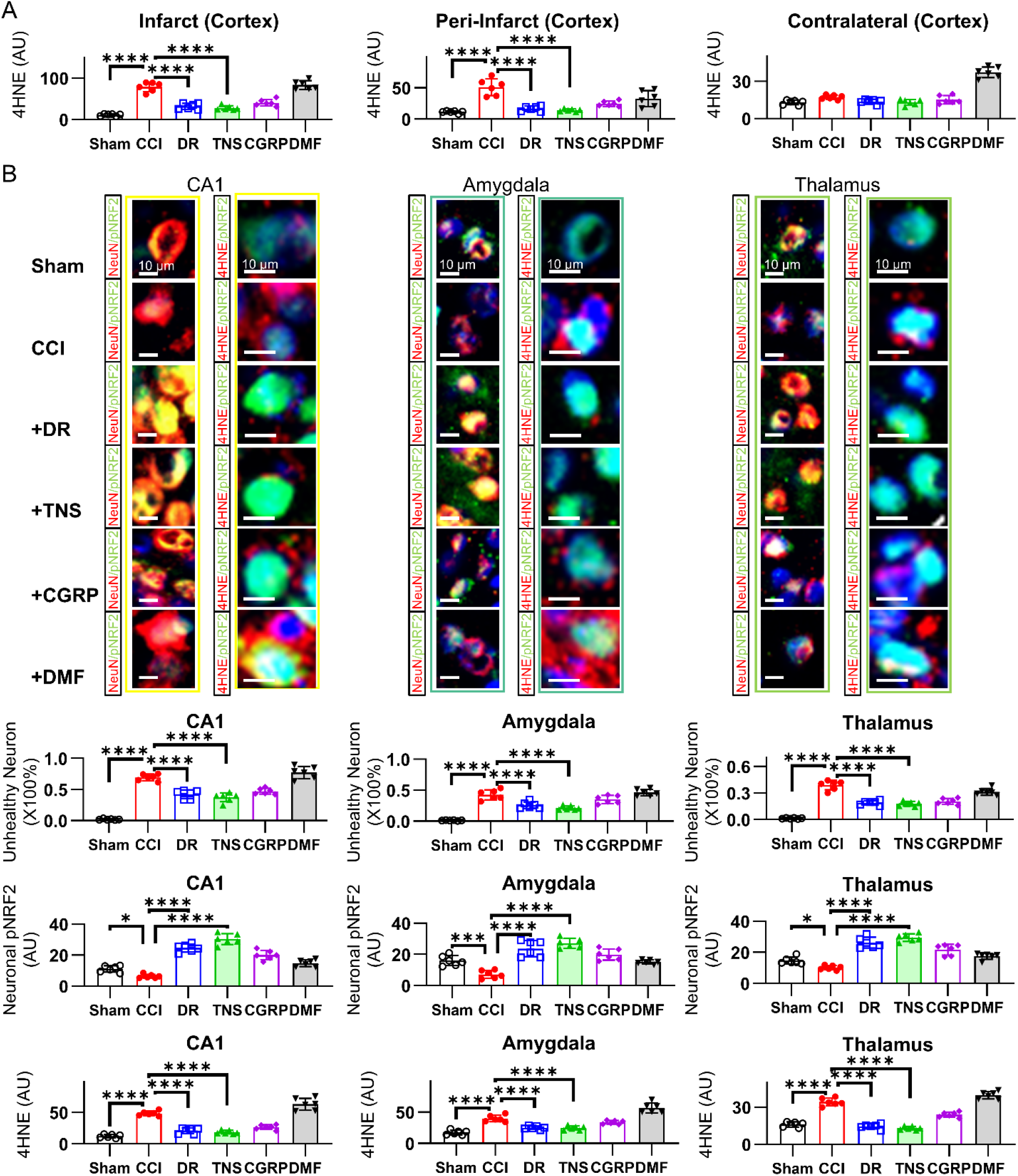
Region-specific analysis of oxidative stress and NRF2 activation post-TBI. **(A)** Quantification of lipid peroxidation (4-HNE) immunofluorescence in the infarct, peri-infarct, and contralateral cortex. **(B)** Detailed immunofluorescence analysis in the hippocampal CA1, amygdala, and thalamus. Statistical significance was determined by one-way ANOVA followed by Tukey’s post-hoc test for multiple comparisons among all groups. Significance is denoted as \**P* < 0.05, \*\**P* < 0.01, \*\*\**P* < 0.001, and \*\*\*\**P < 0.0001*.

**SF14.**
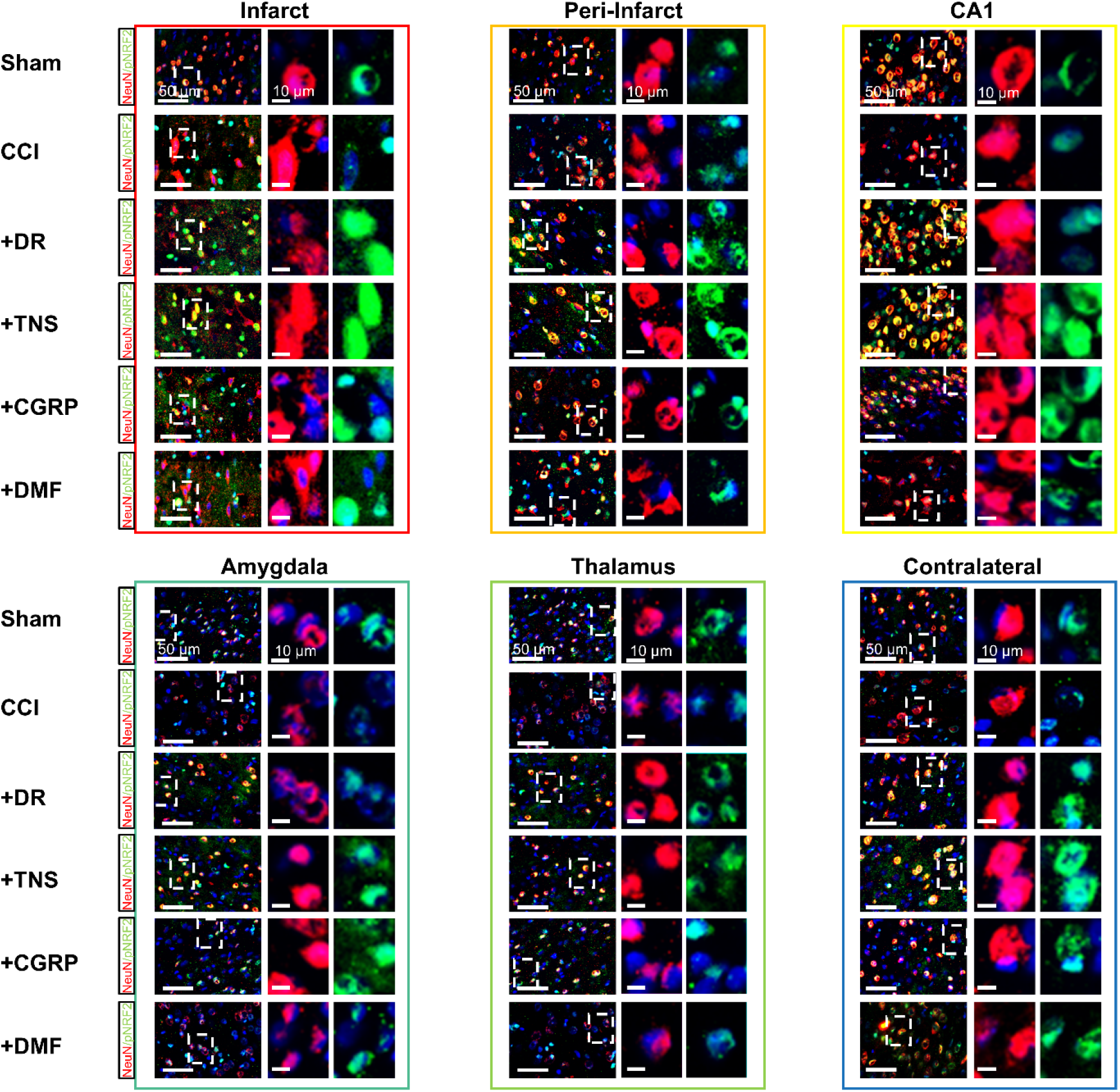
Region-specific neuronal phospho-NRF2 expression across treatment groups. Representative immunofluorescence images showing neuronal phospho-NRF2 (pNRF2; Ser40) expression at 6 hours post-CCI across six brain regions: infarct cortex, peri-infarct cortex, hippocampal CA1, amygdala, thalamus, and contralateral cortex. Images are provided for all experimental groups (sham, CCI, CCI+DR, CCI+TNS, CCI+CGRP, and CCI+DMF). The regions of interest (ROI) highlighted by the squared, dotted white boxes are shown at higher magnification in the corresponding insets.

**SF15.**
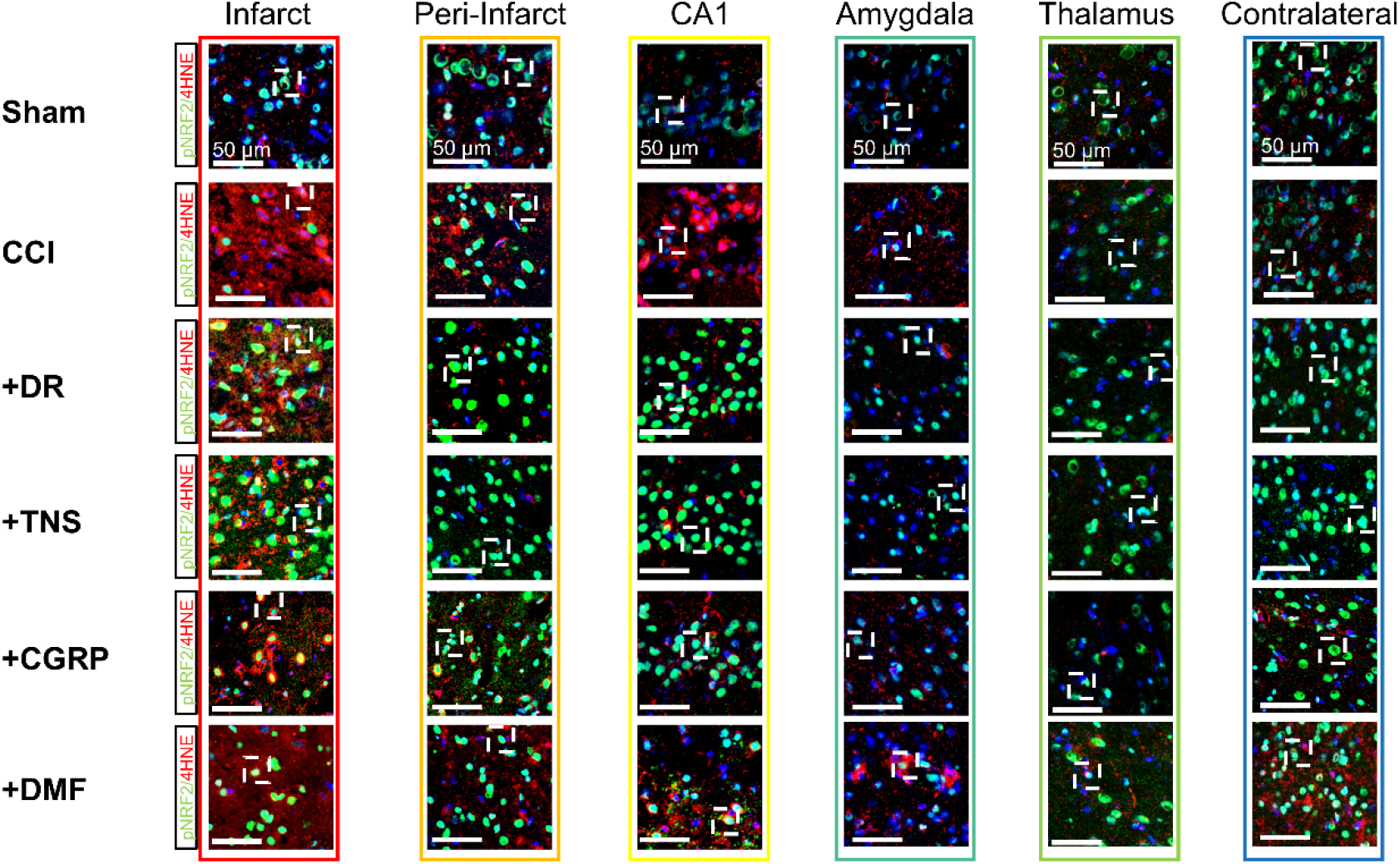
Co-localization of phospho-NRF2 and the oxidative stress marker 4HNE across brain regions. Representative high-magnification immunofluorescence images showing co-staining for phospho-NRF2 (pNRF2; Ser40) and the lipid peroxidation marker 4HNE at 6 hours post-CCI. Images are provided for all six assessed brain regions (infarct cortex, peri-infarct cortex, hippocampal CA1, amygdala, thalamus, and contralateral cortex) across all experimental groups (sham, CCI, CCI+DR, CCI+TNS, CCI+CGRP, and CCI+DMF). The regions of interest (ROI) highlighted by the squared, dotted white boxes are shown at higher magnification in the corresponding insets.

**SF16.**
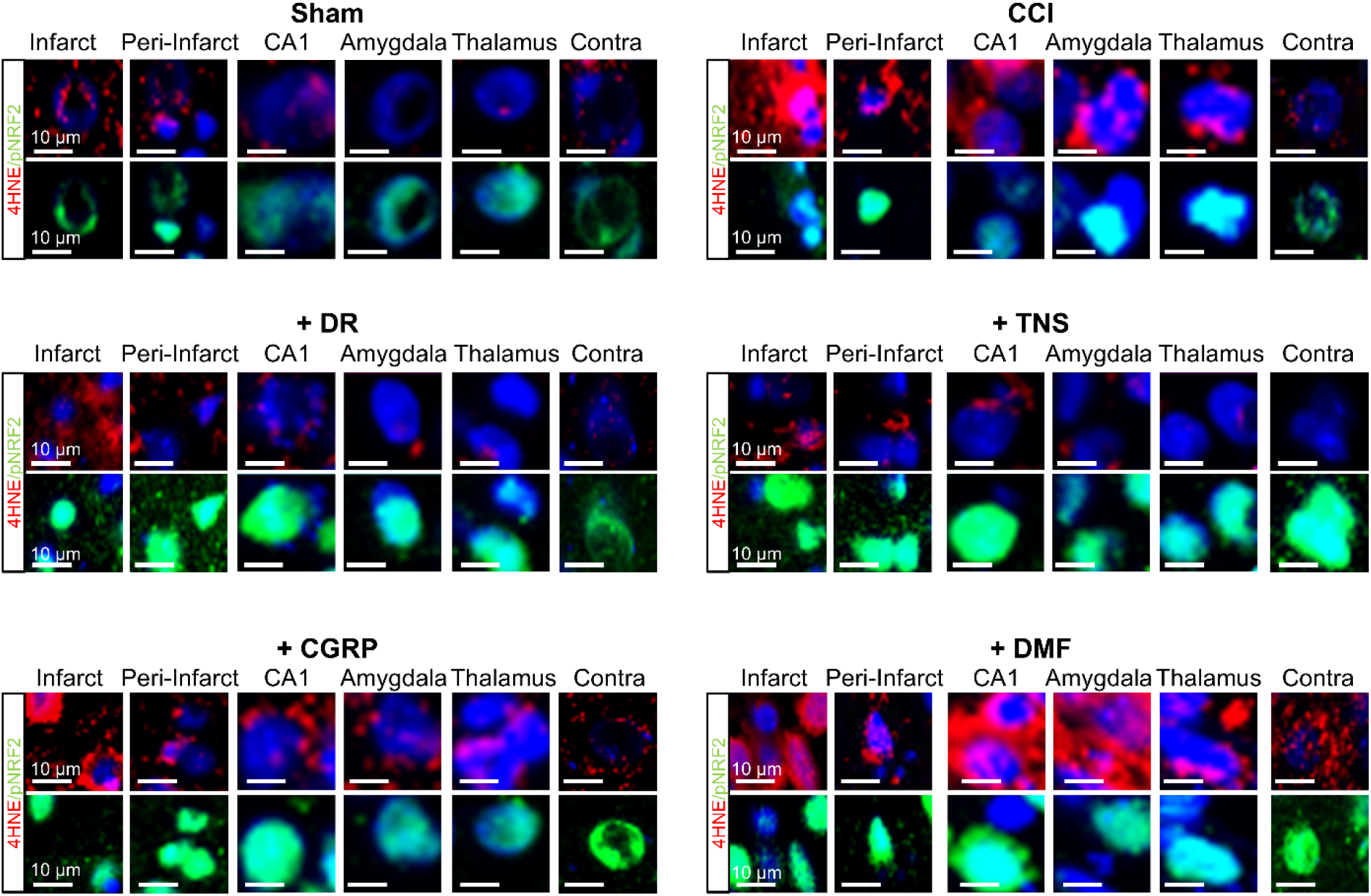
Individual channel immunofluorescence for phospho-NRF2 and 4HNE across brain regions. Representative immunofluorescence images displaying the individual staining channels for phospho-NRF2 (pNRF2; Ser40) and the lipid peroxidation marker 4HNE, corresponding to the merged co-staining images presented. Images are provided for all six assessed brain regions (infarct cortex, peri-infarct cortex, hippocampal CA1, amygdala, thalamus, and contralateral cortex) across all experimental groups (sham, CCI, CCI+DR, CCI+TNS, CCI+CGRP, and CCI+DMF). These images allow for clear, independent visualization of the expression patterns of both the NRF2 activation marker and the oxidative stress marker.

**SF17.**
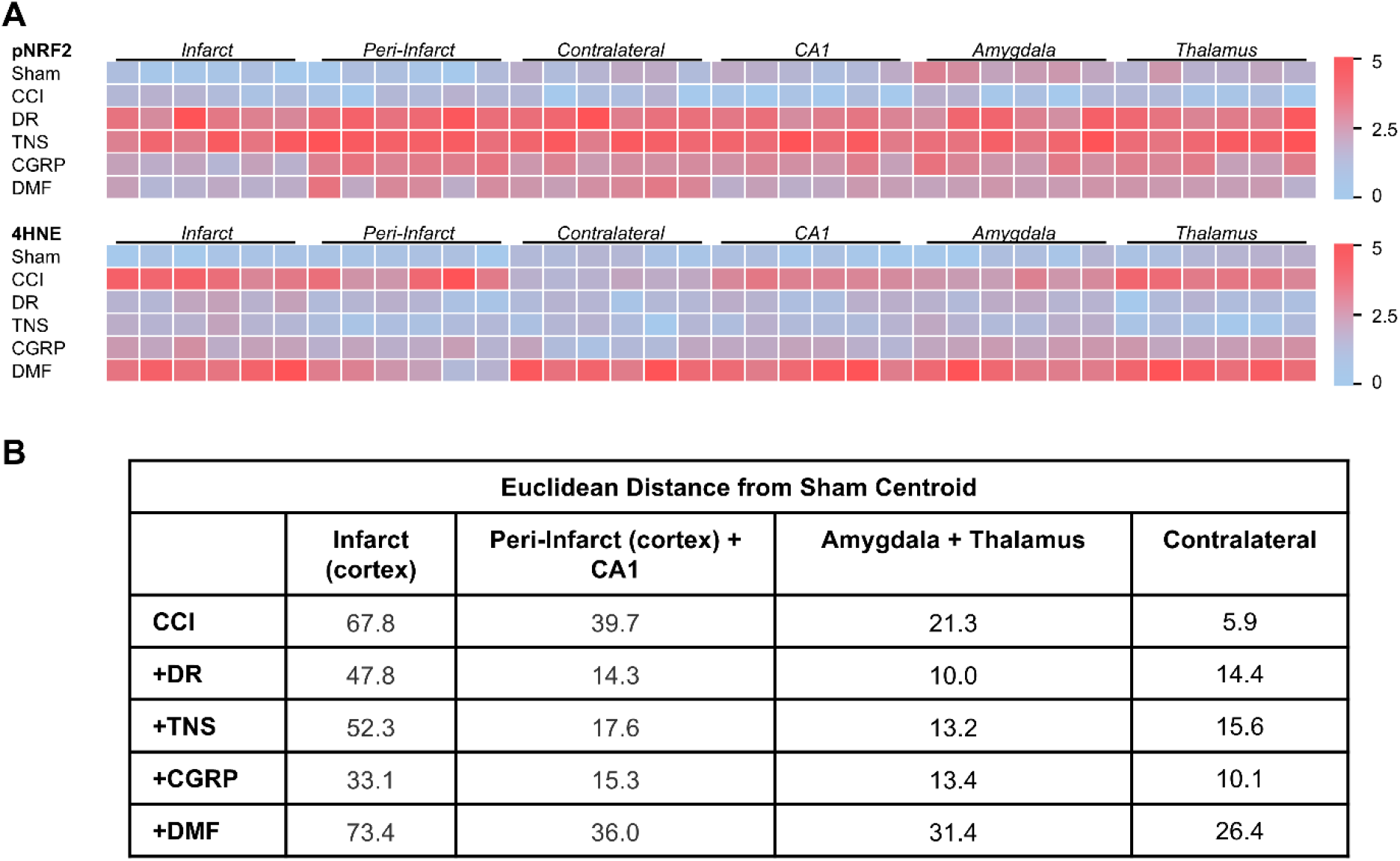
Heatmap and euclidean distance analyses quantify the divergent effects of non-electrophilic vs. electrophilic treatments. **(A)** Heatmap analysis visualizing the relative expression of phospho-NRF2 (pNRF2) and the oxidative stress marker 4HNE across all six assessed brain regions (infarct, peri-infarct, CA1, amygdala, thalamus, and contralateral cortex). The maps illustrate that DR, TNS, and CGRP treatments lead to a profile characterized by high pNRF2 and low 4-HNE, whereas the CCI and DMF groups show a pathological profile of low pNRF2 and high 4-HNE. **(B)** Corresponding table providing the exact Euclidean Distance (ED) values for each treatment group’s centroid relative to the sham centroid in the pNRF2 vs. 4-HNE feature space. The data confirm that the non-electrophilic treatment groups (DR, TNS, CGRP) cluster closely with the healthy sham state, while the DMF and untreated CCI groups remain distant, quantifying the superior restorative effect of the non-electrophilic strategies.

**SF18.**
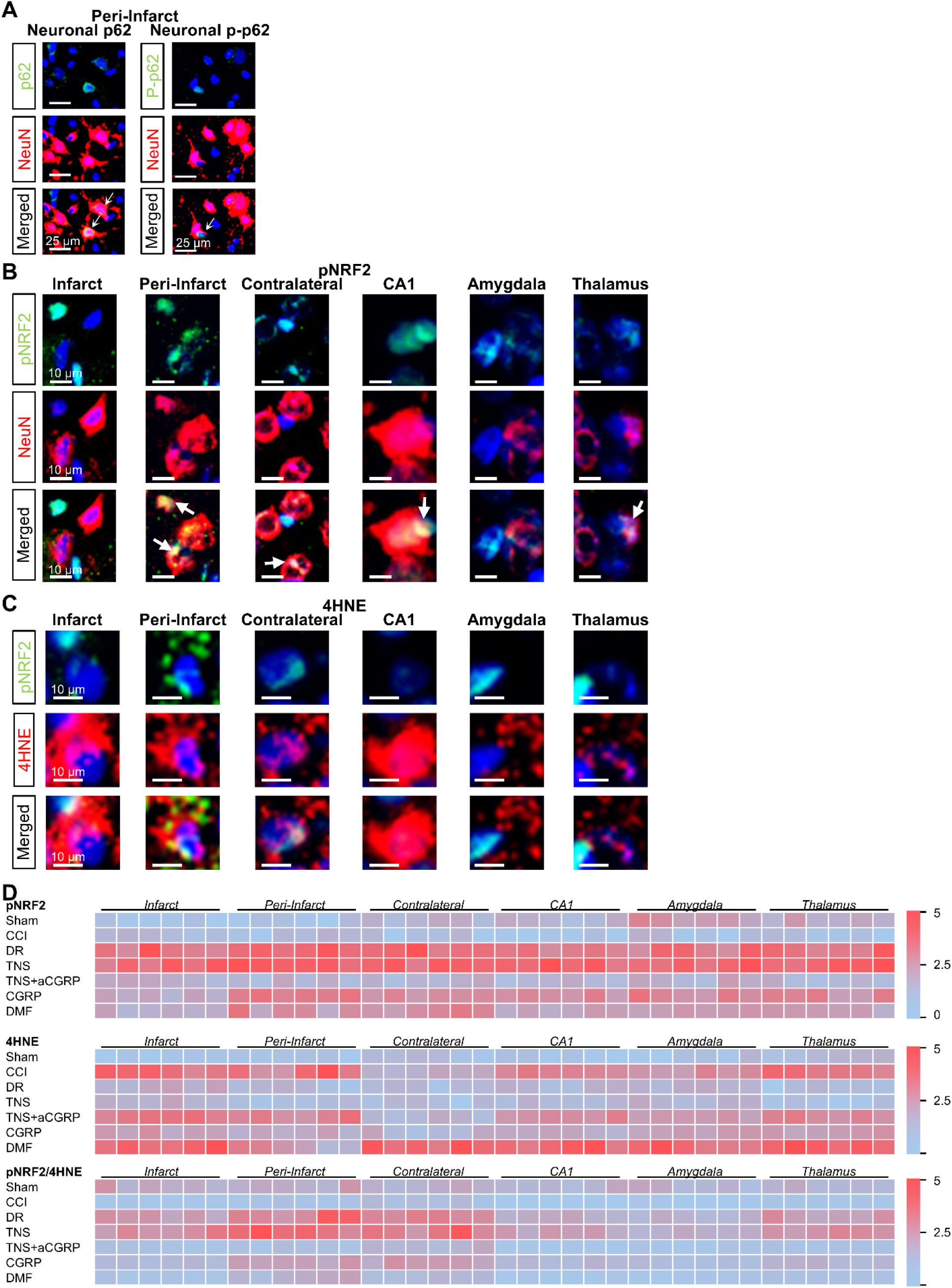
Pharmacological CGRP blockade abrogates the TNS-induced p62-NRF2 antioxidant cascade. Analysis of brain tissue from animals treated with TNS plus the CGRP receptor antagonist CGRP_8-37_ to confirm the CGRP-dependency of the protective cascade. **(A)** Representative single- and merged-channel immunofluorescence images of total p62 and phospho-p62 (Ser351) co-stained with NeuN in the peri-infarct cortex. **(B)** Representative single- and merged-channel images of neuronal phospho-NRF2 (pNRF2) co-stained with NeuN across all six assessed brain regions (infarct, peri-infarct, CA1, amygdala, thalamus, and contralateral cortex). **(C)** Representative single- and merged-channel images of phospho-NRF2 and the oxidative stress marker 4HNE across the same six brain regions. **(D)** Summary heatmap analysis of pNRF2, 4HNE, and the pNRF2-to-4HNE ratio, now including the CCI+TNS+aCGRP group for direct comparison against all other experimental groups. The heatmap visually confirms that blocking CGRP shifts the TNS-treated group’s molecular profile away from the healthy, sham-like state and back towards the pathological profile observed in untreated CCI.

**SF19.**
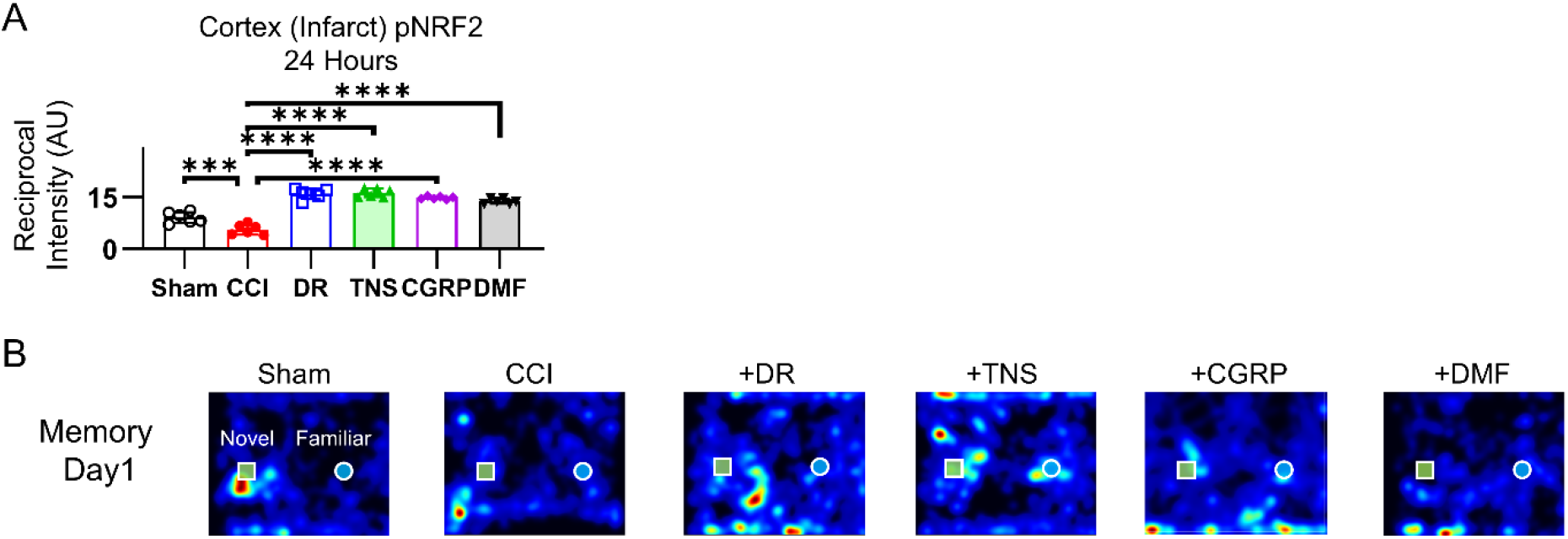
Molecular and behavioral data at 24 Hours post-CCI. **(A)** Quantification of neuronal phospho-NRF2 (pNRF2; Ser40) immunofluorescence in the infarct cortex at 24 hours post-CCI. **(B)** Representative traces from the Novel Object Recognition (NOR) test at 24 hours post-injury. The traces visually depict the exploratory paths and object interaction times for animals from each of the six experimental groups (sham, CCI, CCI+DR, CCI+TNS, CCI+CGRP, and CCI+DMF), corresponding to the short-term memory assessment. Statistical significance was determined by one-way ANOVA followed by Tukey’s post-hoc test for multiple comparisons among all groups. Significance is denoted as \**P* < 0.05, \*\**P* < 0.01, \*\*\**P* < 0.001, and \*\*\*\**P < 0.0001*.

**SF20.**
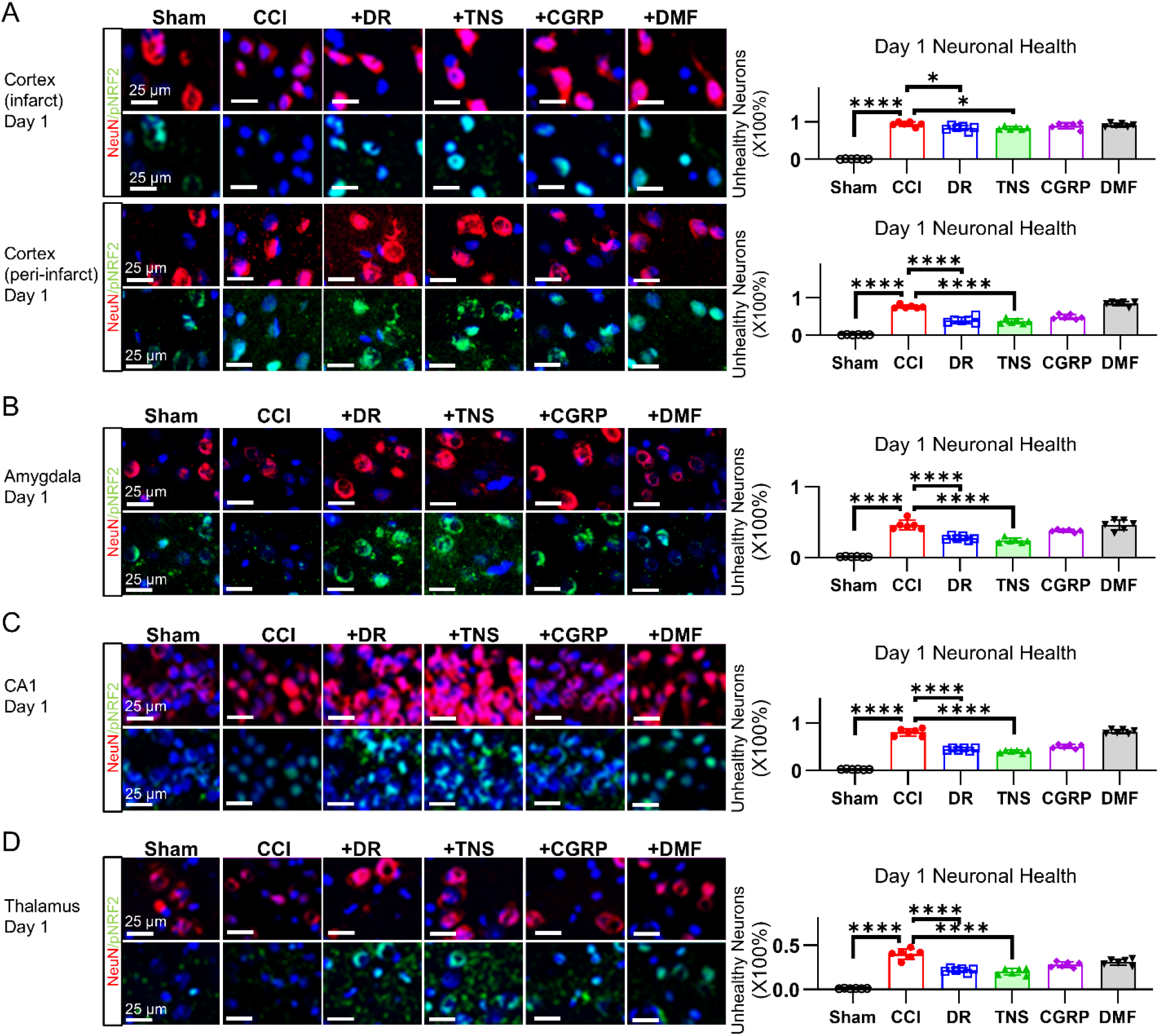
Region-specific neuronal viability and pNRF2 expression at 24 hours post-CCI. Immunofluorescence analysis of neuronal viability (NeuN) and activated phospho-NRF2 (pNRF2) at 24 hours post-CCI across all treatment groups (sham, CCI, CCI+DR, CCI+TNS, CCI+CGRP, and CCI+DMF). **(A)** Infarct cortex: Representative single-channel images for NeuN and pNRF2, with corresponding quantification of neuronal health (NeuN^+^ cells). **(B)** Amygdala. **(C)** Hippocampal CA1. **(D)** Thalamus. Statistical significance was determined by one-way ANOVA followed by Tukey’s post-hoc test for multiple comparisons among all groups. Significance is denoted as \**P* < 0.05, \*\**P* < 0.01, \*\*\**P* < 0.001, and \*\*\*\**P < 0.0001*.

**SF21.**
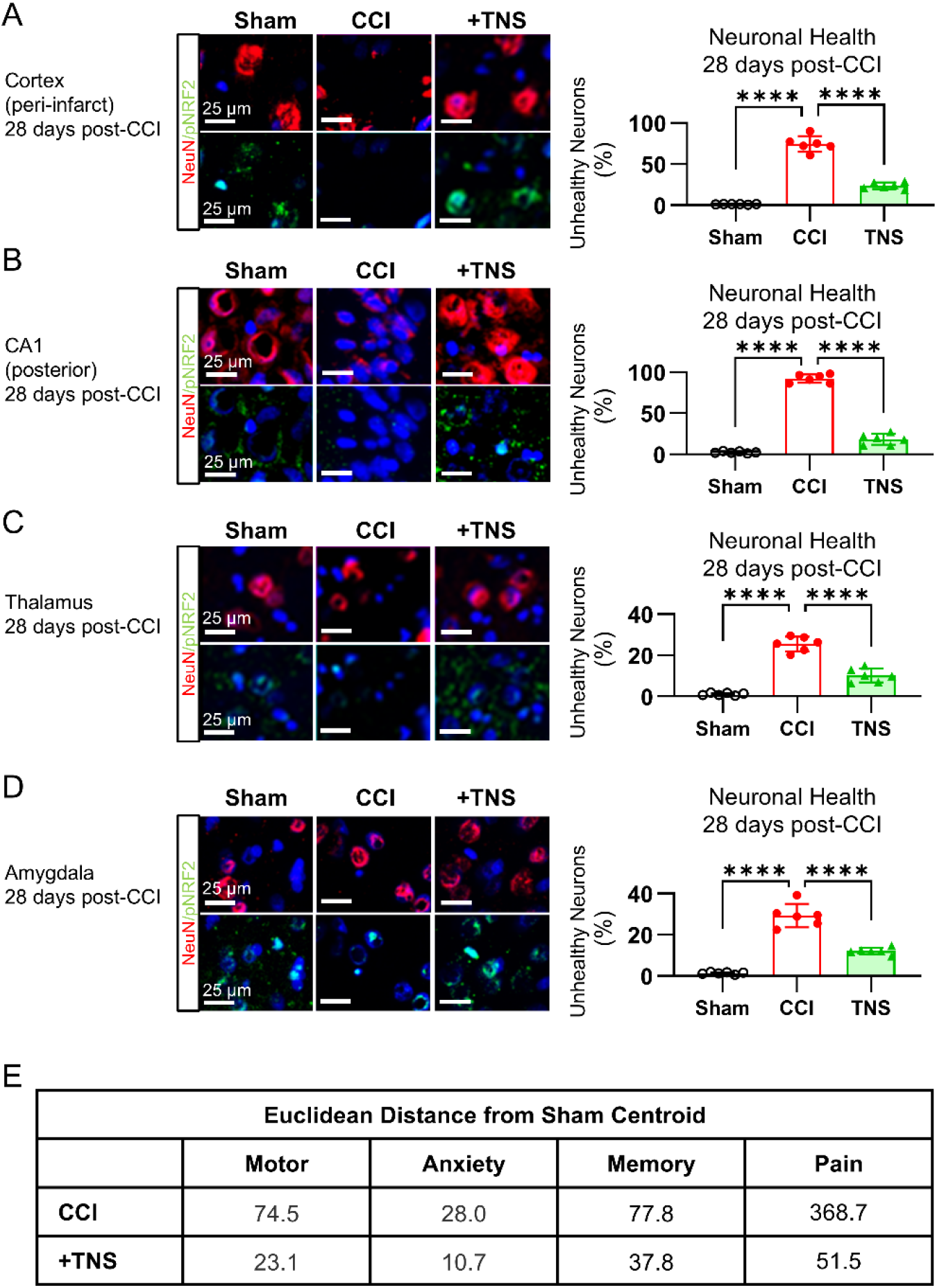
TNS confers sustained neuronal viability and pNRF2 expression at 28 Days post-CCI. Immunofluorescence analysis of long-term neuronal viability (NeuN morphology) and activated phospho-NRF2 (pNRF2) at 28 days post-CCI across the sham, CCI-vehicle, and CCI+TNS treatment groups. **(A)** Cortex: Representative single-channel images for NeuN and pNRF2, with corresponding quantification of neuronal health (NeuN^+^ cells mophology), demonstrating preserved neuronal integrity in the TNS group. **(B)** Hippocampal CA1. **(C)** Thalamus. **(D)** Amygdala. **(E)** Table providing the Euclidean Distance (ED) values for the CCI and CCI+TNS group centroids relative to the sham centroid. Statistical significance was determined by one-way ANOVA followed by Tukey’s post-hoc test for multiple comparisons among all groups. Significance is denoted as \**P* < 0.05, \*\**P* < 0.01, \*\*\**P* < 0.001, and \*\*\*\**P < 0.0001*.

**Table S1.** Product information of commercial kits.

| **Product Name** | **Vendor** | **SKU** |
|---|---|---|
| NE-PER™ Nuclear and Cytoplasmic Extraction Reagents | Thermo Scientific | 78835 |
| GSH/GSSG Ratio Detection Assay Kit (Fluorometric - Green) | Abcam | ab138881 |
| Lipid Peroxidation (MDA) Assay Kit (Colorimetric/Fluorometric) | Abcam | ab118970 |

**Table S2.** The antibodies used in this study.

| Antibodies | Source | SKU | Host | Application and dilution |
|---|---|---|---|---|
| CGRP | Santa Cruz Biotechnology | sc-57053 | ms | Western blotting, 1:100 in 5% milk in TBST |
| CGRP | Abcam | ab81887 | ms | Immunofluorescence, 1:50 in 1% BSA in TBST |
| NRF2 | Proteintech | 16396-1-AP | rb | Western blotting, 1:1000 in 5% BSA in TBST |
| Phosphorylated NRF2, s40 | Abcam | AB76026 | rb | Immunofluorescence, 1:500 in 1% BSA in TBST |
| p62 | Abcam | Ab109012 | rb | Western blotting, 1:1000 in 5% BSA in TBST<br>Immunofluorescence, 1:500 in 1% BSA in TBST |
| Phosphorylated p62, ser349 | Proteintech | 29503-1-AP | rb | Western blotting, 1:1000 in 5% milk in TBST<br>Immunofluorescence, 1:500 in 1% BSA in TBST |
| Phosphorylated p62, ser403 | Cell Signaling Technology | 39786 | Rb | Western blotting, 1:1000 in 5% milk in TBST |
| LC3 | Proteintech | 14600-1-AP | Rb | Western blotting, 1:2000 in 5% milk in TBST |
| LC3II | Proteintech | 18725-1-AP | Rb | Western blotting, 1:1000 in 5% milk in TBST |
| KEAP1 | Proteintech | 10503-2-AP | rb | Western blotting, 1:1000 in 5% BSA in TBST |
| HO1 | Proteintech | 10701-1-AP | rb | Western blotting, 1:1000 in 5% milk in TBST |
| SOD1 | Proteintech | 10269-1-AP | rb | Western blotting, 1:2500 in 5% milk in TBST |
| Nitrotyrosine | Santa Cruz Biotechnology | Sc-101358 | ms | Immunofluorescence, 1:100 in 1% BSA in TBST |
| 4-HNE | Abcam | ab48506 | N/A | Immunofluorescence, 1:25 in 1% BSA in TBST |
| NeuN | Abcam | Ab104224 |  | Immunofluorescence, 1:500 in 1% BSA in TBST |
| β-actin | Sigma | A5441-.2ML | ms | Western blotting, 1:25000 in 5% milk in TBST |
| Lamin B1 | Proteintech | 66095-1-IG | ms | Western blotting, 1:5000 in 5% milk in TBST |
| DAPI Solution | Thermo Fisher Scientific | 62248 | N/A | Immunofluorescence, 1:2000 |

